# The transcription factor EGR2 is indispensable for tissue-specific imprinting of alveolar macrophages in health and tissue repair

**DOI:** 10.1101/2021.05.06.442095

**Authors:** Jack McCowan, Phoebe M. Kirkwood, Frédéric Fercoq, Wouter T’Jonck, Connar M. Mawer, Richard Cunningham, Ananda S. Mirchandani, Anna Hoy, Gareth-Rhys Jones, Carsten G. Hansen, Nik Hirani, Stephen J. Jenkins, Sandrine Henri, Bernard Malissen, Sarah R. Walmsley, David H. Dockrell, Philippa T. K. Saunders, Leo M. Carlin, Calum C. Bain

## Abstract

Alveolar macrophages are the most abundant macrophages in the healthy lung where they play key roles in homeostasis and immune surveillance against air-borne pathogens. Tissue-specific differentiation and survival of alveolar macrophages relies on niche-derived factors, such as colony stimulating factor 2 (CSF-2) and transforming growth factor beta (TGF-β). However, the nature of the downstream molecular pathways that regulate the identity and function of alveolar macrophages and their response to injury remains poorly understood. Here, we identify that the transcriptional factor EGR2 is an evolutionarily conserved feature of lung alveolar macrophages and show that cell-intrinsic EGR2 is indispensable for the tissue-specific identity of alveolar macrophages. Mechanistically, we show that EGR2 is driven by TGF-β and CSF-2 in a PPAR-γ-dependent manner to control alveolar macrophage differentiation. Functionally, EGR2 was dispensable for lipid handling, but crucial for the effective elimination of the respiratory pathogen *Streptococcus pneumoniae*. Finally, we show that EGR2 is required for repopulation of the alveolar niche following sterile, bleomycin-induced lung injury and demonstrate that EGR2-dependent, monocyte-derived alveolar macrophages are vital for effective tissue repair following injury. Collectively, we demonstrate that EGR2 is an indispensable component of the transcriptional network controlling the identity and function of alveolar macrophages in health and disease.

**One Sentence Summary:** EGR2 controls alveolar macrophage function in health and disease

## Introduction

Tissue resident macrophages play fundamental roles in protective immunity and wound repair following injury, but also in the maintenance of homeostasis. The functions of macrophages vary to meet the demands of the local environment, and this is reflected in the phenotypic diversity detected amongst macrophages in different tissues. Indeed, although all tissue macrophages possess a common ‘core’ transcriptional signature *(1)*, there are additional tissue-specific gene expression characteristics that enable organ- and even niche-specific functions *(2, 3)*. However, the local environmental signals and the downstream molecular pathways that control this tissue-specific imprinting of macrophages in different environments are incompletely understood.

Alveolar macrophages are the most abundant macrophage population in the healthy lung, where they provide a first line of defence against airborne pathogens, as well as maintaining lung homeostasis, for instance through the regulation of pulmonary surfactant. However, in chronic lung pathologies such as allergic asthma, idiopathic pulmonary fibrosis (IPF) and chronic obstructive pulmonary disease (COPD), alveolar macrophages display aberrant activity and, in many cases, appear to perpetuate disease *(4)*. Moreover, monocytes and macrophages appear to play a particular pathogenic role in the context of severe coronavirus disease 2019 (COVID-19) *(5)*. Thus, the mechanisms governing alveolar macrophage imprinting may yield important insights into how lung-specific cues regulate homeostasis and susceptibility to disease.

Alveolar macrophages are derived from erythromyeloid progenitors (EMPs) and foetal liver monocytes that seed the lung during embryonic development *(6–8)*. However, the characteristic phenotype and functional properties of alveolar macrophages do not develop until the first few days of postnatal life in parallel with alveolarisation of the lung and are controlled by CSF-2 (GM-CSF) *(8–10)* and the immunoregulatory cytokine TGF-β *(11)*. Together these cytokines induce expression of the transcription factor peroxisome proliferator-activated receptor gamma (PPAR-γ) to promote survival and tissue-specific specialisation, including upregulation of genes involved in lipid uptake and metabolism. Consequently, mice in whom *Csf2rb, Tgfbr2* or *Pparg* has been genetically ablated in myeloid cells develop spontaneous pulmonary alveolar proteinosis *(9–11)*. However, alveolar macrophages largely fail to develop in the absence of CSF-2 and TGF-β receptor signalling due to their key role in macrophage survival. Therefore, it remains unclear if or how these factors control the tissue-specific identity and function of alveolar macrophages. Moreover, while considered the ‘master transcription factor’ of alveolar macrophages, PPAR-γ has been implicated in the control of other tissue macrophages, including splenic red pulp macrophages *(12, 13)*, and thus, the transcriptional network responsible for conferring specificity upon alveolar macrophage differentiation remains unclear. Finally, if and how additional transcriptional regulators are involved in regulating these processes in the context of inflammation and repair is largely unexplored.

Here, we have used single cell RNA sequencing (scRNA-seq) to identify the transcriptional regulators expressed by alveolar macrophages. We show that expression of the transcriptional factor EGR2 is a unique feature of lung alveolar macrophages compared with other lung mononuclear phagocytes and macrophages resident in other tissues. Using cell-specific ablation of *Egr2* and mixed bone marrow chimeric mice, we show that cell-intrinsic EGR2 is indispensable for the tissue-specific identity of alveolar macrophages and their ability to control infection with a major respiratory pathogen, *Streptococcus pneumoniae*. RNA sequencing (RNA-seq) shows that EGR2 controls a large proportion of the core transcriptional signature of alveolar macrophages, including expression of *Siglec5, Epcam* and *Car4*. Mechanistically, we show that EGR2 expression is induced by TGF-β and CSF-2-dependent signalling, and acts to maintain expression of CCAAT-enhancer-binding protein beta (C/EBPβ) to control alveolar macrophage differentiation. Finally, using the bleomycin-induced model of lung injury and a combination of fate mapping approaches, we show that post-injury repopulation of the alveolar macrophage niche occurs via differentiation of bone marrow-derived cells in an EGR2-dependent manner and that these monocyte-derived macrophages are indispensable for effective tissue repair and resetting of tissue homeostasis.

## Results

### EGR2 expression is a selective property of alveolar macrophages

To begin to dissect the molecular pathways underlying the niche-specific imprinting of alveolar macrophages, we performed scRNA-seq of murine lung mononuclear phagocytes from lung digests to identify the unique transcriptional profile of alveolar macrophages. To this end, non-granulocytic CD45^+^ cells from lungs of *Rag1*^−/−^ mice were purified by FACS and sequenced using the 10x Chromium platform (**Supplementary Figure 1A**). 3936 cells passed quality control and were clustered using Uniform Manifold Approximation and Projection (UMAP) dimensionality reduction analysis within the *Seurat* R package. NK cells, identified by their expression of *Ncr1, Nkg7* and *Gzma*, were excluded (**Supplementary Figure 1A**) and the remaining myeloid cells were re-clustered to leave six clusters of mononuclear phagocytes, and these were annotated using known landmark gene expression profiles (**Figure 1A, B**). Cluster 1 represented monocytes based on their expression of *Itgam* (encoding CD11b)*, Csf1r* and *Cd68*, and could be divided into classical and non-classical monocytes based on expression of *Ly6c2* and *Treml4* respectively (**Figure 1A, B**). Cluster 2 represented interstitial macrophages based on their high expression of *Cx3cr1*, *Cd68, Csf1r* and *H2-Aa* and lack of the *Xcr1* and *Cd209a* genes which defined cDC1 (cluster 5) and cDC2 (cluster 6) respectively. Alveolar macrophages (cluster 3) formed the largest population and could be defined by their expression of *Itgax* (encoding CD11c)*, Siglec5* (encoding SiglecF) and *Car4*, and lack of *Cx3cr1* and *Itgam*. Cluster 4 was transcriptionally similar to cluster 3, but was defined by genes associated with cell cycle, including *Mki67, Birc5* and *Tubb5*, suggesting these represent proliferating alveolar macrophages (**Figure 1A, B**). Next, we compared gene expression profiles of these clusters, focussing on genes more highly expressed by alveolar macrophages relative to all other mononuclear phagocytes. 722 genes fitted these criteria, including *Fapb1*, *Spp1* (encoding osteopontin) and *Cidec* which are known to be uniquely and highly expressed by alveolar macrophages (**Supplementary Table 1**) *(1, 3)*. Within this cassette of genes, we turned our attention to genes encoding transcription factors/regulators, as we hypothesised that these might control the tissue specific differentiation of alveolar macrophages. As expected, these included *Pparg, Cebpb* and *Bhlhe41* which have been shown to control the development and self-renewal capacity of alveolar macrophages *(9, 14, 15)* (**Figure 1C**). However, this analysis also revealed transcription factors such as *Id1, Klf7* and *Egr2* which have not previously been implicated in the control of alveolar macrophage differentiation. We focussed on EGR2, which is part of a family of early growth response (EGR) transcription factors, comprising EGR1-4, as *Egr2* appeared to be expressed in a particularly selective manner by alveolar macrophages (**Figure 1D**) when compared with other tissue macrophages at mRNA (**Figure 1E**) and protein level (**Figure 1F, G & Supplementary Figure 2A**). In contrast, while highly expressed by alveolar macrophages, *Pparg* was also expressed at a high level by splenic red pulp macrophages (**Figure 1E**), consistent with previous reports *(12, 13)*. Next, we performed analogous analysis of *EGR2* expression across a variety of human macrophage populations from scRNA-seq data sets within the Human Cell Atlas. Consistent with our analysis in the mouse, this showed that *EGR2* expression was confined to lung macrophages, and in particular *FABP4*^+^ macrophages which correspond to airway macrophages (**Figure 1H**), and we confirmed this at protein level, showing that CD163^+^HLA-DR^+^ bronchoalveolar lavage (BAL) macrophages uniformly express EGR2 (**Supplementary Figure 2B**). Thus, these data demonstrate that EGR2 expression is a constitutive, specific and evolutionary conserved feature of alveolar macrophages.

**Figure 1.**
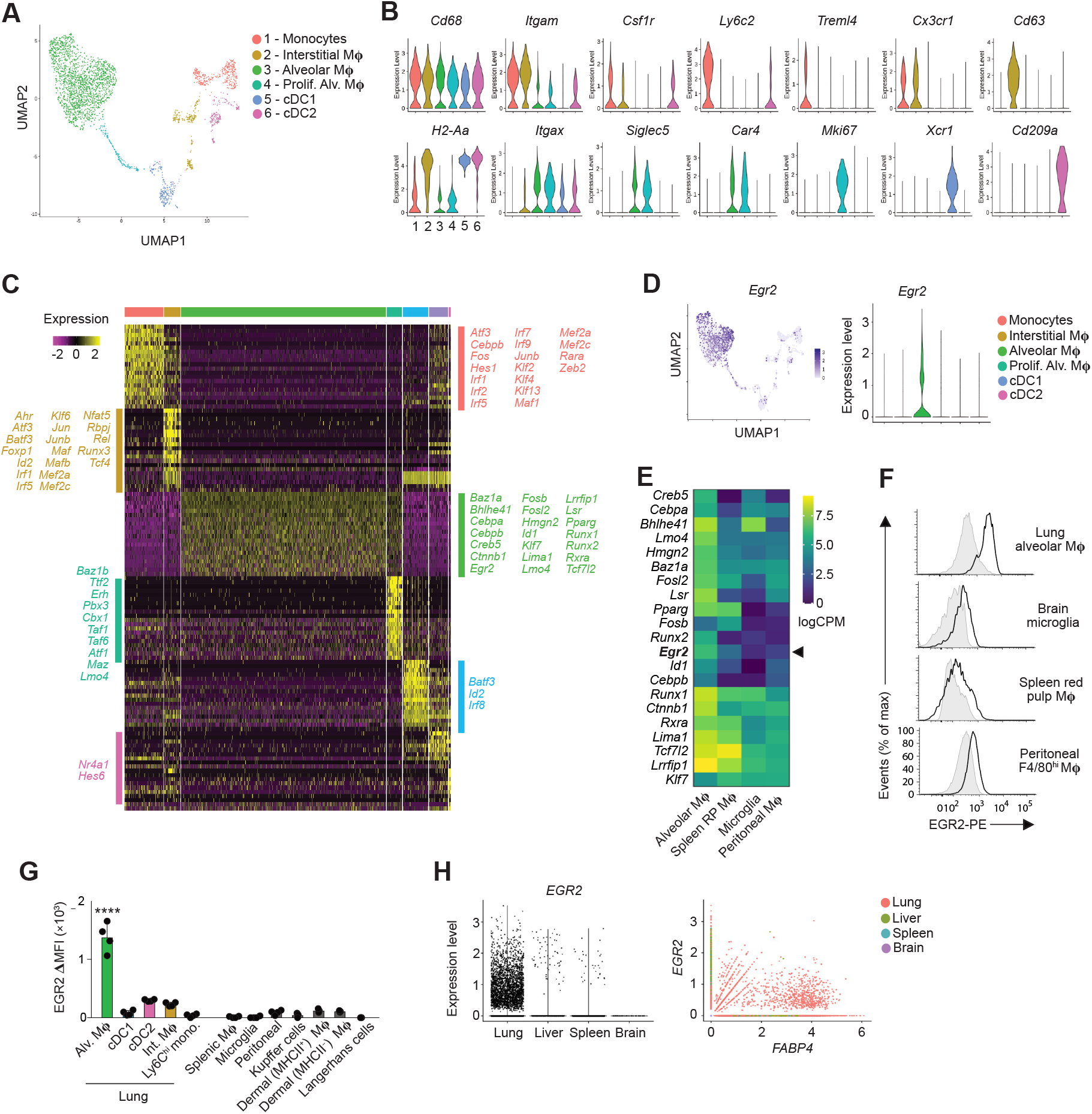
EGR2 expression is a selective property of alveolar macrophages. **A.** UMAP dimensionality reduction analysis of 3936 cells (non-granulocyte, myeloid cells) reveals six clusters of mononuclear phagocytes, including monocytes, macrophages (Mϕ) and conventional dendritic cells (cDC) in murine lungs. **B.** Feature plots displaying expression of individual genes by clusters identified in **A.** **C.** Heatmap showing the top 20 most differentially expressed genes by each cluster defined in **A**. and annotated to show the upregulated transcription factors/regulators within each cluster. **D.** Overlay UMAP plot and feature plot showing expression of *Egr2* by clusters identified in **A.** **E.** Heatmap showing relative expression of selected transcription factors by lung alveolar macrophages, CD102^+^ peritoneal macrophages, brain microglia and red pulp splenic macrophages as derived from the ImmGen consortium. **F.** Representative expression of EGR2 by lung alveolar macrophages, CD102^+^ peritoneal macrophages, brain microglia and red pulp splenic macrophages obtained from adult unmanipulated C57BL/6 mice. Shaded histograms represent isotype controls. Data are from one of three independent experiments with 3-4 mice per experiment. **G.** Expression of EGR2 by the indicated macrophage and myeloid cell populations shown as relative mean fluorescence intensity (MFI) (MFI in *Egr2*^fl/fl^ (Cre^−^) – MFI in *Lyz2*^Cre/+^.*Egr2*^fl/fl^ (Cre^+^) mice). Data represent 3-4 mice per tissue. Error bars represent S.D. One-way ANOVA followed by Tukey’s multiple comparisons post-test. **** p<0.0001 **H.** *In silico* analysis of *EGR2* and *FABP4* expression by lung, liver, spleen and brain macrophages extracted on the basis of *C1QA*^+^ expression from *(16–18).*

### EGR2 is required for the phenotypic identity of alveolar macrophages

Previous work has suggested that EGR1 and EGR2 act in a redundant manner *(19)*, while other studies have suggested EGR transcription factors are completely dispensable for macrophage differentiation *(20)*. However, many of these studies were performed *in vitro* and the roles of EGRs in tissue-specific macrophage differentiation has not been assessed comprehensively *in vivo*, in part, due to the postnatal lethality of global *Egr2^−/−^* mice *(21, 22)*. To determine the role of EGR2 in alveolar macrophage development and differentiation, we crossed *Lyz2*^Cre^ mice *(23)* with *Egr2*^fl/fl^ mice *(24)*, to generate a strain in which myeloid cells, including monocytes, macrophages, dendritic cells and neutrophils, lack EGR2 in a constitutive manner. We performed unbiased UMAP flow cytometry analysis on lung leukocytes obtained from *Lyz2*^Cre^.*Egr2*^fl/fl^ mice (referred to here as Cre^+^) and *Egr2*^fl/fl^ littermate controls (referred to here as Cre^−^ mice), focussing on ‘lineage’ negative (CD3^−^CD19^−^NK1.1^−^ Ly6G^−^) CD11c^+^ and CD11b^+^ cells in lung digests (**Figure 2A**). Surface marker analysis of cells pooled from Cre^−^ and Cre^+^ mice confirmed the presence of alveolar and interstitial macrophages, eosinophils and subsets of dendritic cells and monocytes (**Figure 2A**) and this was validated by manual gating (**Figure 2B** & **Supplementary Figure 3A**). Due to their CD11c^hi^CD11b^−^ phenotype, alveolar macrophages clustered separately from the other CD11b^+^ myeloid cells (**Figure 2A-C**). All myeloid cells, including alveolar macrophages, were equally abundant in the lungs of Cre^−^ and Cre^+^ mice. (**Figure 2D**). However, whereas alveolar macrophages from Cre^−^ mice expressed high levels of SiglecF, the majority of alveolar macrophages obtained from Cre^+^ mice lacked SiglecF expression (**Figure 2E**), explaining their distinct positioning within the alveolar macrophage cluster in the UMAP analysis. Indeed only ~7% of alveolar macrophages in Cre^+^ mice expressed high levels of SiglecF, and further analysis showed that these expressed high levels of EGR2 (**Figure 2F**), suggesting that the SiglecF^+^ cells remaining in the Cre^+^ mouse represent cells that have escaped Cre-mediated recombination. Consistent with this, SiglecF^+^ cells in the Cre^+^ mouse expressed high levels of CD11c equivalent to alveolar macrophages from Cre^−^ mice, whereas SiglecF^−^ alveolar macrophages expressed lower levels of CD11c (**Figure 2G**). We did not detect differences in the proliferative activity of *Egr2-*sufficient and - deficient alveolar macrophages (**Figure 2G**). Importantly and consistent with the lack of EGR2 expression by other tissue resident macrophages, we saw no effect on the cell number and expression of signature markers by resident macrophages in other tissues, including in the spleen where macrophages share a dependence on PPAR-γ *(12, 13)* (**Supplementary Figure 3B, C**). Thus, these data demonstrate that while EGR2 expression is dispensable for alveolar macrophages survival and self-maintenance, it is indispensable for imprinting key phenotypic features of the cells in the healthy lung.

**Figure 2:**
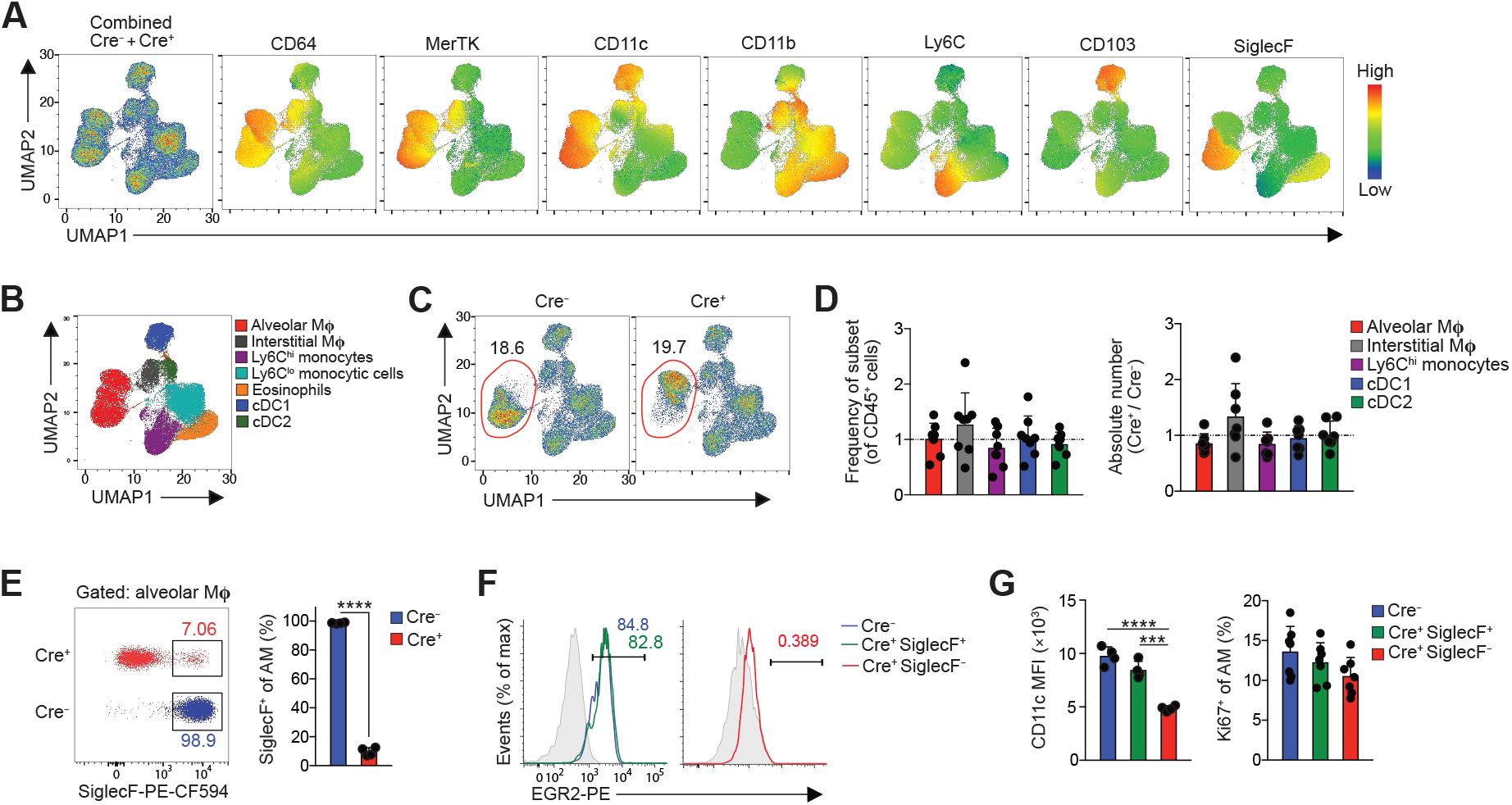
EGR2 is required for the phenotypic identity of alveolar macrophages. **A.** UMAP analysis of CD3^−^CD19^−^NK1.1^−^Ly6G^−^CD11b^+^/CD11c^+^ cells pooled from adult unmanipulated Cre^−^ and Cre^+^ mice (*left panel*). Heatmap plots showing the relative expression of the indicated markers by myeloid clusters. **B.** Cluster identity confirmed by manual gating (see **Supplementary Figure 3A**). **C.** Relative frequency of alveolar macrophages (red gate) of all CD45^+^ leukocytes in unmanipulated adult Cre^−^ and Cre^+^ mice. **D.** Relative frequency and absolute number of alveolar macrophages, cDC1, cDC2, Ly6C^hi^ monocytes and CD64^+^MHCII^+^ interstitial macrophages in lung digests from adult unmanipulated Cre^+^ mice compared with their abundance in Cre^−^ littermates. Symbols represent individual mice and data are pooled from three independent experiments with 8 mice per group. Error bars represent S.D. **E.** Representative expression of SiglecF by CD11c^hi^CD11b^lo^ alveolar macrophages (from **F**) obtained from lung digests from adult unmanipulated Cre^−^ or Cre^+^ mice (*left*), frequency of SiglecF^+^ macrophages in each strain (*right*). Symbols represent individual mice with 4 mice per group. Data are from one of at least 5 independent experiments. Error bars represent S.D. Student’s *t* test, ****p<0.0001 (SiglecF) **F.** Representative expression of EGR2 by SiglecF-defined alveolar macrophages. Shaded histograms represent isotype controls. Data are from one of three independent experiments with 3-4 mice per experiment. **G.** Mean fluorescence intensity (MFI) of CD11c expression by SiglecF-defined CD11c^hi^CD11b^lo^ alveolar macrophages from lung digests from adult unmanipulated Cre^−^ or Cre^+^ mice. Symbols represent individual mice with 4 mice per group. Data are from one of at least 5 independent experiments. Error bars represent S.D. One-way ANOVA followed by Tukey’s multiple comparisons post-test. *** p<0.001, **** p<0.0001.

### EGR2 controls the tissue-specific transcriptional programme of alveolar macrophages

The failure of alveolar macrophages from Cre^+^ mice to express SiglecF suggested that the tissue-specific differentiation programme of these cells may be altered by *Egr2* deficiency. Hence, to ascertain the global effects of *Egr2* deletion on alveolar macrophage differentiation, we next performed bulk RNA-seq of CD11c^hi^CD11b^lo^ alveolar macrophages from lung digests of Cre^−^ and Cre^+^ mice (using only SiglecF^−^ macrophages from Cre^+^ mice to exclude confounding effects of EGR2-sufficient alveolar macrophages) (**Supplementary Figure 4**). Unbiased clustering confirmed the biological replicates from each group were highly similar (**Figure 3A**) and differential gene expression (DEG) analysis revealed that 858 genes were differentially expressed by at least 2-fold (417 and 441 genes downregulated and upregulated, respectively) (**Supplementary Table 2**). Consistent with our flow cytometry analysis, *Siglec5*, which encodes SiglecF, was one of the most downregulated genes in *Egr2* deficient alveolar macrophages (**Figure 3B**). Many of the most differentially expressed genes formed part of the alveolar macrophage gene set identified in our scRNA-seq analysis. Moreover, approximately 30% of the core alveolar macrophage signature identified by the ImmGen consortium *(1)* was altered by *Egr2* deficiency (32 genes) (**Figure 3B, C**), including the expression of *Spp1, Epcam, Car4* and *Fabp1*, all of which were confirmed by flow cytometry or qPCR (**Figure 3D, E**). The vast majority of these ‘signature’ genes was downregulated in *Egr2*-deficient macrophages compared with their *Egr2*-sufficient counterparts. Gene Ontology (GO) analysis revealed that the top pathways affected by *Egr2* deficiency were ‘Chemotaxis’, ‘Cell chemotaxis’ and ‘Immune system process’ (**Supplementary Table 3**). Consistent with this, the expression of chemokine receptors, such as *Ccr2* and *Cx3cr1*, was elevated in alveolar macrophages from Cre^+^ mice compared with their Cre^−^ counterparts (**Figure 3F**). Genes encoding antigen presentation machinery, such as *H2-Aa, H2-Eb1, Ciita* and *Cd74* were also upregulated in alveolar macrophages from Cre^+^ mice. In parallel, there was significantly greater expression of MHCII at the protein level in *Egr2* deficient alveolar macrophages (**Figure 3F**). Indeed, over 50 genes upregulated in *Egr2* deficient alveolar macrophages were genes that defined interstitial macrophages in our scRNA-seq analysis, including *Cd63, Mafb, Mmp12* and *Msr1* (**Figure 3D**, **Supplementary Table 2**). Thus, EGR2 ablation renders alveolar macrophages transcriptionally more similar to their interstitial counterparts.

**Figure 3:**
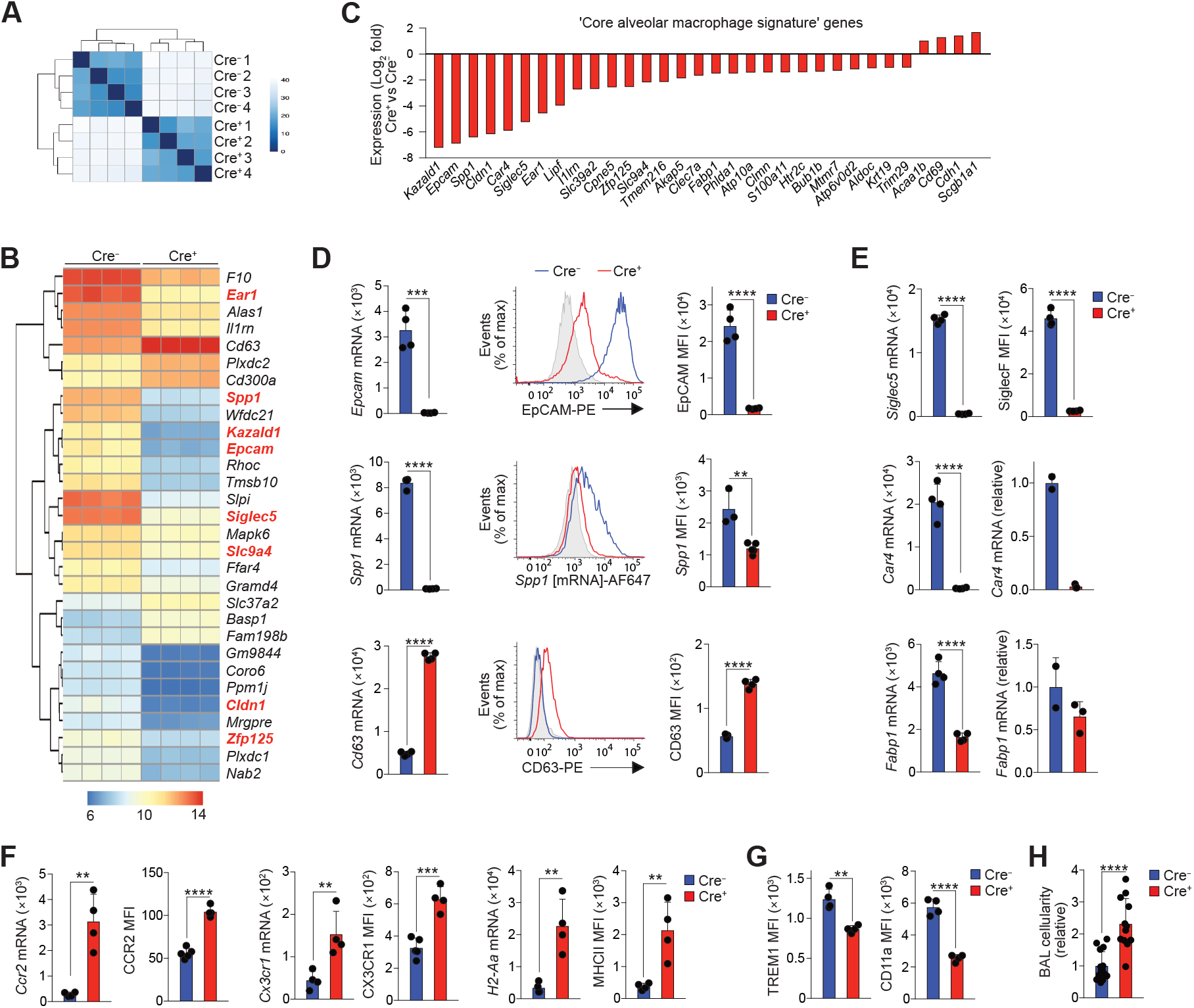
EGR2 controls tissue-specific transcriptional programme of alveolar macrophages. **A.** Heatmap of RNA-seq data showing the distance between samples from Cre^−^ and Cre^+^ mice. **B.** Heatmap showing expression of the 30 most differentially expressed genes by alveolar macrophages from Cre^−^ and Cre^+^ mice. Each column represents a biological replicate with four mice per group. Genes highlighted in red appear in the ‘core signature’ of alveolar macrophages as defined by the ImmGen Consortium *(1)*. **C.** Log_2_-fold expression of differentially expressed genes that form part of the ‘core signature’ of alveolar macrophages as defined by the ImmGen Consortium *(1)*. **D.** Expression of *Epcam*, *Spp1* and *Cd63* from the RNA-seq dataset (*left panels*), representative flow cytometric validation of EpCAM, *Spp1* (mRNA detected by PrimeFlow technology) and CD63 expression (*middle panels*) and replicate MFI expression data of each of these markers by alveolar macrophages from adult unmanipulated Cre^−^ and Cre^+^ mice. Symbols represent individual mice and data are from one of two independent experiments with 5 (Cre^−^) and 4 (Cre^+^) mice per group. Student’s *t* test, **p<0.01, ***p<0.001, ****p<0.0001. **E.** Expression of *Siglec5*, *Car4* and *Fabp1* from the RNA-seq dataset (*left panels*) and validation by flow cytometry (SiglecF) or qPCR (*Car4*, *Fabp1*). Symbols represent individual mice and data for SiglecF is from one of at least 10 independent experiments with 5 (Cre^−^) and 4 (Cre^+^) mice per group. Data for *Car4* and *Fabp1* represents 2 (Cre^−^) and 4 (Cre^+^) mice per group. Student’s *t* test, ****p<0.0001. **F.** Expression of *Ccr2*, *Cx3cr1* and *H2-Aa* from the RNA-seq dataset and replicate MFI expression data of CCR2, CX3CR1 and MHCII as determined by flow cytometry. Symbols represent individual mice and data are from one of two independent experiments with 5 (Cre^−^) and 4 (Cre^+^) mice per group. Student’s *t* test, **p<0.01, ***p<0.001, ****p<0.0001. **G.** Replicate MFI data of for CD11a and TREM1 expression as determined by flow cytometry. Symbols represent individual mice and data are from one of two independent experiments with 5 (Cre^−^) and 4 (Cre^+^) mice per group. Student’s *t* test, **p<0.01, ****p<0.0001. **H.** Absolute number of CD11c^hi^CD11b^−^ alveolar macrophages present in the BAL of adult unmanipulated Cre^+^ mice relative to their abundance in Cre^−^ littermates. Symbols represent individual mice and data are pooled from three independent experiments with 15 (Cre^−^) and 12 (Cre^+^) mice per group. In all graphs error bars represent S.D.

Further phenotypic analysis revealed reduced expression of ‘core signature’ alveolar macrophage markers TREM1 and CD11a at protein level in the context of *Egr2* deficiency (**Figure 3G**). EpCAM and CD11a expression have been implicated in regulating adherence to and patrolling of the lung epithelium by alveolar macrophages *(25)*. Interestingly, while we found equivalent numbers of alveolar macrophages amongst tissue digests, we obtained consistently higher numbers of alveolar macrophages in the bronchoalveolar lavage (BAL) fluid of Cre^+^ mice, suggesting the EGR2-dependent differentiation programme may control the ability of alveolar macrophages to adhere to and interact with cells of their niche in the airways (**Figure 3H**).

### EGR2 controls distinct functional characteristics of alveolar macrophages

Individuals with mutations in *EGR2* develop peripheral neuropathies due to the crucial role for EGR2 in Schwann cell function *(26)*. However, many of these individuals also frequently encounter respiratory complications, including recurrent pneumonias and/or restrictive pulmonary disease, and in some cases respiratory failure *(26)*. The cause of respiratory compromise in these individuals remains unexplained. To determine if alterations in alveolar macrophage behaviour may contribute to this, we next tested the function of *Egr2*-deficient alveolar macrophages. A major homeostatic function of alveolar macrophages is the regulation of pulmonary surfactant, and the absence of alveolar macrophages results in the development of pulmonary alveolar proteinosis (PAP) *(8–10, 27, 28)*. However, *Egr2* deficiency did not lead to spontaneous PAP, as there were no differences in the levels of total protein in BAL fluid from Cre^+^ and Cre^−^ mice at either 4 or >9 months of age, a time at which PAP is detectable in *Csf2rb*^−/−^ mice *(28)* (**Figure 4A**). However, these results were confounded by the fact that the majority of alveolar macrophages in aged (>9 months) Cre^+^ mice was now EGR2-sufficient, with most cells expressing high levels of SiglecF (**Figure 4B, C**). These findings suggested that the cells that had escaped Cre recombination may have a competitive advantage over their EGR2-deficient counterparts and indeed, the absolute number of SiglecF^+^ alveolar macrophages no longer differed between aged Cre^−^ and Cre^+^ mice (**Figure 4D**). These data are consistent with other studies noting age-related repopulation of the alveolar niche with Cre ‘escapees’ in the *Lyz2*^Cre^ mouse *(11)*. Notably, however, this preferential expansion of EGR2-sufficient ‘escapees’ did not relate to differences in the level of proliferation by EGR2-defined subsets, with identical frequencies of Ki67^+^ cells amongst EGR2-sufficient and - deficient macrophages in young adult and aged mice (**Figure 4E**).

**Figure 4:**
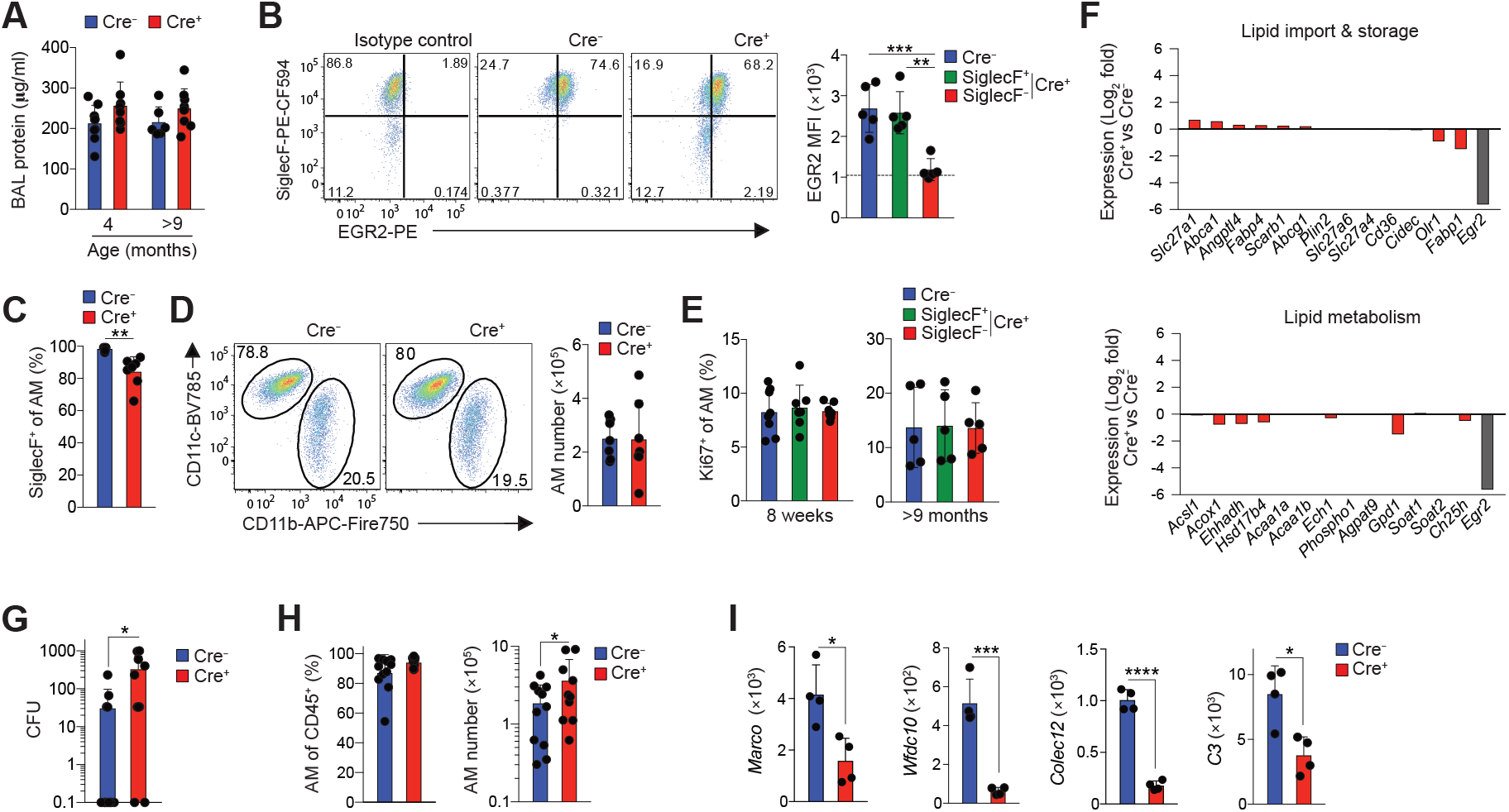
EGR2 controls distinct functional characteristics of alveolar macrophages. **A.** Protein levels in the BAL fluid of Cre^−^ or Cre^+^ mice at 4 or 9-12 months of age. Symbols represent individual mice and error is S.D. Data are from 6-9 mice per group pooled from two independent cohorts of aged mice. **B.** Representative expression of SiglecF and EGR2 by CD11c^hi^CD11b^lo^ macrophages and mean fluorescent intensity (MFI) of EGR2 by SiglecF-defined CD11c^hi^CD11b^lo^ macrophages obtained from 11-12 month old Cre^−^ or Cre^+^ mice. Symbols represent individual mice and error is S.D. Data are from 5 mice per group pooled from two independent cohorts of aged mice. One-way ANOVA followed by Tukey’s multiple comparisons post-test. ** p<0.01, *** p<0.001. **C.** Frequency of SiglecF^+^ cells amongst CD11c^hi^CD11b^lo^ macrophages obtained from 11-12 month old Cre^−^ or Cre^+^ mice. Symbols represent individual mice and error is S.D. Data are from 7 mice per group pooled from three independent cohorts of aged mice. Unpaired Student’s *t*-test, **p<0.01 **D.** Representative expression of CD11c and CD11b by Ly6C^lo^CD64^+^ macrophages obtained from 11-12 month old Cre^−^ or Cre^+^ mice. Symbols represent individual mice and error is s.d.. Data represent 7 mice per group pooled from three independent cohorts of aged mice. **E.** Frequency of Ki67^+^ cells amongst SiglecF-defined CD11c^hi^CD11b^lo^ macrophages obtained from 8 week old or 11-12 month old Cre^−^ or Cre^+^ mice. Symbols represent individual mice and error is s.d.. Data represent 7 (Cre^+^) or 8 (Cre^−^) mice per group (8 week old mice) or 5 mice per group (aged mice) pooled from two independent experiments. **F.** Log_2_-fold expression of genes that are implicated in lipid uptake or metabolism in alveolar macrophages as defined by *(9)*. Expression of *Egr2* is included as a reference. **G.** Bacterial levels (colony forming units, CFU) in the BAL fluid of Cre^−^ or Cre^+^ mice 14hrs after infection. Symbols represent individual mice and error is S.D. Data are from 10 (Cre^+^) or 12 (Cre^−^) mice per group pooled from three independent experiments. Mann Whitney test, *p<0.05. **H.** Frequency and absolute number of CD11c^hi^CD11b^lo^ alveolar macrophages in the BAL fluid of Cre^−^ or Cre^+^ mice 14hrs after infection. Symbols represent individual mice and error is S.D. Data are from 10 (Cre^+^) or 11 (Cre^−^) mice per group pooled from three independent experiments. Mann Whitney test, *p<0.05. **I.** Expression of *Marco*, *Wfdc10, Colec12* and *C3* from the RNA-seq dataset (*left panels*). Each symbol represents a biological replicate with four mice per group.

In an attempt to circumvent the confounding effects of these escapees, we generated a second strain to delete *Egr2* from macrophages by crossing *Egr2*^fl/fl^ mice with mice expressing ‘improved’ Cre recombinase under control of the endogenous *Fgcr1* promoter (*Fcgr1*^iCre^ *(3)*). By using *Fcgr1*^iCre^.*Rosa26*^LSL-RFP^ reporter mice, we confirmed that this approach led to efficient Cre recombination in alveolar macrophages, as well as in other tissue macrophages, but not in other leukocytes (**Supplementary Figure 5A**). Importantly, alveolar macrophages from *Fcgr1*^iCre^.*Egr2*^fl/fl^ mice phenocopied those from *Lyz2*^Cre^.*Egr2*^fl/fl^ mice (**Supplementary Figure 5B**), but the frequency of Cre escapees was markedly lower in *Fcgr1*^iCre^.*Egr2*^fl/fl^ mice compared with *Lyz2*^Cre^.*Egr2*^fl/fl^ mice (**Supplementary Figure 5C, D**). Despite this, we did not detect the development of proteinosis in aged *Fcgr1*^iCre^.*Egr2*^fl/fl^ mice compared to their littermate controls (**Supplementary Figure 5E**). Consistent with this, *Egr2* deficiency had little if any effect on the expression of genes associated with lipid uptake and metabolism that are characteristic of normal alveolar macrophages *(9)* (**Figure 4F**). Thus, while EGR2 is indispensable for the phenotypic identity of alveolar macrophages, it seems to be dispensable for regulating lipid handling.

We next sought to determine if EGR2-dependent differentiation controls protective immune functions of alveolar macrophages. To do so, we infected Cre^−^ (*Egr2*^fl/fl^) mice and Cre^+^ (*Lyz2*^Cre^.*Egr2*^fl/fl^) mice with 1×10^4^ colony forming units (CFU) *Streptococcus pneumoniae*, based on previous work showing that wild type alveolar macrophages efficiently clear infection at this dose *(30, 31)*. This showed that the majority of Cre^−^ mice (8/12) had cleared infection at 14 hours post infection, whereas the majority of Cre^+^ mice (8/10) had detectable bacteria in the airways at this timepoint (**Figure 4G**). Importantly, the failure to clear bacteria did not reflect the loss of tissue resident macrophages that can occur during inflammation or infection, as alveolar macrophages continued to dominate the airways in both Cre^+^ and Cre^−^ mice (**Figure 4H**). However, our RNA-seq analysis showed that expression of genes encoding molecules for the recognition, opsonisation and elimination of bacteria, including *Colec12, Wfdc10, C3* and *Marco*, the latter of which has been shown to be indispensable for immunity to *S. pneumoniae (32)*, were significantly reduced in *Egr2*-deficient alveolar macrophages (**Figure 4I**). Thus, EGR2-dependent differentiation is crucial for equipping alveolar macrophages with the machinery to capture and destroy pneumococci.

### EGR2 expression by alveolar macrophages is dependent on TGFβ and CSF2

Alveolar macrophages derive from foetal monocytes that seed the developing lung in the late gestational period *(8)*. To determine the point at which EGR2 is first expressed, we assessed EGR2 expression by E10.5 yolk sac macrophages, by macrophages in the embryonic lung (E16.5) and by CD11c^hi^CD11b^lo^ alveolar macrophages in the neonatal and adult lung using the ImmGen database. This revealed that *Egr2* was absent from yolk sac macrophages and macrophages in the embryonic lung at E16.5, but it was expressed by both neonatal and adult alveolar macrophages (**Figure 5A**), suggesting that it is induced during alveolarization in the neonatal period. Consistent with this, we found high expression of EGR2 at protein level by neonatal (d1) CD64^+^ lung macrophages (sometimes referred to as ‘pre-alveolar macrophages’) in Cre^−^ mice; as expected, this expression was deleted efficiently in Cre^+^ mice (**Figure 5B, C**). Importantly, Ly6C^hi^ monocytes in the lung of d1 neonatal mice lacked any expression of EGR2 (**Figure 5B, C**), reinforcing the selectivity of EGR2 expression even at this highly dynamic stage of myeloid cell development in the lung. Consistent with our analysis of mature alveolar macrophages in adult mice, *Egr2* deletion had no impact on the frequency and absolute number of pre-alveolar macrophages (**Figure 5D**). However, phenotypic differences were already apparent in Cre^+^ (*Lyz2*^Cre/+^.*Egr2*^fl/fl^) macrophages at this stage, with reduced CD11c and SiglecF expression which persisted into adulthood (**Figure 5E, F**). In parallel, EpCAM expression was absent from alveolar macrophages in the neonatal period and was progressively upregulated with age in an EGR2-dependent manner (**Figure 5F**). CD11b expression, which is downregulated in mature alveolar macrophages, was found on pre-alveolar macrophages in both Cre^−^ and Cre^+^ mice, and it was downregulated to the same extent with age in both strains.

**Figure 5:**
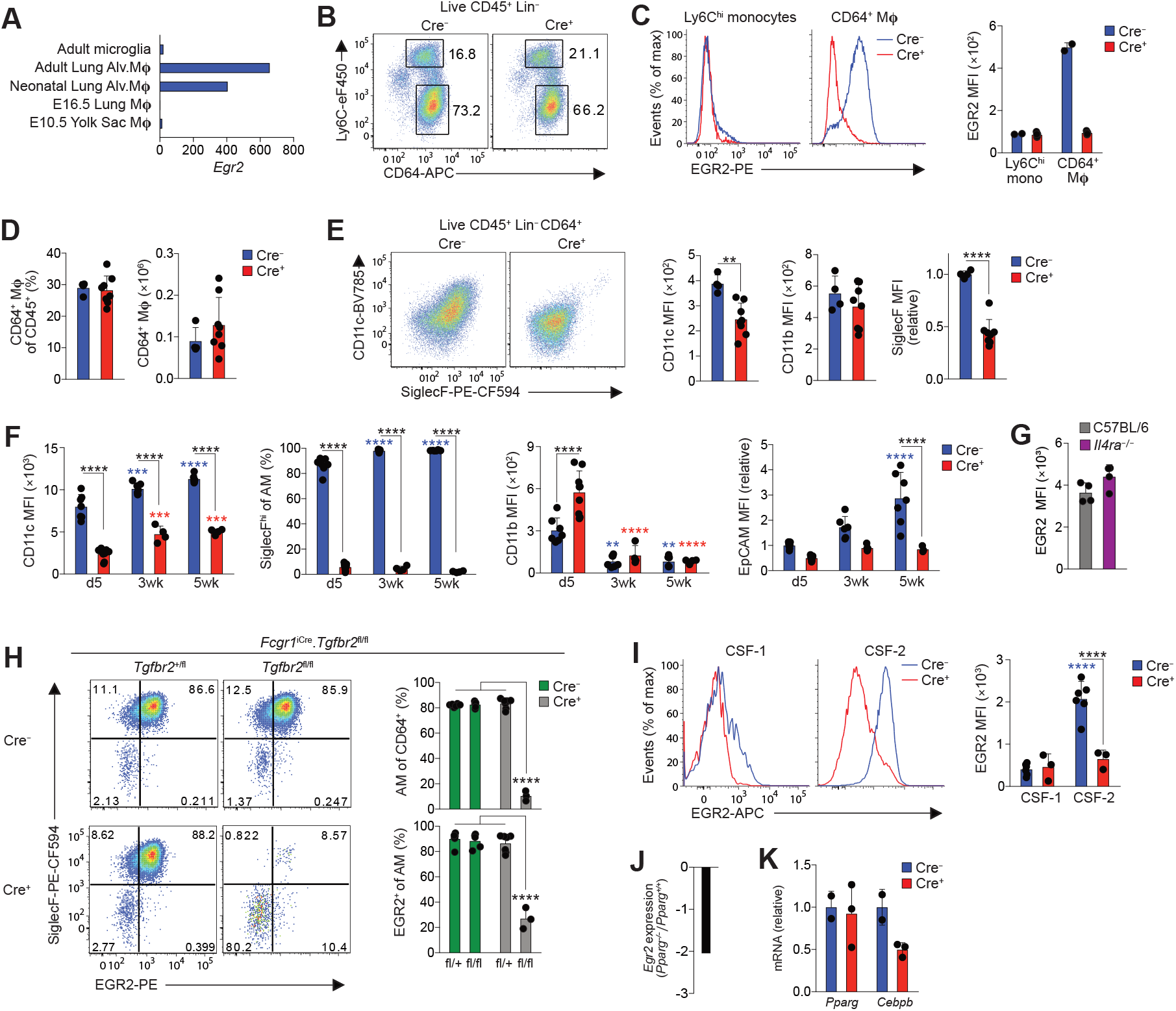
TGFβ and CSF2 drive EGR2 expression. **A.** Normalised expression (by DESeq2) of *Egr2* by the indicated populations (data obtained from the Immgen Consortium). **B.** Representative expression of Ly6C and CD64 by live CD45^+^CD3^−^CD19^−^Ly6G^−^ cells from the lungs of unmanipulated newborn Cre^−^ (*Egr2*^fl/fl^) or Cre^+^ (*Lyz2*^Cre/+^.*Egr2*^fl/fl^) mice. Data are pooled from one of two independent experiments performed. **C.** Histograms show representative expression of EGR2 by CD64^+^ ‘pre-alveolar macrophages’ and Ly6C^hi^ monocytes from the lungs of unmanipulated newborn Cre^−^ (*Egr2*^fl/fl^) or Cre^+^ (*Lyz2*^Cre/+^.*Egr2*^fl/fl^) mice and bar chart shows the mean fluorescent intensity (MFI) of EGR2 expression by these cells. **D.** Frequency and absolute number of CD64^+^ ‘pre-alveolar macrophages’ from mice in **B.** Symbols represent individual mice. Data are pooled from two independent experiments with 4 (Cre^−^) or 8 (Cre^+^) mice per group. **E.** FACS plots show representative expression of CD11c and SiglecF by CD64^+^ ‘pre-alveolar macrophages’ from mice in **B** and bar chart shows the MFI of CD11c, SiglecF and CD11b expression by these cells. Symbols represent individual mice. Data are pooled from two independent experiments with 4 (Cre^−^) or 8 (Cre^+^) mice per group. Unpaired Student’s *t*-test, **p<0.01, ****p<0.0001 **F.** MFI of CD11c, SiglecF, CD11b and EpCAM (relative to d5 Cre^−^ cells) expression by CD11c^hi^CD11b^lo^ alveolar macrophages obtained from unmanipulated *Egr2*^fl/fl^ (Cre^−^) or *Lyz2*^Cre^.*Egr2*^fl/fl^ (Cre^+^) mice at the indicated ages. Symbols represent individual mice. Coloured * denote significance between d5 and 3 and 5 weeks within the Cre^−^ (blue) and Cre^+^ (red) data. Data are pooled from two independent experiments with 4-9 mice per group. Two-way ANOVA with Tukey’s multiple comparisons test, ****p<0.0001 **G.** Representative expression of EGR2 by alveolar macrophages from adult WT (C57BL/6J) and *Il4ra*^−/−^ adult mice. Data from one experiment with 4 mice per group. **H.** Representative expression of EGR2 and SiglecF by CD11c^hi^CD11b^lo^ alveolar macrophages obtained from lungs of neonatal (d8) *Fcgr1*^iCre^.*Tgfbr2*^fl/fl^ and Cre^−^ and *Tgfbr2*^fl/+^ littermate controls. Bar charts show the mean frequencies of CD11c^hi^CD11b^lo^ alveolar macrophages of all Ly6C^lo^CD64^+^ cells (*upper*) and EGR2^+^ cells amongst CD11c^hi^CD11b^lo^ alveolar macrophages (*lower*). Symbols represent individual mice. Data are pooled from two independent experiments with 3-7 mice per group. **** p<0.0001 (One-way ANOVA followed by Tukey’s multiple comparisons post-test). **I.** Representative expression of EGR2 (*left*) and MFI of EGR2 (*right*) by FACS-purified Ly6C^hi^ monocytes cultured *in vitro* with recombinant CSF-1 (20ng/ml) or CSF-2 (20ng/ml) for five days. Symbols represent monocytes isolated from individual mice and horizontal lines represent the mean. Data are from 6 Cre^−^ (*Egr2*^fl/fl^) or 3 Cre^+^ (*Lyz2*^Cre^.*Egr2*^fl/fl^) mice per group pooled from two independents experiment. Two-way ANOVA followed by Tukey’s multiple comparisons post-test, **** p<0.0001. Coloured * denote significance between CSF-1 and CSF-2 within the Cre^−^ (blue) and Cre^+^ (red) data. **J.** Relative expression of *Egr2* by alveolar macrophages obtained from *Pparg*^fl/fl^ or *Itgax*^Cre^.*Pparg*^fl/fl^ mice from *(9)*. **K.** qPCR analysis of *Pparg* and *Cebpb* mRNA by BAL cells from unmanipulated adult *Egr2*^fl/fl^ (Cre^−^) or *Lyz2*^Cre^.*Egr2*^fl/fl^ (Cre^+^) mice. Data represent 2 *Egr2*^fl/fl^ (Cre^−^) or 4 *Lyz2*^Cre^.*Egr2*^fl/fl^ (Cre^+^) mice per group.

We next set out to determine the environmental factors that drive EGR2 expression. Many studies employing *in vitro* culture systems have described EGR2 expression as a feature of ‘alternatively activated’ macrophages, dependent on IL-4R signalling *(33–35)*. Importantly, expression of EGR2 by alveolar macrophages was independent of IL-4R signalling (**Figure 5G & Supplementary Figure 5F**), as were key EGR2-dependent phenotypic traits, such as SiglecF and EpCAM expression (**Supplementary Figure 5F**). TGF-β has recently been shown to be crucial for the development of alveolar macrophages *(11)* and thus we next explored if the TGF-β-TGF-βR axis drives expression of EGR2. To do so, we generated a new mouse line by crossing *Fcgr1*^iCre^ mice to mice with LoxP sites flanking the *Tgfbr2* allele (*Tgfbr2*^fl/fl^). Consistent with the crucial role for TGF-βR in controlling alveolar macrophage development *(11)*, there was a paucity of alveolar macrophages in the lungs of neonatal *Fcgr1*^iCre/+^.*Tgfbr2*^fl/fl^ compared with *Fcgr1*^+/+^.*Tgfbr2*^fl/fl^ and *Fcgr1*^iCre/+^.*Tgfbr2*^fl/+^ controls (**Figure 5H**). Strikingly, while CD11c^+^CD11b^lo^ alveolar macrophages expressed high levels of EGR2 in control groups, EGR2 expression was largely abolished in *Fcgr1*^iCre/+^.*Tgfbr2*^fl/fl^ mice, demonstrating that TGF-βR signalling is vital for EGR2 induction *in vivo*. As *Fcgr1*^iCre/+^.*Tgfbr2*^fl/fl^ developed fatal seizures between d14 and d21 of age, perhaps reflecting the indispensable role for TGF-βR in controlling microglia activity *(37–39)*, we were unable to carry out further analyses using this strain.

Given the central role for CSF-2 in alveolar macrophage development, we also assessed the role of CSF-2 in driving EGR2 expression using an *in vitro* culture system in which Ly6C^hi^ monocytes from mouse bone marrow were FACS-purified and cultured with recombinant CSF-1 or CSF-2. This revealed that CSF-2 was also capable of driving EGR2 expression in this system (**Figure 5I**). Given that CSF-2R and TGF-βR signalling is known to induce expression of PPAR-γ *(9, 11)*, we next determined if PPAR-γ (encoded by *Pparg*) is upstream of EGR2. Analysis of a published dataset comparing the transcriptional profile of *Pparg*-sufficient and - deficient lung macrophages revealed significant downregulation (2.1-fold change) of *Egr2* in the context of *Pparg* deficiency (**Figure 5j**). In contrast, *Pparg* expression was unaffected in alveolar macrophages from *Lyz2*^Cre^.*Egr2*^fl/fl^ mice (**Figure 5K**), suggesting EGR2 is downstream of PPAR-γ. Another transcription factor implicated in controlling alveolar macrophage differentiation is C/EBPβ *(14)* and EGR2 has been shown to modulate C/EBPβ *in vitro (33)*. In our hands, *Egr2-*deficiency led to reduced expression of C/EBPβ at mRNA (**Figure 5K**) and protein level (**Supplementary Figure 5G**). Taken together, these data support the premise that EGR2 expression by alveolar macrophages is induced by TGF-β and CSF-2 in a PPAR-γ-dependent manner in the neonatal period and this in turn induces expression of C/EBPβ to drive tissue-specific differentiation.

### *Egr2* deficiency confers a competitive disadvantage on alveolar macrophages

Given the observation that EGR2-sufficient alveolar macrophages come to dominate the *Lyz2*^Cre^.*Egr2*^fl/fl^ (Cre^+^) mice airspace, we next set out to determine if EGR2 deletion confers an intrinsic competitive disadvantage on alveolar macrophages. To this end, we generated mixed bone marrow chimeric mice by reconstituting lethally irradiated WT (CD45.1^+^/.2^+^) mice with a 1:1 ratio of WT (CD45.1^+^) and either Cre^−^ (*Egr2*^fl/fl^) or Cre^+^ (*Lyz2*^Cre^.*Egr2*^fl/fl^) (CD45.2^+^) bone marrow cells (**Figure 6A**). 8 weeks after reconstitution, we found that *Egr2* deficient (Cre^+^) and *Egr2* sufficient (Cre^−^) bone marrow contributed equally to blood monocytes and neutrophils (data not shown), as they did to the pools of monocytes, interstitial macrophages and dendritic cell subsets in the lung (**Figure 6B, C**). In contrast, alveolar macrophages were derived almost exclusively from WT BM in WT:Cre^+^ chimeric mice, whereas they were derived equally from both BM sources in WT:Cre^−^ chimeric mice (**Figure 6B-D**). The mixed BM chimeric model also confirmed that the phenotypic and morphologic differences seen in intact *Lyz2*^Cre^.*Egr2*^fl/fl^ mice were due to cell intrinsic loss of EGR2, rather than effects of *Egr2* deficiency on the lung environment (**Figure 6E, F**). We also used this system to confirm the reduced expression of C/EBPβ by alveolar macrophages deriving from *Lyz2*^Cre^.*Egr2*^fl/fl^ (Cre^+^) bone marrow (**Figure 6F**). Taken together, these results demonstrate that cell intrinsic EGR2 is indispensable for the differentiation of alveolar macrophages and repopulation of the alveolar niche following radiation-induced depletion.

**Figure 6:**
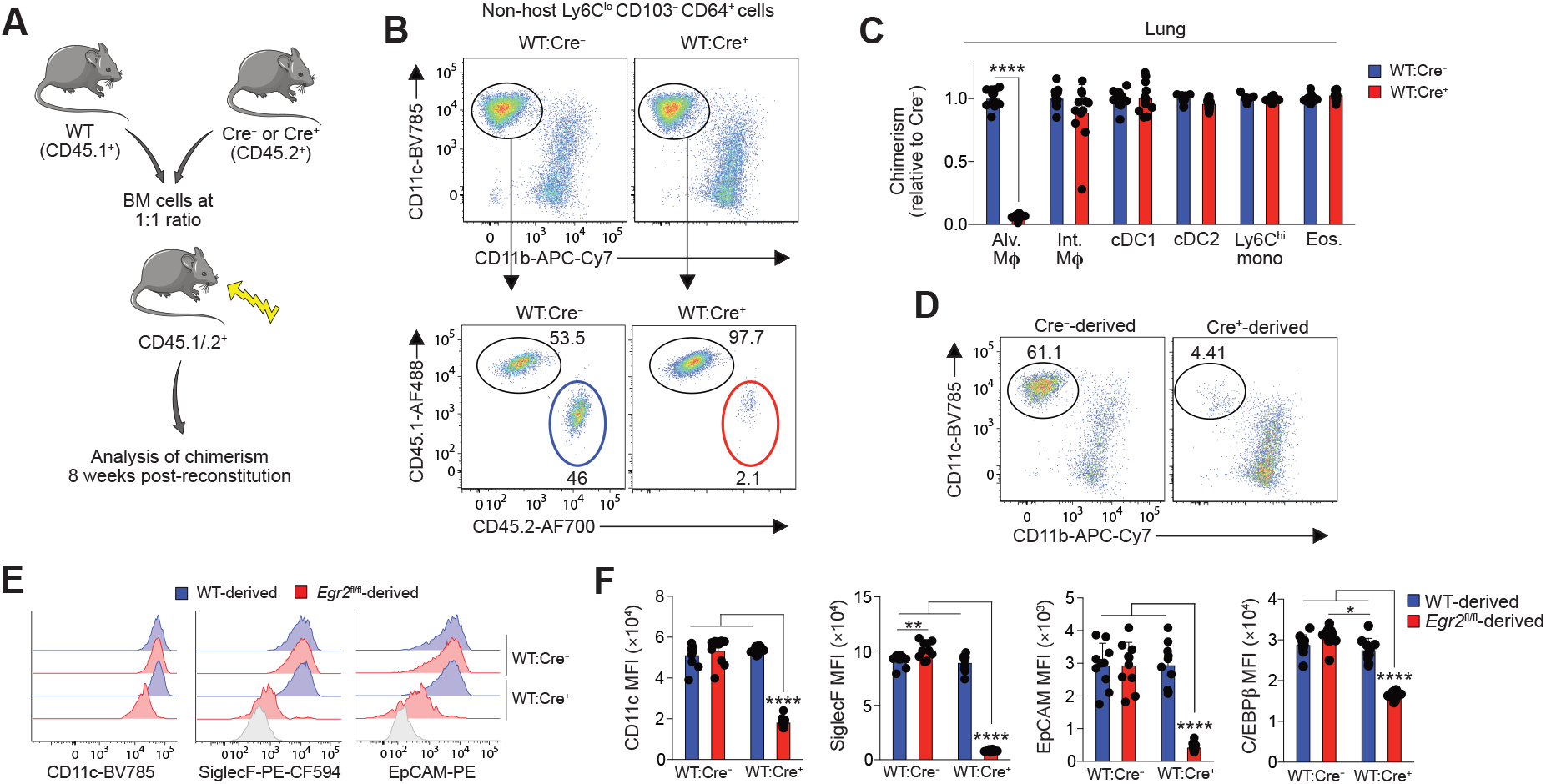
*Egr2* deficiency confers a competitive disadvantage on alveolar macrophages. **A.** Schematic of the generation of mixed bone marrow chimeric mice using Cre^−^ (*Egr2*^fl/fl^) or Cre^+^ (*Lyz2*^Cre^.*Egr2*^fl/fl^) mice as donors. **B.** Representative expression of CD11c and CD11b by Ly6C^lo^CD64^+^ macrophages amongst live CD45^+^CD3^−^CD19^−^Ly6G^−^CD103^−^ cells (*upper panels*) and representative expression of CD45.1 and CD45.2 by CD11c^hi^CD11b^lo^ alveolar macrophages (*lower panels*) from WT:Cre^−^ or WT:Cre^−^ chimeric mice. **C.** Contribution of *Egr2*^fl/fl^ BM to the indicated lung myeloid populations in WT:Cre^+^ chimeric mice relative to WT:Cre^−^ mice. Chimerism was normalised to Ly6C^hi^ blood monocytes before normalisation of Cre^+^ to Cre^−^. Symbols represent individual mice. Data are from 12 (WT:Cre^+^) or 13 (WT:Cre^−^) mice per group pooled from two independent experiments. Student’s *t*-test with Holm-Sidak correction, **** p<0.0001. **D.** Representative composition of the CD64^+^ macrophage compartment deriving from Cre^−^ or Cre^+^ bone marrow in WT:Cre^−^ or WT:Cre^−^ chimeric mice. **E.** Representative expression of CD11c, SiglecF and EpCAM by WT- and *Egr2*^fl/fl^-derived alveolar macrophages in WT:Cre^−^ or WT:Cre^−^ chimeric mice. Shaded histograms represent FMO controls. **F.** Mean fluorescent intensity (MFI) of CD11c, SiglecF, EpCAM and C/EBPβ expression by WT- and *Egr2*^fl/fl^-derived alveolar macrophages in WT:Cre^−^ or WT:Cre^−^ chimeric mice. Symbols represent individual mice. Data are from 10 mice per group from one experiment of two performed. One-way ANOVA followed by Tukey’s multiple comparisons post-test, **** p<0.0001.

### Bone marrow-derived monocytes replenish the alveolar macrophage niche following lung injury

Loss of tissue resident macrophages is a frequent consequence of inflammation, including in the lung *(39)*. Thus, given that *Egr2*-deficient macrophages failed to replenish the alveolar niche following radiation treatment, we next sought to determine if EGR2 plays a role in macrophage repopulation following lung injury. The chemotherapeutic agent bleomycin is a common model of chronic lung injury and self-resolving pulmonary fibrosis *(40)*, which is characterised by initial loss of alveolar macrophages during the inflammatory phase (day 7), followed by repopulation during the fibrotic and resolution phases (from day 14 onwards) (**Figure 7A**). Post-injury repopulation of macrophages can occur via two mechanisms: *in situ* proliferation of resident cells and/or replenishment by bone marrow-derived monocytes. To determine if bone marrow-derived monocytes contribute to the alveolar macrophage compartment following bleomycin-induced injury, we used tissue protected bone marrow chimeric mice to assess replenishment kinetics without exposing the lung to the additional insult of ionising radiation (**Figure 7B**). Consistent with previous studies *(41)*, we found that bleomycin instillation led to progressive replacement of resident alveolar macrophages by BM-derived cells, with the entire alveolar macrophage compartment being replaced at 32 weeks post injury (**Figure 7B**). Interestingly, recently arrived, monocyte-derived alveolar macrophages expressed low-intermediate levels of SiglecF, with acquisition of SiglecF requiring long-term residence in the airway (**Figure 7C)**.

**Figure 7:**
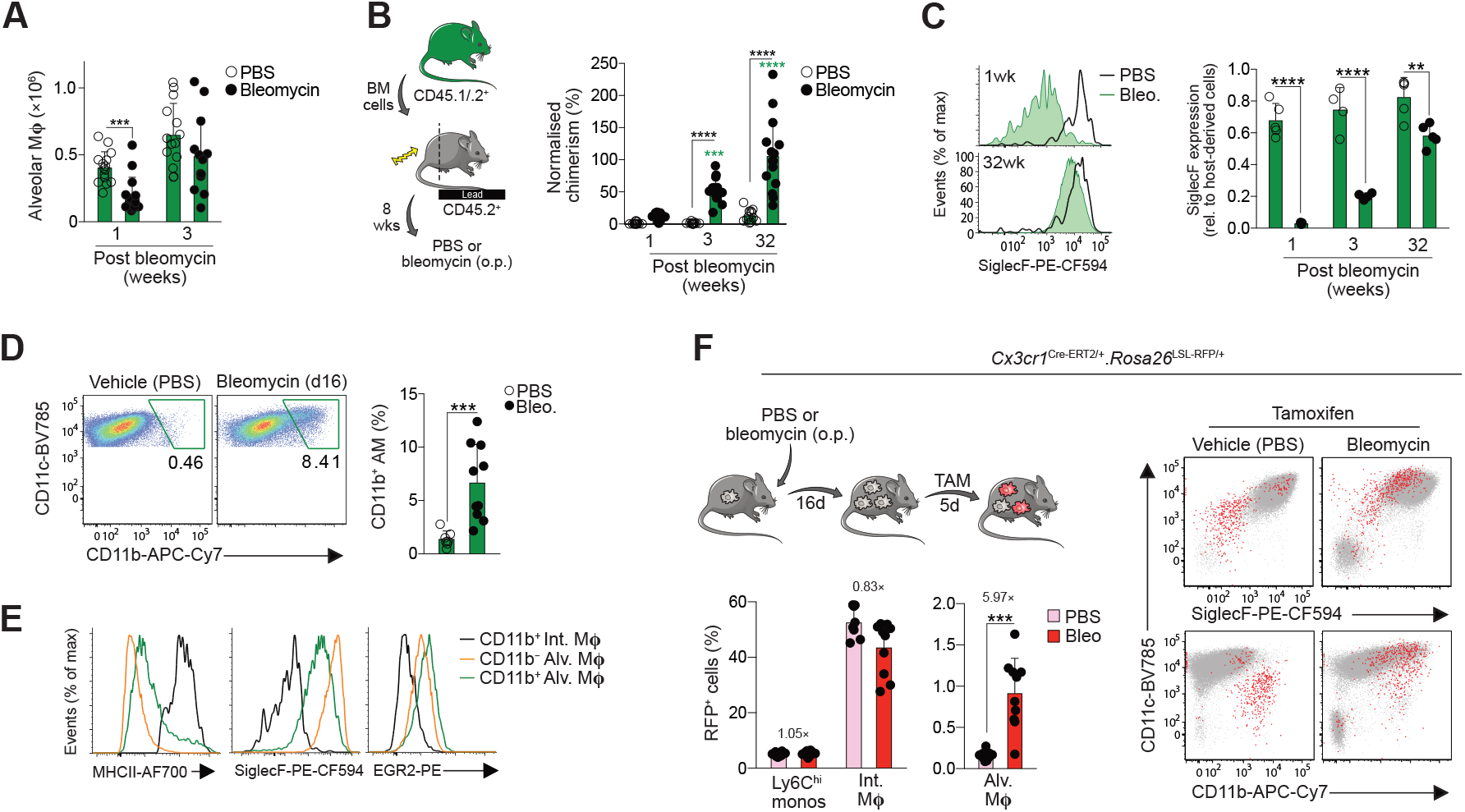
Monocyte-derived, CX3CR1^+^ interstitial macrophages can replenish the alveolar macrophage niche following injury. **A.** Absolute numbers of alveolar macrophages 1- and 3-weeks following bleomycin administration or PBS vehicle control. Symbols represent individual mice and error is S.D. Data are pooled from two independent experiments at each time point with 13-15 mice per group. Student’s *t* test with Holm-Sidak correction, ***p<0.001. **B.** Non-host chimerism of alveolar macrophages in tissue protected bone marrow chimeric mice at 1-, 3- or 32-weeks following administration of bleomycin or PBS vehicle control. Chimerism is normalised to Ly6C^hi^ blood monocytes. Symbols represent individual mice and error is S.D. Data are pooled from two independent experiments at each time point with 13-15 mice per group. Two-way ANOVA with Tukey’s multiple comparisons test. ***p<0.001, ****p<0.0001. **C.** Expression of SiglecF by CD11c^hi^CD11b^lo^ alveolar macrophages from the lung of mice in **B**. at 1 week and 32 weeks post bleomycin or PBS administration. Symbols represent individual mice and error is S.D. Data are from one of two independent experiments at each time point with 4 mice per group. Two-way ANOVA with Tukey’s multiple comparisons test. **p<0.01, ****p<0.0001. **D.** Representative expression of CD11c and CD11b by CD11c^hi^CD64^+^ cells obtained by BAL from WT mice two weeks after instillation of bleomycin or vehicle control (*left)*. Graph shows the mean frequency of CD11b^+^ alveolar macrophages (*right*). Symbols represent individual mice and error is S.D. Data are pooled from two independent experiments with 7 (Cre^−^) or 10 (Cre^+^) mice per group. Mann Whitney test, ***p<0.001. **E.** Representative expression of MHCII, SiglecF and EGR2 by CD11b^+^ interstitial macrophages and CD11b-defined CD11c^hi^ alveolar macrophages from mice in **D.** **F.** Experimental scheme for the induction of lung injury and tamoxifen administration in *Cx3cr1*^Cre-ERT2/+^.*Rosa26*^LSL-RFP/+^ fate mapping mice. Lower graphs show the levels of recombination in Ly6C^hi^ monocytes, CD64^+^ interstitial macrophages and alveolar macrophages from *Cx3cr1*^Cre-ERT2/+^.*Rosa26*^LSL-RFP/+^ mice administered bleomycin or vehicle control. Representative expression of CD11c, SiglecF and CD11b by RFP^+^ (red) or RFP^−^ (grey) cells present in the BAL fluid of *Cx3cr1*^Cre-ERT2/+^.*Rosa26*^LSL-RFP/+^ mice 3 weeks after bleomycin or vehicle instillation. Graphs show the mean fluorescent intensity (MFI) of CD11c and SiglecF expression by RFP^+^ cells. Symbols represent individual mice and error is S.D. Mann Whitney test, ***p<0.001.

We next interrogated this process further to determine if interstitial macrophages that accumulate in the lung parenchyma during injury can subsequently mature into alveolar macrophages during tissue repair *(41, 42)*. Indeed, during the recovery phase of disease, we noted the presence of cells with features of both alveolar and interstitial macrophages (CD11c^hi^CD11b^+^ MHCII^+^CD64^hi^) in the BAL fluid (**Figure 7D**), and these cells expressed intermediate levels of SiglecF (**Figure 7E**), indicative of recent monocyte origin. To examine the relationship of these intermediate cells to interstitial macrophages more directly, we performed fate mapping studies using *Cx3cr1*^Cre-ERT2/+^.*Rosa26*^LSL-RFP/+^ reporter mice, in which administration of tamoxifen leads to irreversible expression of RFP by CX3CR1 expressing cells *(43, 44)* (**Figure 7F**). Interstitial macrophages are characterised by high expression of CX3CR1 and administration of tamoxifen led to labelling of 40-50% of interstitial macrophages in both healthy lung and at d21 post bleomycin administration (**Figure 7F**). No recombination was seen in *Cx3cr1*^Cre-ERT2/+^.*Rosa26*^LSL-RFP/+^ mice in the absence of tamoxifen (**Supplementary Figure 6A**). Although very low levels of recombination were detected in control alveolar macrophages, a clear population of RFP^+^ cells could be detected in the BAL of the recipients of bleomycin following tamoxifen treatment (**Figure 7F**). As monocytes are poorly labelled in this system and *Cx3cr1* levels did not change in *bona fide* resident alveolar macrophages in response to bleomycin treatment (**Supplementary Figure 6B**), these RFP^+^ cells must represent fate-mapped, monocyte-derived CX3CR1^+^ cells. In line with this, RFP^+^ cells had the ‘hybrid’ alveolar/interstitial CD11c^hi^CD11b^+^SiglecF^int^ profile, supporting the idea that these represent transitional cells (**Figure 7F**). Thus, following bleomycin-induced injury, the alveolar macrophage compartment is restored by monocytes that transition through a CX3CR1^hi^ state.

### EGR2 is indispensable for alveolar macrophage repopulation and tissue repair following lung injury

Given that transitional CD11b^+^SiglecF^int^ cells also expressed EGR2, contrasting with its restriction to SiglecF^hi^ alveolar macrophages in health (**Figure 7E**), we examined whether EGR2 is necessary for the replenishment of the alveolar niche during recovery from bleomycin-induced injury. To do this, we administered bleomycin to *Lyz2*^Cre^.*Egr2*^fl/fl^ mice and their Cre^−^ littermates and assessed macrophage dynamics in total lung digests. The inflammatory phase of this disorder (day 7) was associated with accumulation of CD11b^+^ macrophages and this occurred to the same extent in Cre^*–*^ and Cre^*+*^ mice (**Figure 8A, B**). Consistent with recent reports *(45)*, the CD11b^+^CD64^+^ interstitial macrophage population was heterogeneous during the fibrotic phase of disease (d14-d21), with MHCII^+^ and MHCII^lo^CD36^+^Lyve1^+^ subsets. This pattern was identical in between Cre^−^ and Cre^+^ groups (**Supplementary Figure 7A, C**), as were the numbers of Ly6C^hi^ monocytes and neutrophils (**Supplementary Figure 7A, B**). We did however detect a significant reduction in eosinophils in the lung of *Lyz2*^Cre^.*Egr2*^fl/fl^ mice compared with Cre^−^ littermates, despite eosinophils lacking EGR2 expression (**Supplementary Figure 7D, E**).

**Figure 8:**
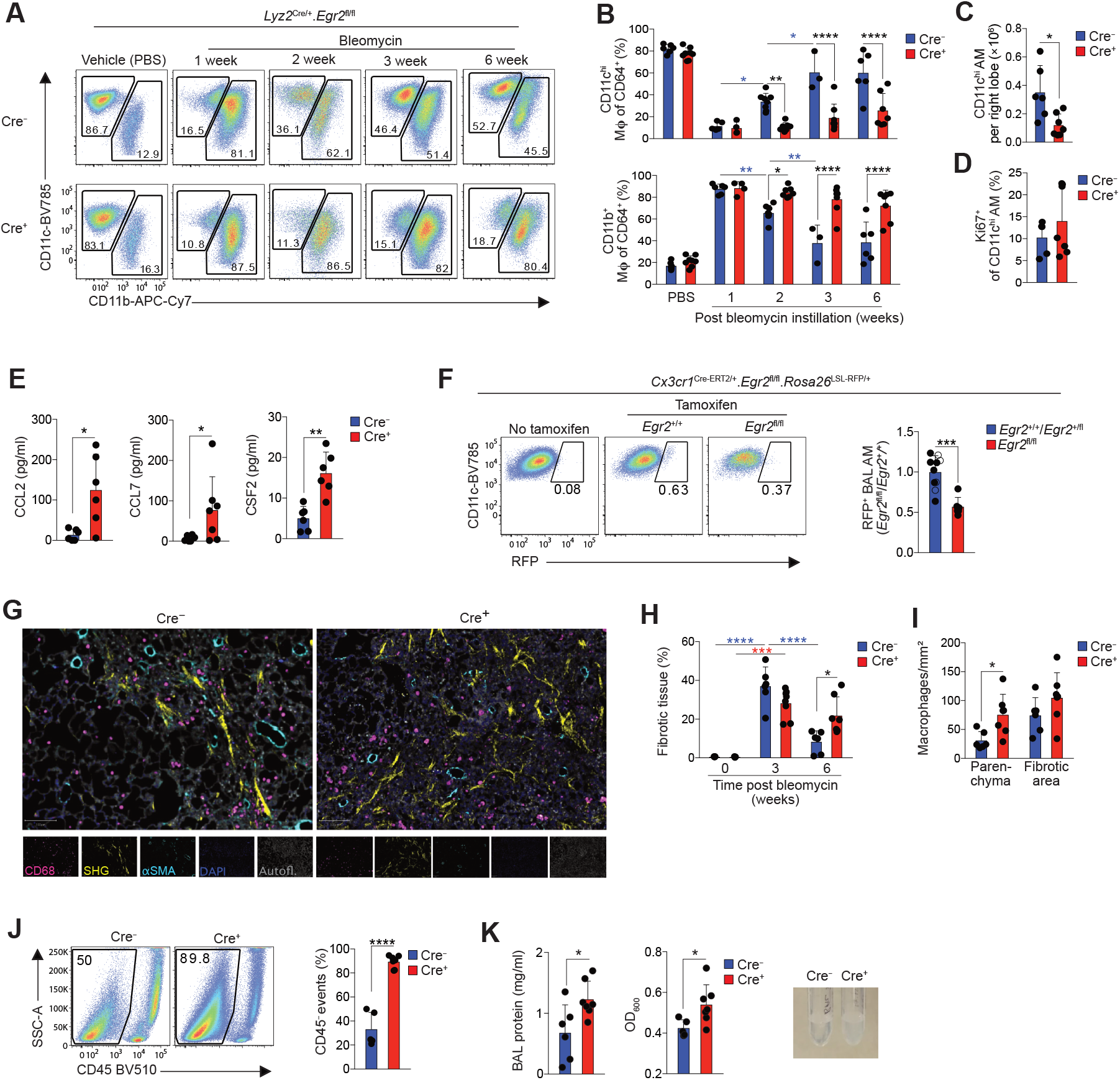
EGR2 is indispensable for the repopulation of the alveolar macrophage niche and tissue repair following lung injury. **A.** Representative expression of CD11c and CD11b by live CD45^+^CD3^−^CD19^−^Ly6G^−^CD64^+^ cells from the lungs of *Egr2*^fl/fl^ (Cre^−^) and *Lyz2*^Cre/+^.*Egr2*^fl/fl^ (Cre^+^) mice at 1, 2, 3 or 6 weeks post bleomycin or vehicle controls. **B.** Frequency of CD11c^hi^CD11b^lo^ alveolar macrophages and CD11c^var^CD11b^+^ cells from mice in **A.** Symbols represent individual mice and error is the S.D. Data are pooled from at least two independent experiments at each time point with 3-7 mice per group. Two-way ANOVA with Tukey’s multiple comparisons test, *p<0.05. **p<0.01, p<0.001, ****p<0.0001 **C.** Absolute number of CD11c^hi^CD11b^lo^ alveolar macrophages in lungs six weeks post bleomycin instillation. Symbols represent individual mice and error is the S.D. Data are pooled from two independent experiments with 6 (Cre^−^) or 7 (Cre^+^) mice per group. Mann Whitney test, *p<0.05. **D.** Frequency of Ki67^+^ CD11c^hi^CD11b^lo^ alveolar macrophages in lungs six weeks post bleomycin instillation. Symbols represent individual mice and error is the S.D. Data are pooled from two independent experiments with 6 (Cre^−^) or 7 (Cre^+^) mice per group. **E.** CCL2, CCL7 and CSF2 levels in BAL fluid obtained from *Egr2*^fl/fl^ (Cre^−^) and *Lyz2*^Cre/+^.*Egr2*^fl/fl^ (Cre^+^) mice six weeks post bleomycin instillation. Symbols represent individual mice and error is the S.D. Data are pooled from two independent experiments with 6 mice per group. Mann Whitney test (CCL2, CCL7), *p<0.05, unpaired Student’s t test (CSF2), **p<0.01. **F.** Representative expression of RFP by CD11c^hi^CD64^+^ alveolar macrophages present in the BAL fluid of *Cx3cr1*^Cre-ERT2/+^.*Rosa26*^LSL-RFP/+^.*Egr2*^fl/fl^ and their *Cx3cr1*^Cre-ERT2/+^.*Rosa26*^LSL-RFP/+^.*Egr2*^+/+^ (open circles) or *Cx3cr1*^Cre-ERT2/+^.*Rosa26*^LSL-RFP/+^.*Egr2*^fl/+^ (solid circles) controls 3 weeks following instillation of bleomycin or vehicle control. Graph shows the relative frequency of RFP^+^ alveolar macrophages present in the BAL fluid. Symbols represent individual mice and error is the S.D. Data are pooled from two independent experiments at each time point with 10 (*Egr2*^+/+^ [open symbols]/*Egr2*^fl/+^ [filled symbols]) or 6 (*Egr2*^fl/fl^) per group. Unpaired Student’s t test, ***p<0.001. **G.** 2-photon fluorescence imaging of lung tissue from adult *Egr2*^fl/fl^ (Cre^−^) and *Lyz2*^Cre/+^.*Egr2*^fl/fl^ (Cre^+^) mice 6 weeks following bleomycin administration. Sections were stained with CD68, αSMA and DAPI. Autofluorescence is depicted in grey and collagen was detected by second harmonic generation (SHG). **H.** Quantification of fibrotic score of lung tissue from *Egr2*^fl/fl^ (Cre^−^) and *Lyz2*^Cre/+^.*Egr2*^fl/fl^ (Cre^+^) 3 or 6 weeks following bleomycin administration or PBS controls (from 3 week time point). See **Supplementary Figure 8.** Symbols represent individual mice and error is the S.D. Data are pooled from two independent experiments from one experiment with 6 (Cre^−^) or 7 (Cre^+^) mice per group. Two-way ANOVA followed by Tukey’s multiple comparisons test, *p<0.05, **p<0.01, ***p<0.001, ****p<0.0001. **I.** Quantification of macrophage density in the parenchyma and fibrotic areas of lung tissue from *Egr2*^fl/fl^ (Cre^−^) and *Lyz2*^Cre/+^.*Egr2*^fl/fl^ (Cre^+^) 6 weeks following bleomycin administration. See **Supplementary Figure 8.** Symbols represent individual mice and error is the S.D. Data are pooled from two independent experiments with 6 (Cre^−^) or 7 (Cre^+^) mice per group. Student’s t test with Holm-Sidak correction for multiple tests, *p<0.05. **J.** SSC-A profile and expression of CD45 by BAL obtained from *Egr2*^fl/fl^ (Cre^−^) and *Lyz2*^Cre/+^.*Egr2*^fl/fl^ (Cre^+^) 6 weeks following bleomycin administration. Graph shows the mean frequency of CD45^+^ cells amongst all live, single events. Symbols represent individual mice and error is the S.D. Data are pooled from two independent experiments with 5 (Cre^−^) or 7 (Cre^+^) mice per group. Unpaired Student’s t test, ****p<0.0001. **K.** Total protein concentration (*left*), turbidity (*centre*) and representative pictures (*right*) of BAL fluid from *Egr2*^fl/fl^ (Cre^−^) and *Lyz2*^Cre/+^.*Egr2*^fl/fl^ (Cre^+^) 6 weeks following bleomycin administration. Symbols represent individual mice. Data are pooled from two independent experiments from one experiment with 6 (Cre^−^) or 7 (Cre^+^) mice per group. Mann Whitney test, *p<0.05.

A reduction in alveolar macrophages was observed in both groups on day 7 after administration of bleomycin. Although this began to be restored by day 14 in Cre^−^ (*Egr2*^fl/fl)^ control mice, this did not occur in Cre^+^ (*Lyz2*^Cre^.*Egr2*^fl/fl^) mice and indeed, the alveolar macrophage compartment remained significantly reduced in Cre^+^ mice compared with Cre^−^ littermates even after 6 weeks (**Figure 8A-C**), suggesting EGR2 is indispensable for the repopulation of the alveolar macrophage niche following bleomycin-induced injury. The lack of repopulation in Cre^+^ mice did not appear to reflect an inability of *Egr2* deficient macrophages to proliferate, as the proportion of Ki67^+^ proliferating cells was equivalent in Cre^−^ and Cre^+^ mice (**Figure 8D**). Equally, this also did not reflect a lack of chemoattractants in the airways to recruit monocyte-derived cells, as both CCL2 and CCL7 were actually elevated in Cre^+^ mice compared with Cre^−^ littermates (**Figure 8E**). Similarly, CSF2 levels were elevated in the BAL fluid of Cre^+^ mice, ruling out the possibility that lack of appropriate growth factors is responsible for defective alveolar macrophage differentiation in the absence of EGR2 (**Figure 8E**). Instead, these data suggested that *Egr2* deficiency led to an intrinsic inability of bone marrow-derived cells to repopulate the macrophage niche. To test this directly, we crossed *Cx3cr1*^Cre-^ ERT2/+.*Rosa26*^LSL-RFP/+^ mice with *Egr2*^fl/fl^ mice to allow for temporal RFP labelling of CX3CR1-expressing cells and *Egr2* deficiency in the same animal. We administered tamoxifen during the period of alveolar macrophage reconstitution (d16 to d21) and assessed the presence of RFP-labelled cells amongst alveolar macrophages. Compared with tamoxifen-treated controls (*Cx3cr1*^Cre-ERT2/+^.*Rosa26*^LSL-RFP/+^.*Egr2*^+/+^ or *Cx3cr1*^Cre-ERT2/+^.*Rosa26*^LSL-RFP/+^.*Egr2*^fl/+^ mice), we found a marked reduction in the frequency of RFP^+^ alveolar macrophages in the BAL of tamoxifen treated *Cx3cr1*^Cre-ERT2/+^.*Rosa26*^LSL-RFP/+^.*Egr2*^fl/fl^ mice during lung repair (**Figure 8F**), demonstrating that EGR2 controls the post-injury repopulation of the alveolar macrophage compartment by CX3CR1^+^ cells.

To determine the consequence of the failure of *Egr2* deficient cells to reconstitute the alveolar niche, we assessed the fibrotic response and subsequent repair processes in *Lyz2*^Cre^.*Egr2*^fl/fl^ mice. Notably, we did not detect differences in the degree of fibrosis or expression of key genes associated with fibrosis, including *Col3a1* and *Pdgfrb* between untreated *Lyz2*^Cre^.*Egr2*^fl/fl^ mice and their *Egr2*^fl/fl^ littermate controls at day 21, a time considered ‘peak’ fibrosis (**Figure 8H, Supplementary Figure 8A, B**). However, analysis at 6 weeks post bleomycin showed that whereas the Cre^−^ mice had largely repaired their lungs, Cre^+^ mice had defective repair evidenced by persistent fibrosis and architectural damage (**Figure 8G, H, Supplementary Figure 8A**). This was paralleled by elevated numbers of macrophages in the lung parenchyma (**Figure 8I, Supplementary Figure 8A**) and parenchymal macrophage persistence correlated with the degree of fibrosis (**Supplementary Figure 8C**). Furthermore, homeostasis failed to be restored in the airways. Flow cytometric analysis of BAL fluid revealed that CD45^+^ leukocytes comprised only 10% of all events in Cre^+^ mice compared with ~60% in their Cre^−^ littermates (**Figure 8J**). The vast majority of the CD45^−^ fraction failed to express signature markers for cells of epithelial, endothelial or fibroblast origin, suggesting this may represent cellular debris, which could also be found amongst lung digests (**Supplementary Figure 9A, B**). This was paralleled by elevated BAL fluid protein levels and turbidity in the Cre^+^ mice compared with Cre^−^ controls, suggesting that the inability to replenish the alveolar macrophage niche following injury was associated with the development of alveolar proteinosis (**Figure 8K**). Thus, loss of EGR2-dependent, monocyte-derived alveolar macrophages leads to defective tissue repair, persistent cellular damage and failed restoration of lung homeostasis.

## Discussion

Given the multifaceted role of macrophages in tissue homeostasis, inflammation and tissue repair, as well as many chronic pathologies, understanding the environmental signals and the downstream molecular pathways that govern macrophage differentiation is a key objective in the field of immunology. Here, we identify the transcription factor EGR2 as a selective and indispensable part of the tissue-specific differentiation of lung alveolar macrophages.

Our transcriptomic analysis identified EGR2 as a feature of murine lung alveolar macrophages, a finding consistent with previous studies using bulk transcriptomics to compare macrophages from different tissues *(1, 6)* and a recent study using a similar scRNA-seq based approach, which identified EGR2 specifically upregulated in alveolar, and not interstitial, macrophages *(46)*. Importantly, EGR2 is also expressed by alveolar macrophages in rats *(47)* and we showed that EGR2 is a feature of alveolar macrophages in man, a finding consistent with recent cross-species analysis *(34)*. Thus, EGR2 appears to represent an evolutionarily conserved transcriptional regulator of alveolar macrophages.

While EGR2 has been implicated in controlling monocyte to macrophage differentiation in the past, these studies have often reached discrepant conclusions *(19, 20, 34)*. This could reflect the fact that most studies examining the role of EGR2 in monocyte-macrophage differentiation have employed *in vitro* culture systems due to the postnatal lethality of global *Egr2* deficient mice *(21, 22)*. By generating *Lyz2*^Cre^.*Egr2*^fl/fl^ and *Fcgr1*^iCre^.*Egr2*^fl/fl^ mice, we circumvented this lethality and allowed for myeloid- and macrophage-specific deletion of EGR2, respectively. Our analysis showed that only macrophages in the airways were affected by *Egr2* deletion and our transcriptional profiling and extensive phenotypic characterisation demonstrated that EGR2 controls a large proportion of the alveolar macrophage ‘signature’, including key phenotypic traits such as SiglecF, EpCAM and TREM1. This is consistent with recent epigenetic analysis showing an overrepresentation of EGR motifs in the genes defining alveolar macrophages *(2, 46)*. Importantly, although previous work has suggested that there is redundancy between EGR family members, specifically EGR1 and EGR2, we found *Egr1* expression was unaffected by EGR2 deficiency and was unable to rescue alveolar macrophage differentiation. Indeed, consistent with a recent study *(46)*, we found EGR1 to be expressed at low levels by alveolar macrophages, but higher by interstitial macrophages. Thus, it is plausible that distinct EGR family members may be involved in the differentiation of anatomically distinct lung macrophages.

Notably, if assessed simply on the basis of their unique CD11c^hi^CD11b^lo^ profile, the absolute number of alveolar macrophages was equivalent between adult Cre^−^ (*Egr2*^fl/fl^) and Cre^+^ (*Lyz2*^Cre^.*Egr2*^fl/fl^) mice. This could explain why a recent study by the Nagy lab using an independent strain of *Lyz2*^Cre^.*Egr2*^fl/fl^ mice concluded that EGR2 is not needed for macrophage differentiation *(29)*. Alternatively, this could reflect that the majority of their studies involved *in vitro* generated macrophages obtained by culturing bone marrow cells with CSF-1 *(29)*. Indeed, we found that *Egr2* deficient monocytes matured into macrophages equally well when cultured *in vitro* with CSF-1. However, despite having been reported to drive expression of EGR family members, including EGR2 *(20)*, in our hands, CSF-1 led to poor upregulation of EGR2 in maturing macrophages *in vitro*. Instead, we identified CSF-2 (GM-CSF) to be a potent inducer of EGR2 expression in maturing macrophages *in vitro*, a finding consistent with the almost unique dependence of alveolar macrophages on alveolar epithelial cell-derived CSF-2, and not CSF-1, for their development and survival *(8–10)*. However, TG-Fβ also induced EGR2 and we confirmed that TGF-β is indispensable for the development of alveolar macrophages *(11)*. Indeed, TGF-β has been shown to induce EGR2 expression outwith the monocyte/macrophage lineage, such as in mammary epithelial cells *(49)*. Exactly how CSF-2 and TGF-β cooperate to promote alveolar macrophage differentiation is incompletely understood, however they both induce expression of PPAR-γ (9, 11) and *Pparg* deficient alveolar macrophages expressed reduced EGR2 *(9)*, suggesting EGR2 lies downstream of PPAR-γ. Notably, alveolar macrophages from mice with genetic ablation of *Pparg, Csf2rb* or *Tgfbr2* (as shown here) have defects in their ability to establish and self-maintain, a phenotype not seen in mice with *Egr2* deficiency. Thus, the EGR2-dependent programme appears to represent a discrete part of alveolar macrophage differentiation, independent of cell survival in health.

EGR2 is consistently referred to as a feature of alternative macrophage activation induced by IL-4. Indeed, addition of IL-4 to *in vitro* macrophage cultures leads to upregulation of EGR2 and deletion of STAT6, a downstream adaptor molecule in the IL-4R signalling cascade, abrogates EGR2 upregulation by IL-4 treated, *in vitro* generated macrophages *(33–35)*. However, mature alveolar macrophages are considered relatively refractory to IL-4 *(50)* and we found no effect of *Il4ra* deficiency on EGR2 expression in health and nor did we detect upregulation of EGR2 by interstitial macrophages which reside in the IL-4/IL-13-rich lung parenchyma during bleomycin-induced fibrosis. Thus, the IL-4–IL-4R axis is sufficient, but not necessary, for inducing EGR2 expression *in vivo*.

Unlike mice deficient in *Pparg, Csf2rb* or *Tgfbr2*, mice with myeloid or macrophage deletion of *Egr2* did not develop spontaneous alveolar proteinosis, suggesting EGR2 does not control the lipid handling and metabolic capacity of alveolar macrophages. However, *Egr2*-deficient mice displayed functional deficiencies in the ability to control low dose *S. pneumoniae* infection, suggesting EGR2 controls the immune protective features of alveolar macrophages. Although we cannot rule out the possibility that this reflects differences in the killing capacity of *Egr2*-deficient alveolar macrophages, genes encoding e.g. reactive oxygen and nitrogen species were unaffected by *Egr2* deficiency. Instead, genes encoding key pathogen recognition receptors and opsonins, were significantly downregulated in the absence of EGR2. These included MARCO and the complement component C3, both of which have been shown to be crucial for the effective elimination of *S. pneumoniae (32, 51).* Indeed, opsonisation is a critical factor in optimizing bacterial clearance by alveolar macrophages in health and disease *(52)*. Thus, EGR2-dependent differentiation equips alveolar macrophages with the machinery to recognise and eliminate pneumococci and this may explain the recurrent pneumonias in individuals with mutations in *EGR2 (26)*. In future work it will be important to determine if this extends to other respiratory pathogens.

Loss of tissue resident macrophages is a common feature of inflammation or tissue injury. We showed that the resident alveolar macrophage population is diminished markedly following administration bleomycin, a chemotherapeutic agent routinely used to generate lung fibrosis. Consistent with previous work *(41, 53)*, we found that the principal mechanism of macrophage replenishment was through recruitment of BM-derived cells which mature into bona fide alveolar macrophages with time. Using *Cx3cr1*-based genetic fate mapping, we also showed that CX3CR1^+^MHCII^+^ cells with a hybrid phenotype could be found in the airways during the fibrotic phase of injury, suggesting that monocyte-derived interstitial macrophages that accumulate following injury may replenish the alveolar macrophage niche. Although we cannot rule out that monocytes enter the airways during this phase to give rise to alveolar macrophages directly, the phenotype of the RFP^+^ transitional cells was more aligned with the phenotype of interstitial macrophages, including high levels of MHCII. Importantly, repopulation of the alveolar macrophage compartment was dependent on EGR2, with constitutive deletion of EGR2 severely blunting the engraftment of monocyte-derived cells into the alveolar macrophage niche. This contrasts with initial population of the developing alveolar niche by foetal liver-derived monocytes, where *Egr2* deficiency does not affect the development of alveolar macrophages. This could indicate differential dependence of developmentally distinct monocytes on EGR2, or the presence of compensatory pathways during development that are not present during repopulation and further work is required to fully understand this.

Interestingly, although previous work has suggested that monocyte-derived alveolar macrophages are key pro-fibrotic cells *(41, 53)*, fibrosis appeared to develop normally in *Egr2* deficient mice, despite the near absence of monocyte-derived alveolar macrophages. The reason for the discrepancy in our findings and those of Misharin *et al. (41)* is unclear, but it could reflect differences in the systems used. For instance, the Misharin study exploited the dependence of alveolar macrophages on Caspase-8 to impede monocyte differentiation into alveolar macrophages by using *Lyz2*^Cre^.*Casp8*^fl/fl^ and *Itgax*^Cre^.*Casp8*^fl/fl^ mice. However, deletion of Caspase-8 also affects the ability of interstitial macrophages to repopulate following depletion, meaning that *Casp8* deficiency may have wider effects on lung macrophage behaviour than disrupting the differentiation of monocyte-derived alveolar macrophages. In contrast, EGR2 expression is restricted to alveolar macrophages and deletion does not affect the reconstitution of the interstitial macrophage compartment. The location of interstitial macrophages in the parenchyma adjacent to fibroblasts and their production of the fibroblast mitogen PDGF-aa, suggests that interstitial macrophages are likely to be key to the fibrotic process *(42)*. Indeed, depletion of interstitial macrophages using *Cx3cr1*^Cre-ERT2^.*Rosa26*^LSL-DTA^ mice reduces lung fibrosis *(42)*, although as we show here, this will also target CX3CR1^+^ cells destined to become monocyte-derived alveolar macrophages. Nevertheless, our data show a clear role for monocyte-derived macrophages in tissue repair processes, as *Lyz2*^Cre^.*Egr2*^fl/fl^ mice failed to repair the lung after injury, a finding consistent with an older study using non-specific, clodronate-mediated depletion of lung macrophages *(54)* and a recent study implicating ApoE-producing, monocyte-derived alveolar macrophages in lung fibrosis resolution *(55)*. These results may help explain the development of restrictive pulmonary disease in individuals with mutations in *EGR2 (26)*.

In summary, our results demonstrate that EGR2 is an evolutionarily conserved transcriptional regulator of alveolar macrophage differentiation, loss of which leads to major phenotypic, transcriptional and functional deficiencies. By identifying EGR2 as a transcriptional regulator, we have begun to dissect how common factors such as CSF2 and TGFβ confer specificity during macrophage differentiation. Importantly, given that recent studies using human systems have proposed that alveolar macrophage maintenance in humans requires monocyte input *(56, 57)*, EGR2 may play a particularly important role in alveolar macrophage differentiation in man. Thus, further work is required to fully understand the molecular pathways downstream of EGR2 and whether this is conserved between mouse and humans, and if EGR2 plays distinct roles in different pathological settings.

## Materials and Methods

### Experimental Animals

Mice were bred and maintained in specific pathogen free (SPF) facilities at the University of Edinburgh or University of Glasgow, UK. All experimental mice were age matched and both sexes were used throughout the study. The mice used in each experiment is documented in the appropriate figure legend. Experiments performed at UK establishments were permitted under licence by the UK Home Office and were approved by the University of Edinburgh Animal Welfare and Ethical Review Body. Genotyping was performed by Transnetyx using real-time PCR.

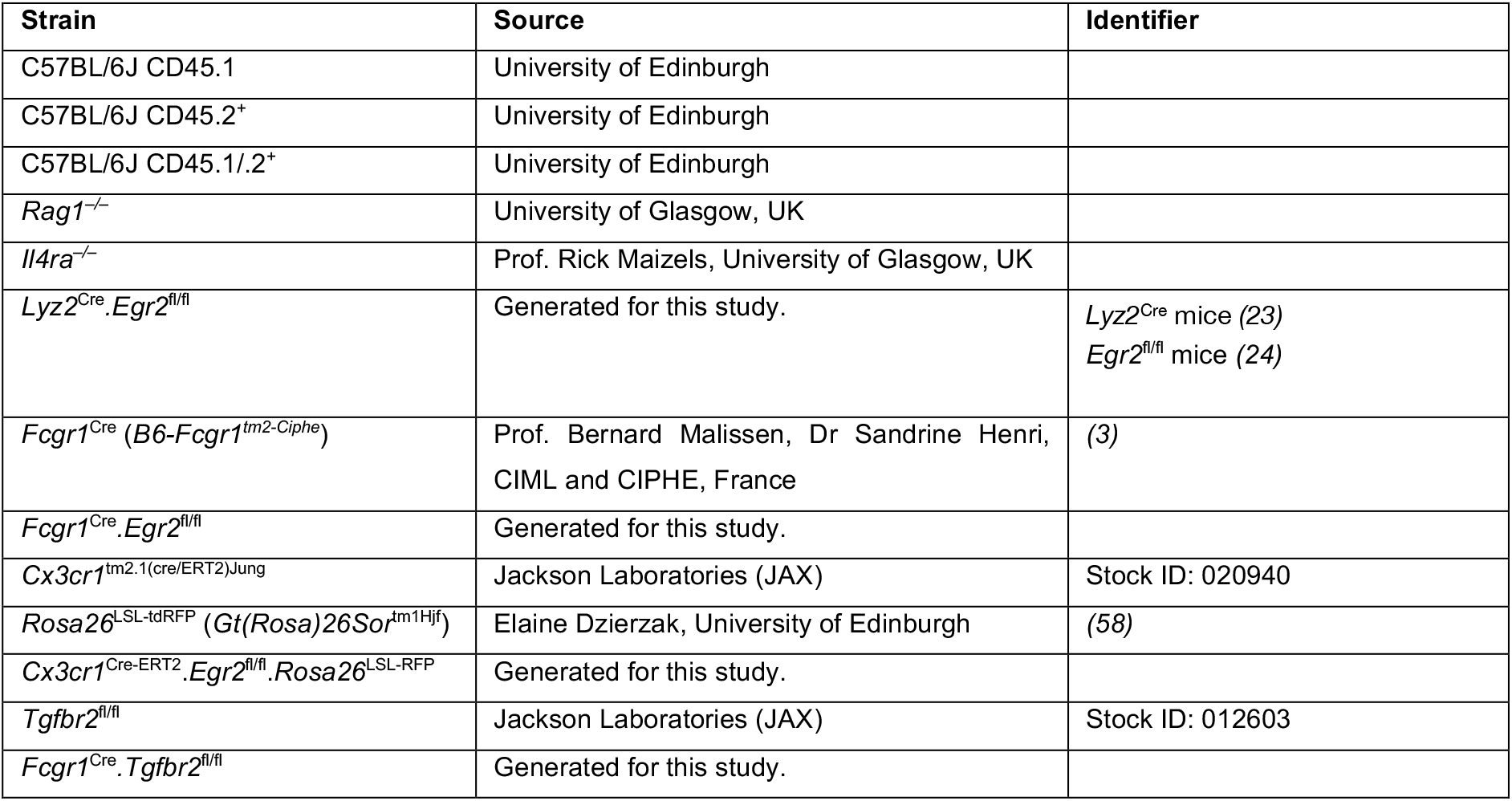

#### Human cells

BAL fluid was obtained from patients attending the Edinburgh Lung Fibrosis Clinic. Ethical permission was granted from the NHS Lothian Research ethics board (LREC 07/S1102/20 06/S0703/53). BAL fluid cells were stained for flow cytometric analysis with antibodies listed in Supplementary Table 4.

#### Tamoxifen-based fate mapping

For induction of Cre activity in *Cx3cr1*^Cre-ERT2/+^ mice, tamoxifen was dissolved in sesame oil overnight at 50mg/ml in a glass vial and administered by oral gavage at 5mg per day for five consecutive days. In bleomycin experiments, tamoxifen was administered from d16 post bleomycin administration for 5 days. Fresh tamoxifen was prepared for each experiment.

#### Bleomycin lung injury

Bleomycin sulphate (Cayman chemicals) was prepared by first dissolving in sterile DMSO (Sigma) and further in sterile PBS at 0.66μg/ml. 8-12-week-old *Lyz2*^Cre^.*Egr2*^fl/fl^ and *Egr2*^fl/fl^ littermate controls were anaesthetised with isofluorane and administered 50μl bleomycin (33ng) or vehicle control (DMSO/PBS) by oropharyngeal aspiration.

#### Streptococcus pneumoniae infection

*Lyz2*^Cre/+^.*Egr2*^fl/fl^ mice and *Egr2*^fl/fl^ littermate control male mice (8–14-week-old) were anaesthetised ketamine/medetomidine and inoculated intratracheally with 50μl of PBS containing 10^4^ CFU *S. pneumoniae* (capsular type 2 strain D39). 100μl of inoculum was plated on blood agar to determine exact dose. Mice were culled 14 h later and BAL fluid collected by lavage performed using sterile PBS. 100μl of lavage fluid was cultured for bacterial growth for 24 h. The remaining lavage fluid was centrifuged at 400g for 5 mins and the resulting cells counted and prepared for flow cytometric analysis.

#### BM chimeric mice

To generate WT:*Lyz2*^Cre^.*Egr2*^fl/fl^ mixed chimeras, CD45.1^+^CD45.2^+^ WT mice were lethally irradiated with two doses of 5 Gy 1 hour apart before being reconstituted immediately WT (CD45.1^+^) and *Lyz2*^Cre/+^.*Egr2*^fl/fl^ or *Egr2*^fl/fl^ (CD45.2^+^) bone marrow at a ratio of 1:1.Chimerism was assessed at 8 weeks after reconstitution.

#### Processing of tissues

Mice were sacrificed by overdose with sodium pentobarbitone followed by exsanguination. Mice were then gently perfused with PBS through the heart. In lung injury/fibrosis experiments, the right lobe was tied off, excised and stored in RPMI with 10% FCS on ice before being prepared for enzymatic digestion (see below). The left lung lobe was inflated with 600μl 4% PFA through an intra-tracheal canula. The trachea was tied off with thread and the lung and heart carefully excised and stored in 4% PFA overnight. Fixed lung tissue was moved to 70% ethanol before being processed for histological assessment. Right lung lobes were chopped finely and digested in pre-warmed RPMI1640 with ‘collagenase cocktail’ (0.625mg ml^−1^ collagenase D (Roche), 0.425mg ml^−1^ collagenase V (Sigma-Aldrich), 1mg ml^−1^ Dispase (ThermoFisher), and 30 U ml^−1^ DNase (Roche Diagnostics GmbH)) for 25 minutes in a shaking incubator at 37°C before being passed through a 100μm strainer followed by centrifugation at 300g for 5 mins. Cell suspensions were then incubated in 2mls Red Blood Cell Lysing Buffer Hybri-Max (Sigma-Aldrich) for 2mins at room temperature to lyse erythrocytes. Cell suspensions were then washed in FACS buffer (2% FCS/2mM EDTA/PBS) followed by centrifugation at 300g for 5 mins. Cells were resuspended in 5mls of FACS buffer, counted and kept on ice until staining for flow cytometry. In some experiments BAL fluid was obtained by lavaging the lungs with 0.8ml DPBS/2mM EDTA via an intra-tracheal catheter. This was repeated three times, with the first wash kept separate for analysis of BAL cytokines, turbidity and protein concentration. To obtain splenic leukocytes, spleens were chopped and digested in HBSS with 1mg/ml collagenase D for 45 mins in a shaking incubator at 37°C before being passed through a 100μm strainer followed by centrifugation at 400g for 5 mins. Tissue preparations were washed in FACS buffer (2% FCS/2mM EDTA/PBS) followed by centrifugation at 300g for 5 mins. Erythrocytes were lysed using red blood cell lysis buffer (Sigma-Aldrich). To obtain liver leukocytes, livers were perfused through the inferior vena cava with sterile PBS and liver tissue excised. Livers were then chopped finely and digested in pre-warmed collagenase ‘cocktail’ (5ml/liver) for 30 minutes in a shaking incubator at 37°C before being passed through an 100μm filter. Cells were washed twice in 50ml ice cold RPMI followed by centrifugation at 300g for 5 mins *(59)*. Supernatants were discarded and erythrocytes were lysed. Epidermal and dermal leukocytes were isolated as described previously *(61)*. To obtain peritoneal leukocytes, the peritoneal cavity was lavaged with RPMI containing 2mM EDTA and 10mM HEPES (both ThermoFisher) as described previously *(62)*. Cells were resuspended in FACS buffer, counted and kept on ice until staining for flow cytometry.

#### Flow cytometry

For analysis of unfixed cells, an equal number of cells were first incubated with 0.025 μg anti-CD16/32 (2.4G2; Biolegend) for 10mins on ice to block Fc receptors and then stained with a combination of the antibodies detailed in Supplementary Table 4. Where appropriate, cells were subsequently stained with streptavidin-conjugated BV650 (Biolegend). Dead cells were excluded using DAPI or 7-AAD (Biolegend) added 2mins before acquisition. When assessing intracellular markers, cells were first washed in PBS and then incubated with Zombie NIR fixable viability dye (Biolegend) for 10mins at room temperature protected from light before following the approach detailed above. Following the final wash step, cells were subsequently fixed and permeabilized using FoxP3/Transcription Factor Staining Buffer Set (eBioscience), and intracellular staining performed using antibodies detailed in Supplementary Table 4. Fluorescence-minus-one (FMO) controls confirmed gating strategies, while discrete populations within lineage^+^ cells were confirmed by omission of the corresponding population-specific antibody. Samples were acquired using a FACS LSRFortessa or AriaII using FACSDiva software (BD) and analyzed with FlowJo software (version 9 or 10; Tree Star). Analysis was performed on single live cells determined using forward scatter height (FCS-H) versus area (FSC-A) and negativity for viability dyes. mRNA was detected by flow cytometry using PrimeFlow technology (ThermoFisher) using probes against Spp1 (AF647) according to the manufacturer’s guidelines. For staining controls in PrimeFlow analysis, the Target Probe Hybridization step was omitted with all other steps identical to samples.

### Transcriptional Analysis

#### qPCR

Real-time PCR assays for the detection of mRNAs were performed using Light Cycler System (Roche) and 384-Well Reaction Plates (Roche). Primer sequences are as follows:

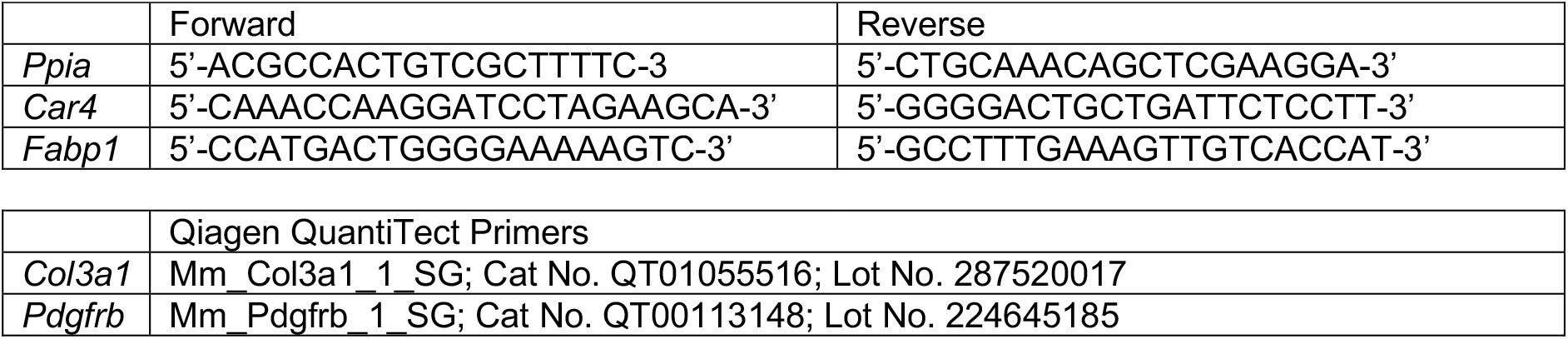

Reactions were performed using SYBR Green System (LightCycler^®^ 480 SYBR Green I Master) according to the manufacturer protocol. 1ul of cDNA (1:50 dilution) were used per sample in a total reaction volume of 10uL. The temperature profile used was as follows: pre-denaturation 5 min at 95°C and then 45 cycles of denaturation for 10s at 95°C, annealing 10s at 60°C, elongation 10s at 72°C. Fluorescence data collection was performed at the end of each elongation step. All samples were tested in duplicates and nuclease free water was used as a non-template control. The relative change was calculated using the 2^−ΔΔCt^ method *(63)*, normalized to *Ppia*.

#### Bulk sequencing

Alveolar macrophages were FACS-purified from lung digests from unmanipulated female *Lyz2*^Cre^.*Egr2*^fl/fl^ mice or *Egr2*^fl/fl^ controls. For each population, 25,000 cells were sorted into 500μl RLT buffer (Qiagen) and snap frozen on dry ice. RNA was isolated using the RNeasy Plus Micro kit (Qiagen). RNA samples were quantified using Qubit 2.0 Fluorometer (Life Technologies, Carlsbad, CA, USA) and RNA integrity was checked with 2100 TapeStation (Agilent Technologies, Palo Alto, CA, USA). SMART-Seq v4 Ultra Low Input Kit for Sequencing was used for full-length cDNA synthesis and amplification (Clontech, Mountain View, CA), and Illumina Nextera XT library was used for sequencing library preparation. Briefly, cDNA was fragmented and adaptor was added using Transposase, followed by limited-cycle PCR to enrich and add index to the cDNA fragments. The final library was assessed with Qubit 2.0 Fluorometer and Agilent TapeStation. The sequencing libraries were multiplexed and clustered on two lanes of a flowcell. After clustering, the flowcell were loaded on the Illumina HiSeq instrument according to manufacturer’s instructions. The samples were sequenced using a 2×150 Paired End (PE) configuration. Image analysis and base calling were conducted by the HiSeq Control Software (HCS) on the HiSeq instrument. Raw sequence data (.bcl files) generated from Illumina HiSeq were be converted into fastq files and de-multiplexed using Illumina bcl2fastq v. 2.17 program. One mis-match was allowed for index sequence identification. After demultiplexing, sequence data was checked for overall quality and yield. Then, sequence reads were trimmed to remove possible adapter sequences and nucleotides with poor quality using Trimmomatic v.0.36. The trimmed reads were mapped to the *Mus musculus* mm10 reference genome available on ENSEMBL using the STAR aligner v.2.5.2b. The STAR aligner uses a splice aligner that detects splice junctions and incorporates them to help align the entire read sequences. BAM files were generated as a result of this step. Unique gene hit counts were calculated by using featureCounts from the Subread package v.1.5.2. Only unique reads that fell within exon regions were counted. After extraction of gene hit counts, the gene hit counts table was used for downstream differential expression analysis. Using DESeq2, a comparison of gene expression between the groups of samples was performed. The Wald test was used to generate p-values and Log2 fold changes. Genes with adjusted p-values < 0.05 and absolute log2 fold changes > 1 were called as differentially expressed genes for each comparison.

#### scRNA-seq

CD11c^+^ and CD11b^+^ cells, excluding Ly6G^+^ and SiglecF^+^CD11b^+^ eosinophils, were FACS-purified from unmanipulated an adult *Rag1*^−/−^ mouse. Single cells were processed through the Chromium Single Cell Platform using the Chromium Single Cell 3’ Library and Gel Bead Kit V2 and the Chromium Single Cell A Chip Kit (both 10X Genomics) as per the manufacturer’s protocol *(64)*. Briefly, single myeloid cells were purified by FACS into PBS/2% FBS, washed twice and cell number measured using a Bio-Rad TC20 Automated Cell Counter (BioRad). Approximately 10,000 cells were loaded to each lane of a 10X chip and partitioned into Gel Beads in Emulsion containing distinct barcodes in the Chromium instrument, where cell lysis and barcoded reverse transcription of RNA occurred, followed by amplification, fragmentation and 5’ adaptor and sample index attachment. Libraries were sequenced on an Illumina HiSeq 4000. For analysis, Illumina BCL sequencing files were demultiplexed using 10x Cell Ranger (version 2.1.1; https://www.10xgenomics.com; ‘cellranger_mkfastq’). Resultant FASTQ files were fed into ‘cellranger_count’ with the transcriptome ‘refdata-cellranger-mm10-1.2.0’ to perform genome alignment, filtering, barcode counting and UMI counting. Downstream QC, clustering and gene expression analysis was performed using the Seurat R package (V3; R version 4.0.2) following the standard pre-processing workflow *(65)*. Cells were filtered on QC covariates used to identify nonviable cells or doublets: number of unique genes per cell (nFeatureRNA >200 & <4000); percentage mitochondrial genes (<20%). Data for resultant 3936 cells were normalized and scaled prior to PCA analysis. Unsupervised clustering based on the first 20 principal components of the most variably expressed genes was performed using a KNN graph-based approach and resultant clusters visualised using the Uniform Manifold Approximation and Projection (UMAP) method. Differential gene expression analysis was used to identify genes expressed by each cell cluster relative to all others, using the nonparametric Wilcoxon rank-sum test and p-value threshold of <0.05. Canonical cell phenotypes were assigned to individual clusters based on the expression of known landmark gene expression profiles.

Publicly available datasets were downloaded from the COVID-19 Cell Atlas *(13–15)* to perform in silico analysis of EGR2 expression in human tissue macrophages. Data were pre-processed and merged using the Seurat R package (V3; R version 4.0.2) following standard methods. Macrophages were extracted based on the expression of C1QA > 0 to compare expression of EGR2 in different human tissue settings.

#### BAL fluid analysis

The first BAL wash was centrifuged at 400g for 5mins and supernatant removed and stored at −80°C until analysis. Total protein concentrations in BAL fluid were measured by BCA Protein Assay according to the manufacturer’s instructions (ThermoFisher). Turbidity was determined following gentle mixing by diluting 25ul of sample with 75ul DPBS and measuring the optical density of 600nm and multiplying by the dilution factor. BAL cytokines were measured using 50ul undiluted sample and the Cytokine & Chemokine 26-Plex ProcartaPlex (Panel 1) assay according to manufacturer’s guidelines (ThermoFisher).

#### Lung histology

Formalin-inflated lungs were fixed overnight in 4% buffered formalin and stored in 70% ethanol. Paraffin-embedded sections of mouse lungs were stained with Masson’s trichome as per the manufacturer’s guidelines.

#### Immunofluorescence imaging

Imagin was performed as described recently (Fercoq et al., 2020). Briefly, samples were permeabilized and blocked for 20min in PBS/Neutral goat serum (NGS) 10%/BSA1%/TritonX-100 (Tx100) 0.3%/Azide 0.05% at 37 °C and stained with 150 μl rabbit anti-CD68 Ab (Polyclonal, ab125212, abcam, 1/200) diluted in PBS/ NGS10%/BSA1%/ TX-100 0.3%/Azide 0.05% for 20min. Samples were washed 3 times with PBS/BSA1%/TX-100 0.1%/Azide 0.05% before adding 150 μl of a solution containing DAPI (1/10000), aSMA-Cy3 (clone 1A4, Sigma, 1/1000), anti-rabbit-AF488 (polyclonal, A-21206 ThermoFisher) diluted in PBS/ NGS10%/BSA1%/ TX-100 0.3%/Azide 0.05% for 1h. Samples were washed 3 times with PBS/BSA1%/TX-100 0.1%/Azide 0.05% and 2 times in PBS. Finally slides were mounted with Vectashield (Vector Laboratories, H-1700). Images were acquired with a Zeiss LSM 880 NLO multiphoton microscope (Carl Zeiss, Oberkochen, Germany) equipped with a 32 channel Gallium arsenide phosphide (GaAsP) spectral detector using 20×/1 NA water immersion objective lens. Samples were excited with a tunable laser (680–1300 nm) set up at 1000 nm and signal was collected onto a linear array of the 32 GaAsp detectors in lambda mode with a resolution of 8.9 nm over the visible spectrum. Spectral images were then unmixed with Zen software (Carl Zeiss) using references spectra acquired from unstained tissues (tissue autofluorescence and second harmonic generation) or beads labelled with Cy3- or AF488-conjugated antibodies.

#### Image analysis

Fluorescence images were analysed with QuPath *(66)*. Full lung section was annotated using the “simple tissue detection” tool and non-pulmonary tissue (trachea, heart tissue) were manually removed from the annotation. In order to refine the analysis, “Pixel classification” was used to segment lung regions of interest. Briefly, software was trained to recognize the different regions using fluorescence (αSMA, SHG and autofluorescence) and texture (all available) features from example images. 2-3 example areas per regions of interest were annotated for each lung to train the pixel classifier. The following regions were analysed: (1) normal lung parenchyma/alveolar tissue, (2) pathologic/fibrotic tissue and (3) collagen rich areas: perivascular/(peri)bronchial spaces + pleura were segmented to avoid false fibrotic region detection. Macrophages were detected using the “Positive cell detection” tool and were expressed as the number DAPI^+^ CD68^+^ cells/mm^2^ of analysed region (full section or regions of interest). Fibrosis was defined as percentage of full section with fibrotic features. All fibrosis scoring and macrophage quantification was performed in a blinded fashion.

#### Statistics

Statistics were performed using Prism 7 (GraphPad Software). The statistical test used in each experiment is detailed in the relevant figure legend.

#### Accession codes

RNA sequencing data that support the findings of this study will be deposited in National Center for Biotechnology Information Gene Expression Omnibus public database (http://www.ncbi.nlm.nih.gov/geo/) upon acceptance.

#### Data availability

Data that support the findings of this study are available from the corresponding authors upon reasonable request.

Further information and requests for resources and reagents should be directed to and will be fulfilled by the Lead Contact, Calum Bain.

## Acknowledgements

We are grateful to Prof. Ping Wang and Dr. Su-Ling Li, QMUL, London for the kind gift of the *Egr2*-floxed strain and to Prof. Rick Maziels, University of Glasgow for the provision of *Il4ra*-deficient mice. Flow cytometry data were generated with support from the QMRI Flow Cytometry and Cell Sorting Facility, University of Edinburgh. mRNA sequencing was performed by Edinburgh Genomics, The University of Edinburgh. Edinburgh Genomics is partly supported through core grants from NERC (R8/H10/56), MRC (MR/K001744/1) and BBSRC (BB/J004243/1). Genewiz performed the bulk RNA-seq analysis. We would like to thank Beth Henderson for technical expertise in setting up 10X sequencing; Dr. Jordan Portman for initial processing of the scRNAseq raw data; the ShIELD (Sheffield, Edinburgh, Newcastle and Birmingham) consortium for access to bacterial stocks; Dr. Duncan Humphries for training and technical assistance with infection studies and Dr. Brian McHugh for advice on bacterial studies. We would also like to thank the Core Services and Advanced Technologies at the Cancer Research UK Beatson Institute, with particular thanks to the Beatson Advanced Imaging Resource (BAIR). Finally, we would like to thank the Bioresearch and Veterinary Services at the University of Edinburgh for husbandry of our mice and other technical assistance. Images from Servier Medical Art (CC BY licence 3 https://creativecommons.org/licenses/by/3.0/) were used and adapted for the generation of some of the graphics.

This research was funded by a Sir Henry Dale Fellowship jointly funded by the Wellcome Trust and the Royal Society [Grant number 206234/Z/17/Z to C.C.B]. CGH is supported by a University of Edinburgh Chancellor’s Fellowship and CGH and RC are supported by Worldwide Cancer Research. The *Lyz2*^*Cre*^.*Egr2*^*fl/fl*^ line was originally generated with funding from the Medical Research Council UK (MR/L008076/1 to S.J.J). GRJ is funded by a Wellcome Trust Clinical Career Development Fellowship (220725/Z/20/Z). SRW is funded by a Wellcome Trust Senior Clinical Fellowship (209220). PTKS received funding from the MRC (MR/N024524/1). FF and LMC are funded by Cancer Research UK core funding (A23983 & A31287), and Breast Cancer Now (2019DecPR1424).

**For the purpose of open access, the author has applied a CC BY public copyright licence to any Author Accepted Manuscript version arising from this submission.**

## Author Contributions

J. McC. performed experiments, analysed the data and edited the manuscript. P.M.K. Performed scRNA-seq analysis. F.F. Designed and performed immunofluorescence analysis of lung tissue. W. T’J. Performed experiments, analysed data and edited the manuscript. C.M.M performed experiments and analysed the data. R. C. and C. G. H provided bioinformatic analysis of ImmGen data. A.S.M. provided advice on and help with the execution of fibrosis experiments. A. H. performed analysis. G.R.J. performed histological analysis of lung sections. S.J.J. generated the *Lyz2*^Cre^.*Egr2*^fl/fl^ mice. N. H. provided access to human bronchoalveolar samples. S.H. and B.M. generated and provided the *Fcgr1*^iCre^ mice. S.R.W. advised on the design of fibrosis experiments and provided reagents for infection experiments. D.D. advised on the design and execution of infection experiments. P.T.K.S. advised and provided infrastructure to perform scRNA-seq analysis. L.M.C. advised and provided infrastructure to perform multi-parameter immunofluorescence analysis. C.C.B. conceived and performed experiments, analysed and interpreted the data, wrote the manuscript, obtained funding and supervised the project.

