## Supplementary Data and Tables for "The transcription factor EGR2 is indispensable for tissue-specific imprinting of alveolar macrophages in health and tissue repair"

***Supplementary Figures***

Supplementary Figure 1: related to Figure 1

Supplementary Figure 2: related to Figure 2

Supplementary Figure 3: related to Figure 2

Supplementary Figure 4: related to Figure 3

Supplementary Figure 5: related to Figure 4 & 5

Supplementary Figure 6: related to Figure 7

Supplementary Figure 7: related to Figure 7

Supplementary Figure 8: related to Figure 8

Supplementary Figure 9: related to Figure 8

***Supplementary Tables***

Supplementary Table 1: Cluster defining genes in scRNA-seq (relates to Figure 1).

Supplementary Table 2: Differentially expressed genes between alveolar macrophages from *Egr2*$^{fl/fl}$ and *Lyz2*$^{Cre/+}$.*Egr2* mice (relates to Figure 3).

Supplementary Table 3: Gene ontology analysis of differentially expressed genes between alveolar macrophages from *Egr2*$^{fl/fl}$ and *Lyz2*$^{Cre/+}$.*Egr2* mice (relates to Figure 3).

Supplementary Table 4: List of antibodies

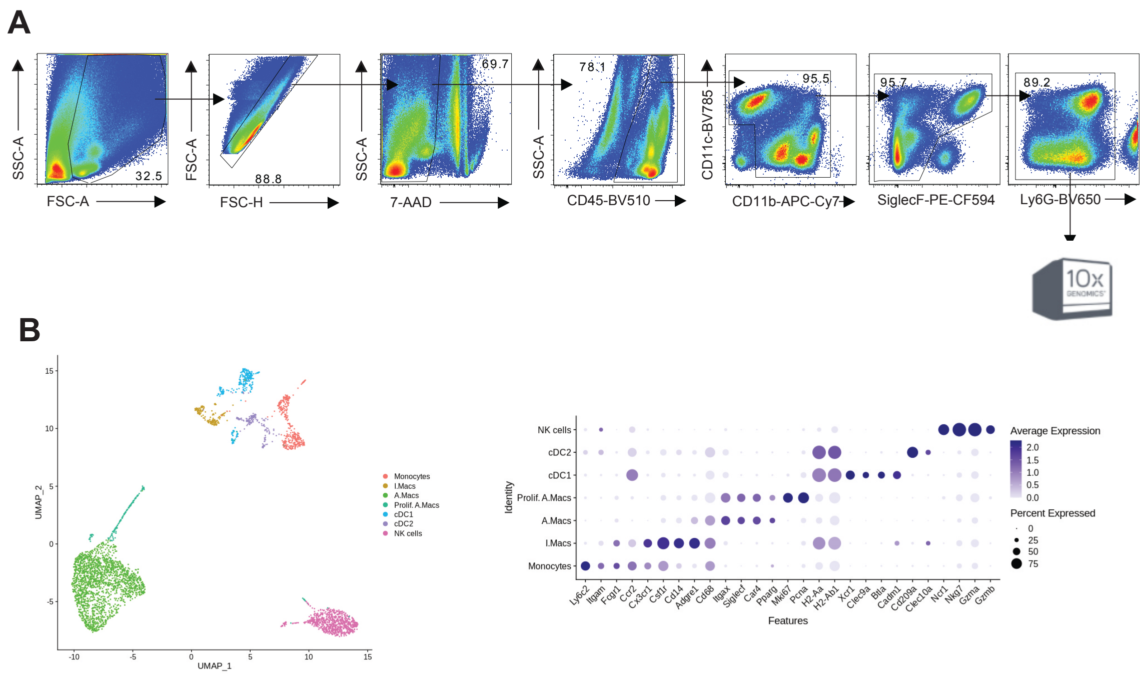

**Supplementary Figure 1**

**A.** Gating strategy used for the purification of mononuclear phagocytes for scRNA-seq analysis using the 10X Genomics platform.

**B.** UMAP dimensionality reduction analysis of non-granulocyte reveals seven clusters of cells in lungs of adult *Rag1*$^{-/-}$ mice (*left*). Right, expression of canonical markers to validate annotations.

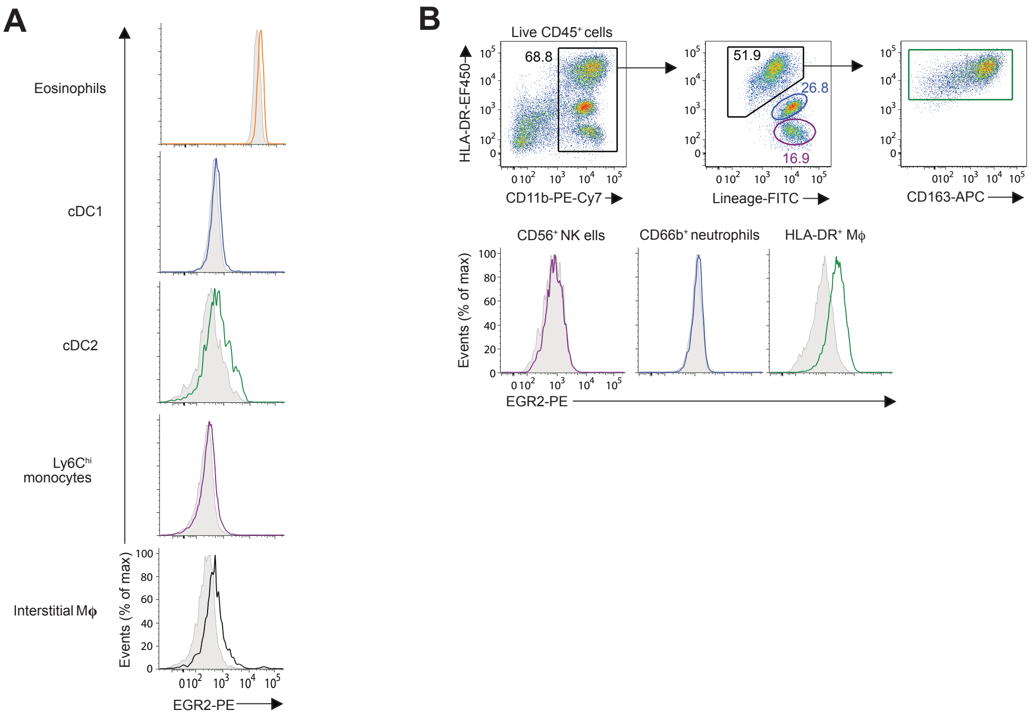

**Supplementary Figure 2**

**A.** Expression of EGR2 by indicated myeloid cells in the lungs of *Egr2*$^{fl/fl}$ (Cre$^{-}$) mice [coloured line] or *Lyz2*$^{Cre/+}$.*Egr2*$^{fl/fl}$ (Cre$^{+}$) mice [shaded histogram]. Data from one of at least three independent experiments with 3-4 mice per group.

**B**. Gating strategy for the identification of alveolar macrophages and granulocytes (lineage$^{+}$) in the BAL fluid from an individual with idiopathic pulmonary fibrosis (IPF) and expression of EGR2 by the indicated populations. Data are representative of two individual patients.

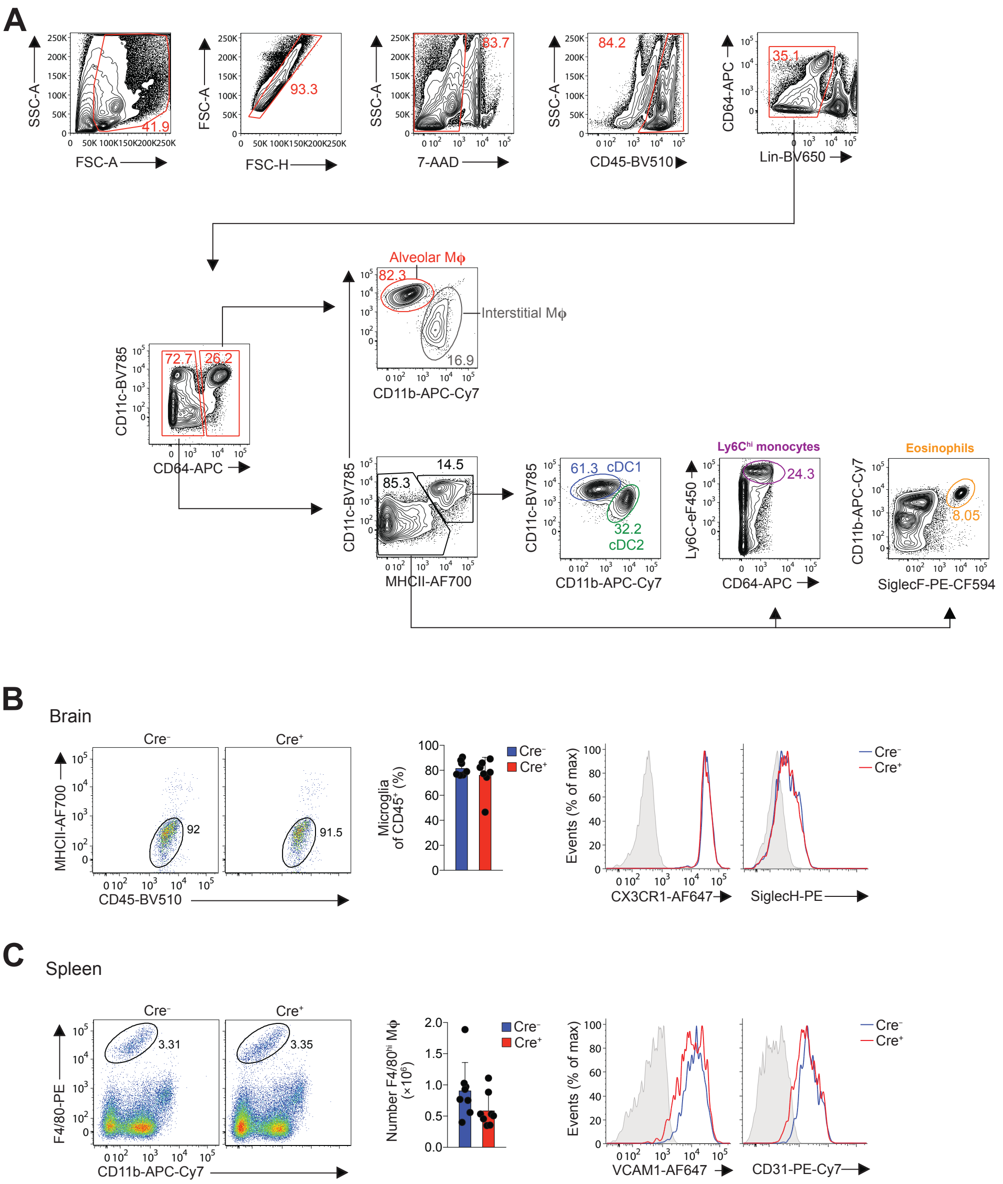

**Supplementary Figure 3**

**A.** Gating strategy for the identification of distinct myeloid cell subsets in the lungs of *Egr2*$^{fl/fl}$ (Cre$^{-}$) or *Lyz2*$^{Cre/+}$.*Egr2*$^{fl/fl}$ (Cre$^{+}$) mice.

**B.** Representative expression of CD45 and MHCII by CD11b$^{+}$Ly6C$^{-}$lineage$^{-}$ cells from brains of adult *Egr2*$^{fl/fl}$ (Cre$^{-}$) or *Lyz2*$^{Cre/+}$.*Egr2*$^{fl/fl}$ (Cre$^{+}$) mice. Graph shows the frequency of CD45$^{lo}$MHCII$^{-}$ microglia of CD45$^{+}$ cells. Symbols represent individual mice and error is S.D. Histograms show representative expression of CX3CR1 and SiglecH by CD45$^{lo}$MHCII$^{-}$ microglia. Shaded histograms represent FMO controls. Data are pooled from three independent experiments with 8 mice per gorup.

**C.** Representative expression of F4/80 and CD11b by lineage$^{-}$ cells from spleens of adult *Egr2*$^{fl/fl}$ (Cre$^{-}$) or *Lyz2*$^{Cre/+}$.*Egr2*$^{fl/fl}$ (Cre$^{+}$) mice. Graph shows the absolute number of F4/80$^{hi}$ macrophages per spleen in each group. Symbols represent individual mice and error is S.D. Histograms show representative expression of VCAM1 and CD31 by F4/80$^{hi}$ macrophages. Shaded histograms represent FMO controls. Data are pooled from three independent experiments with 8 mice per gorup..

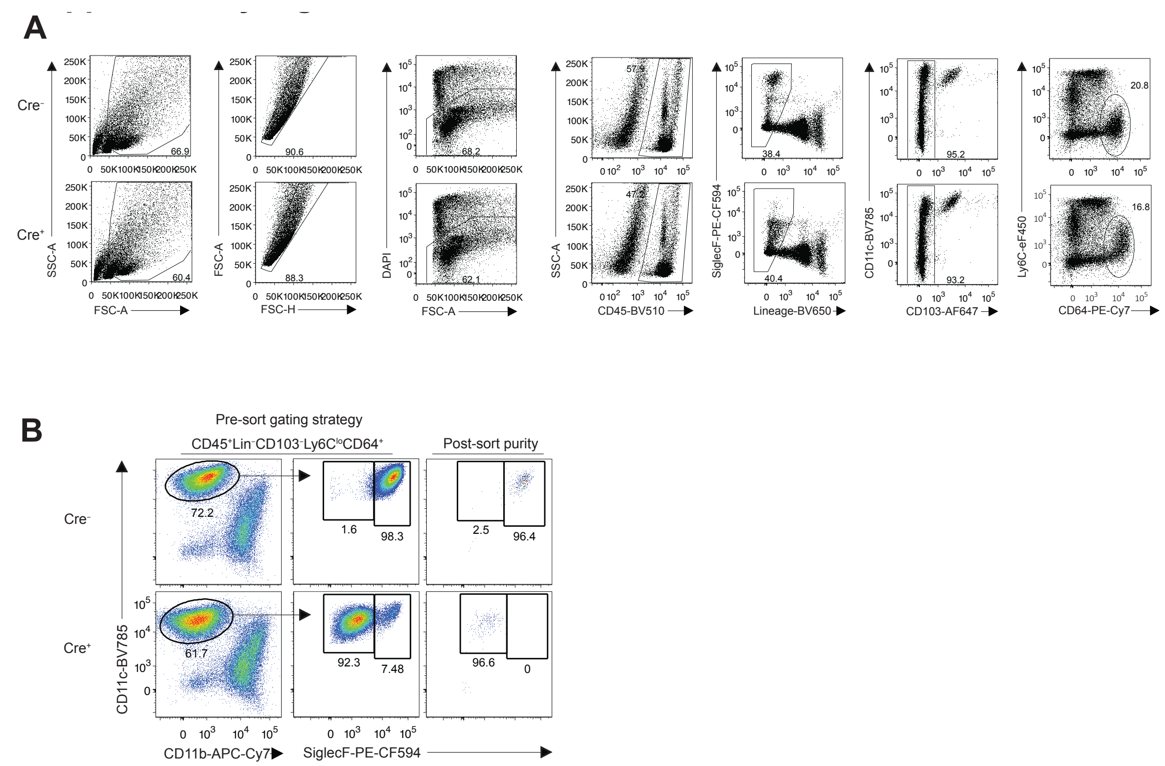

**Supplementary Figure 4**

**A.** Gating strategy for the FACS purification of alveolar macrophages from adult *Egr2*$^{fl/fl}$ (Cre$^{-}$) or *Lyz2*$^{Cre/+}$.*Egr2*$^{fl/fl}$ (Cre$^{+}$) mice for bulk RNA-seq.

**B.** Representative post-sort purity of alveolar macrophages in each group.

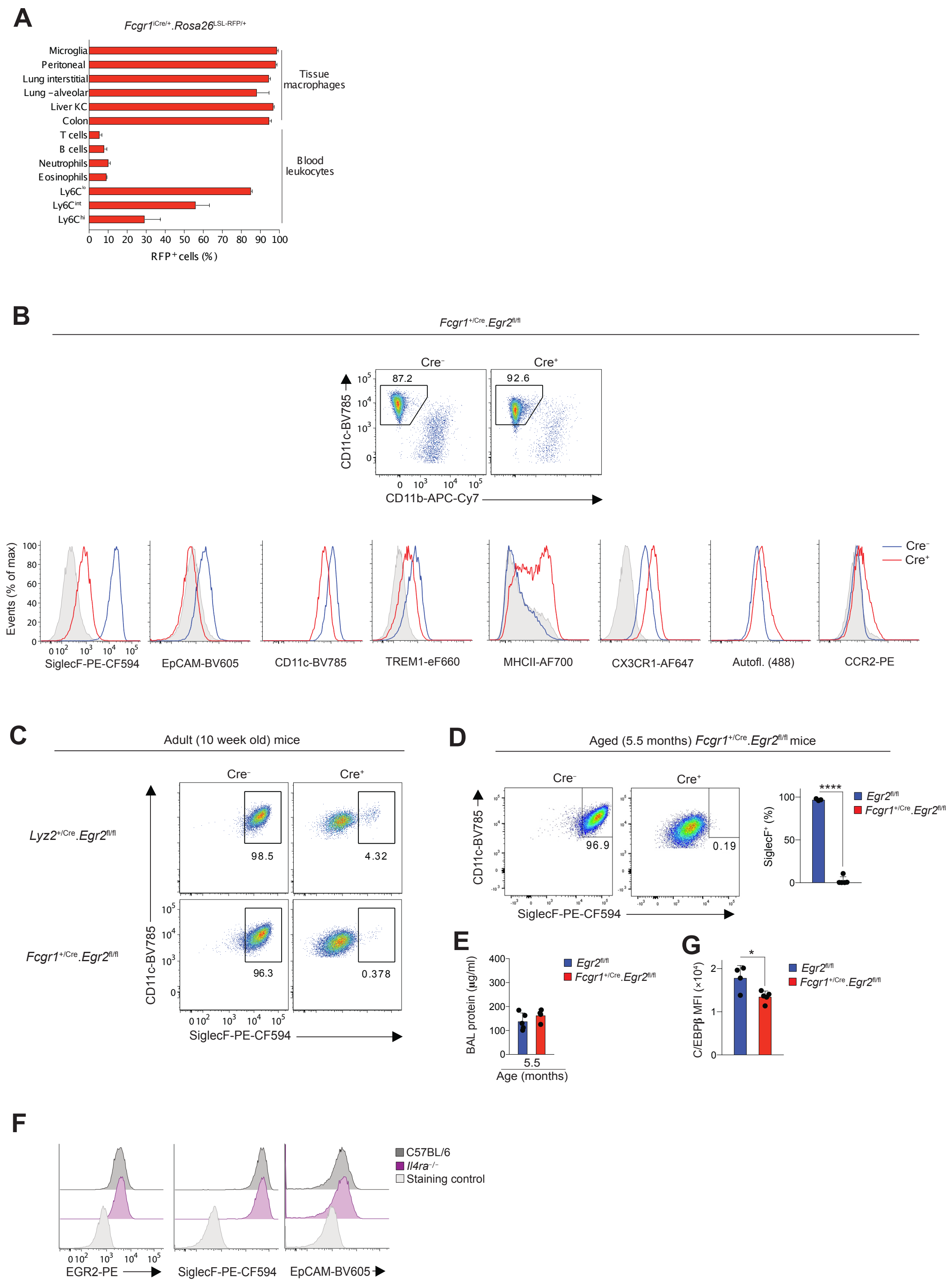

**Supplementary Figure 5**

**A.** Expression of RFP by indicated leukocytes obtained from adult *Fcgr1*$^{\text{iCre/+}}$.*Rosa26*$^{\text{LSL-RFP/+}}$ mice. Data are from 3 mice from one of two independent experiments. Error is S.D.

**B.** Representative expression of CD11c and CD11b by lineage$^-$Ly6C$^-$CD64$^+$ cells amongst lung tissue isolates from *Egr2*$^{\text{fl/fl}}$ mice or *Fcgr1*$^{\text{iCre/+}}$.*Egr2*$^{\text{fl/fl}}$ littermates. Data are from one experiment of two performed with 2 mice per group.

**C.** Representative expression of CD11c and SiglecF by lineage$^-$Ly6C$^-$CD64$^+$CD11c$^+$CD11b$^-$ alveolar macrophages from 8 week old *Egr2*$^{\text{fl/fl}}$ mice or *Lyz2*$^{\text{Cre/+}}$.*Egr2*$^{\text{fl/fl}}$ littermates and an independent colony of *Egr2*$^{fl/fl}$ mice or *Fcgr1*$^{iCre/+}$.*Egr2*$^{fl/fl}$ littermates. Data are from one experiment of two performed with 2 mice per group.

**D.** Representative expression of CD11c and SiglecF by lineage$^{-}$Ly6C$^{-}$CD64$^{+}$CD11c$^{+}$CD11b$^{-}$ alveolar macrophages from 5.5 month old *Egr2*$^{fl/fl}$ mice or *Fcgr1*$^{iCre/+}$.*Egr2*$^{fl/fl}$ littermates. Graph shows the frequency of SiglecF$^{+}$ macrophages of all alveolar macrophages in each group. Data are from one experiment with 3 (Cre$^{-}$) or 5 (Cre$^{+}$) mice per group.

**E**. Protein levels in the BAL fluid of *Egr2*$^{fl/fl}$ mice or *Fcgr1*$^{iCre/+}$.*Egr2*$^{fl/fl}$ littermates at 5.5 months months of age. Symbols represent individual mice and error is s.d.. Data represent 3-4 mice per group from one experiment. Data are from one experiment with 5 (Cre$^{-}$) or 5 (Cre$^{+}$) mice per group.

**F**. Expression of EGR2, SiglecF and EpCAM by alveolar macrophages amongst lung digests from adult C57BL/6 or *Il4ra*$^{-/-}$ mice. Data are from 4 mice per group from one experiment.

**G**. Expression of C/EBP$\beta$ (MFI) by lineage$^{-}$Ly6C$^{-}$CD64$^{+}$CD11c$^{+}$CD11b$^{-}$ alveolar macrophages from 5.5 month old *Egr2*$^{fl/fl}$ mice or *Fcgr1*$^{iCre/+}$.*Egr2*$^{fl/fl}$ littermates. Data are from one experiment with 3 (Cre$^{-}$) or 5 (Cre$^{+}$) mice per group.

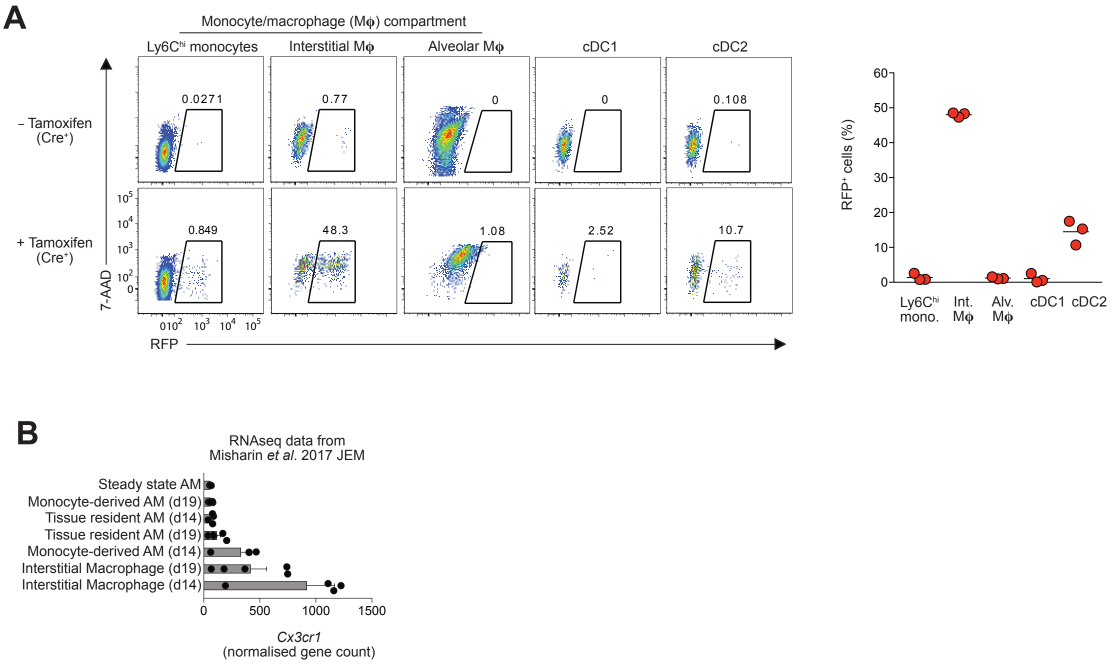

**Supplementary Figure 6**

**A.** Expression of RFP by indicated myeloid cells obtained from adult *Cx3cr1*$^{Cre\text{-}ERT2/+}$.*Rosa26*$^{LSL\text{-}RFP/+}$ mice 24hrs after the final dose of tamoxifen. Mice were administered 5mg tamoxifen by oral gavage for 5 consecutive days. Graph shows the frequency of RFP$^{+}$ amongst each myeloid population. Data are from 1 (Cre$^{-}$) or 3 (Cre$^{+}$) mice from one of three independent experiments performed.

**B.** Expression of *Cx3cr1* by the indicated populations in steady state or the fibrotic phase of bleomycin-induced fibrosis. Data obtained from *(41)* with 4 biological repeats per condition. Error represents SEM.

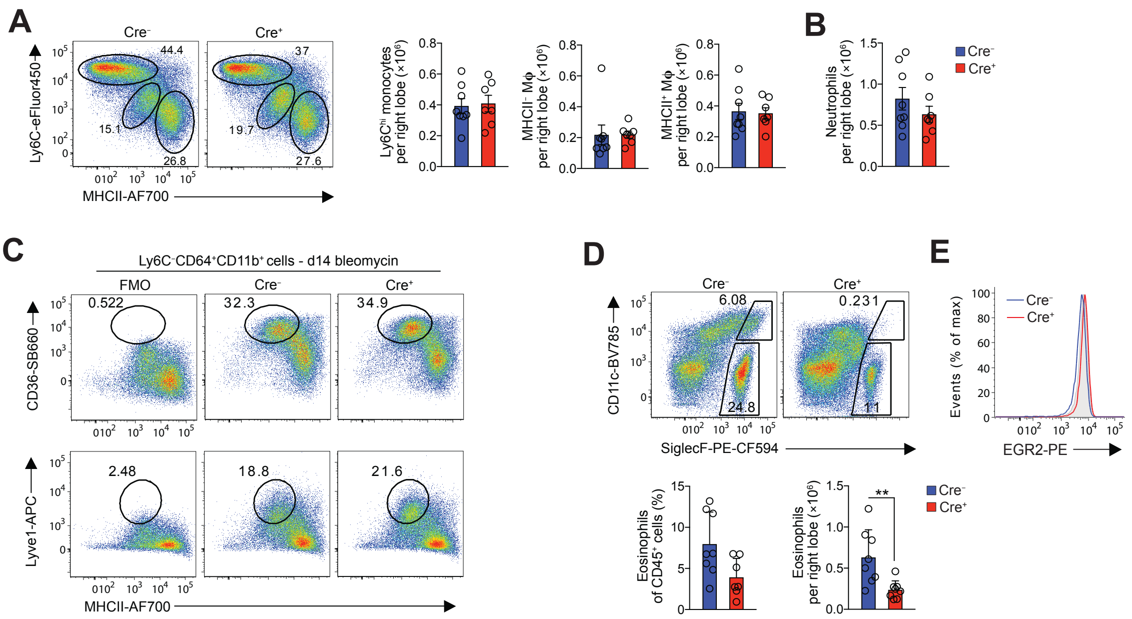

**Supplementary Figure 7**

**A.** Representative expression of Ly6C and MHCII by lineage$^{-}$CD64$^{+}$CD11c$^{var}$CD11b$^{+}$ cells obtained from tissue digests of lungs obtained from adult Cre$^{-}$ (*Egr2*$^{fl/fl}$) mice or Cre$^{+}$ (*Lyz2*$^{Cre/+}$.*Egr2*$^{fl/fl}$) littermates 14 days after the administration of bleomycin. Graphs show the absolute number of Ly6C$^{hi}$ monocytes and MHCII-defined macrophages per right lung lobe. **p<0.01, unpaired Student's *t* test.

**B.** Absolute number of Ly6G$^{+}$ neutrophils per right lung lobe of mice in **A**.

**C.** Representative expression of CD36 and Lyve-1 by Ly6C$^{-}$ interstitial macrophages in lungs of adult *Egr2*$^{fl/fl}$ mice or *Lyz2*$^{Cre/+}$.*Egr2*$^{fl/fl}$ littermates 14 days after the administration of bleomycin compared with fluorescence minus one (FMO) controls.

**D.** Representative expression of CD11c and SiglecF by live CD45$^{+}$ leukocytes obtained from tissue digests of lungs obtained from adult *Egr2*$^{fl/fl}$ mice or *Lyz2*$^{Cre/+}$.*Egr2*$^{fl/fl}$ littermates 14 days after the administration of bleomycin. Graphs show the frequency and absolute number of SiglecF$^{+}$CD11c$^{lo}$ eosinophils per right lung lobe. Mann Whitney test, **p<0.01.

**E.** Representative expression of EGR2 by CD11c$^{lo}$SiglecF$^{+}$ eosinophils 14 days after the administration of bleomycin. Shaded histogram represents staining with isotype control.

Symbols represent individual mice and error is S.D.. Data are pooled from two independent experiments with 8 (*Egr2*$^{fl/fl}$) or 7 (*Lyz2*$^{Cre/+}$.*Egr2*$^{fl/fl}$) mice per group.

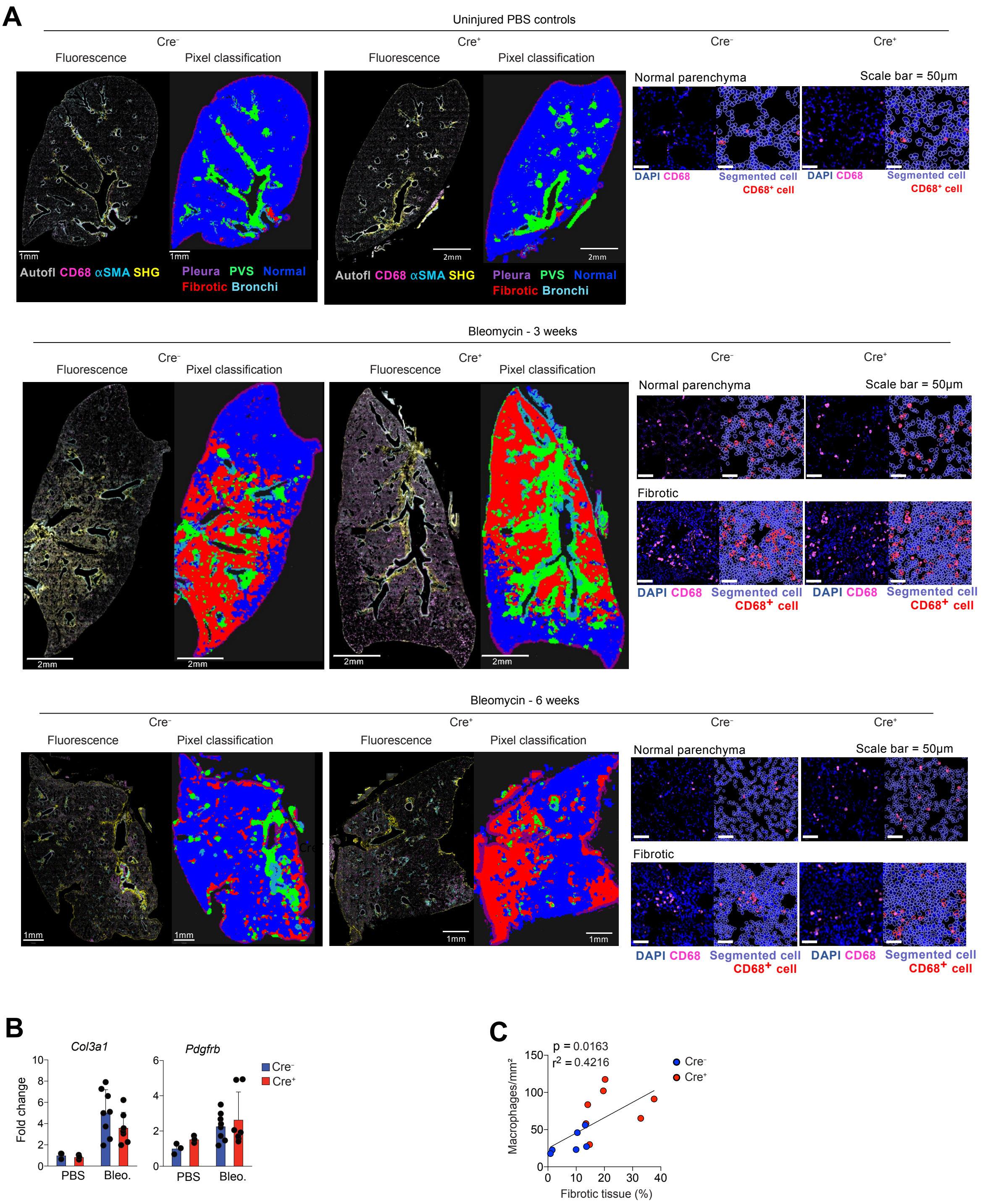

**Supplementary Figure 8**

**A.** 2-photon fluorescence imaging analysis of lung tissue from adult *Egr2*$^{fl/fl}$ mice or *Lyz2*$^{Cre/+}$.*Egr2*$^{fl/fl}$ mice administered PBS (uninjured) or bleomycin 3 or 6 weeks earlier. Sections were stained for CD68, aSMA and DAPI. Autofluorescence is depicted in grey and collagen detected by second harmonic generation (SHG). Pixel classification was used to segment lung regions of interest: (1) normal lung parenchyma/alveolar tissue, (2) pathologic/fibrotic tissue and (3) collagen rich areas (perivascular/bronchial spaces and pleura) were segmented to avoid false fibrotic region detection. *Right,* CD68$^{+}$ macrophage segmentation in lung parenchyma and fibrotic areas.

**B.** Quantitative RT-PCR analysis of *Col3a1* and *Pdgfrb* mRNA in tissue homogenates from lungs of uninjured adult *Egr2*$^{fl/fl}$ mice or *Lyz2*$^{Cre/+}$.*Egr2*$^{fl/fl}$ littermates or mice administered bleomycin 14 days earlier. Symbols represent individual mice and error is S.D. Data are pooled from two independent experiments with 3 (PBS groups), 7 (Cre$^{+}$) or 8 (Cre$^{-}$) mice per bleomycin group.

**C.** Correlation between the number of macrophages (per mm2) in the lung section and lung fibrosis (% of tissue). Linear regression: $R^2$ = 0,4216, p=0.0163. n=7 (Cre$^{+}$) and 8 (Cre$^{-}$) mice.

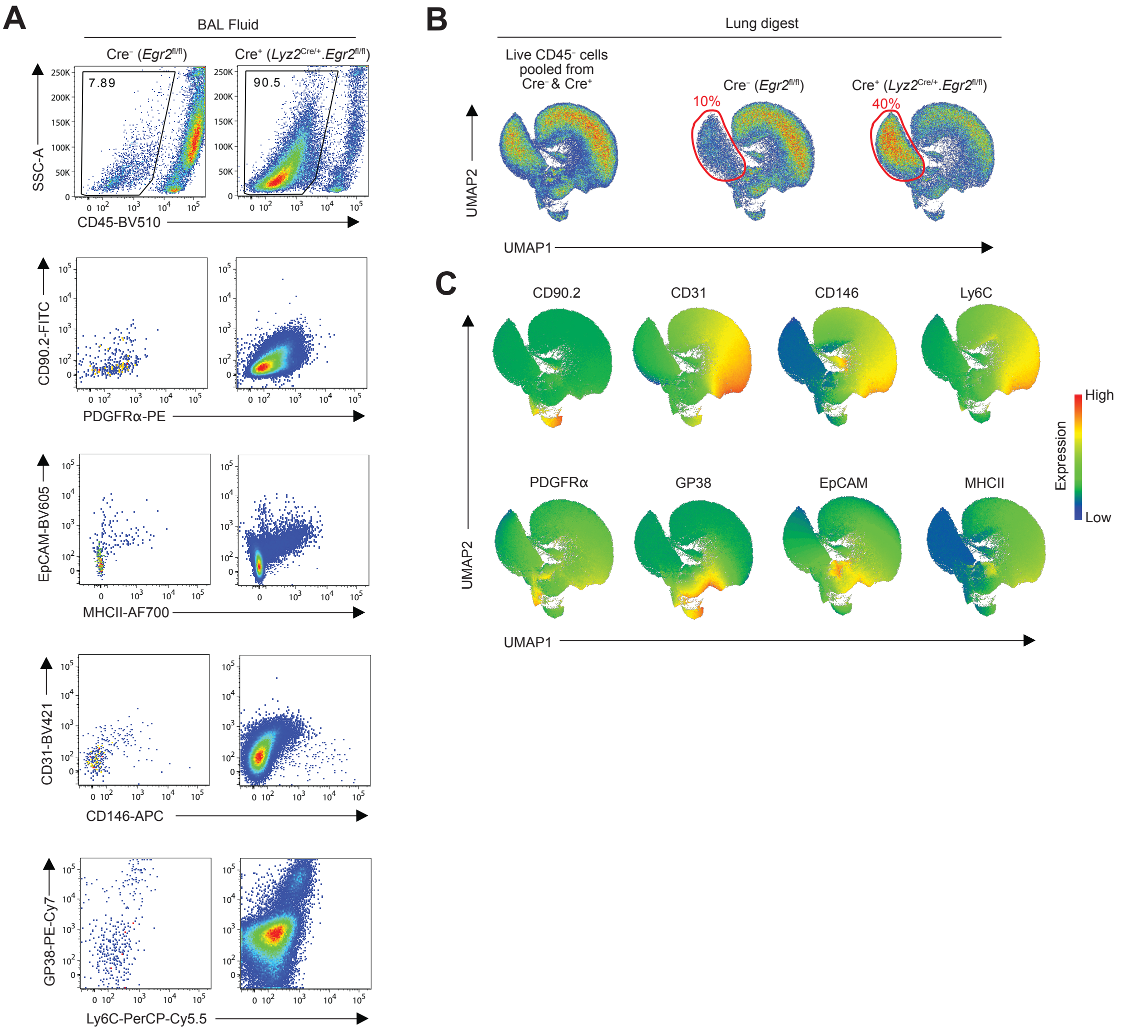

**Supplementary Figure 9**

**A.** Representative expression of CD45, CD90.2, PDGFRα, EpCAM, MHCII, CD31, CD146, GP38 and Ly6C by events present in the BAL fluid of adult Cre$^{-}$ (*Egr2*$^{fl/fl}$) mice or Cre$^{+}$ (*Lyz2*$^{Cre/+}$.*Egr2*$^{fl/fl}$) littermates administered bleomycin 6 weeks earlier.
**B.** UMAP analysis of CD45$^{-}$ cells pooled from adult *Egr2*$^{fl/fl}$ mice or *Lyz2*$^{Cre/+}$.*Egr2*$^{fl/fl}$ littermates 6 weeks after bleomycin-induced injury (*left panel*). Right panel show the contribution of cells deriving from *Egr2*$^{fl/fl}$ mice and *Lyz2*$^{Cre/+}$.*Egr2*$^{fl/fl}$ mice to each cluster.
**C**. Heatmap plots showing the relative expression of the indicated markers by clusters in **B**.

**Supplementary Table 1: Cluster defining genes in scRNAseq**

| Cluster | gene |
|---|---|
| Monocytes | Ifitm6 |
| Monocytes | Plac8 |
| Monocytes | Ms4a4c |
| Monocytes | Ms4a6b |
| Monocytes | Hp |
| Monocytes | Ifitm3 |
| Monocytes | Gm9733 |
| Monocytes | Clec4a1 |
| Monocytes | Ifitm2 |
| Monocytes | Ms4a6c |
| Monocytes | Lst1 |
| Monocytes | Pou2f2 |
| Monocytes | Ly6c2 |
| Monocytes | Ifi27l2a |
| Monocytes | Fyb |
| Monocytes | Cd300a |
| Monocytes | S100a4 |
| Monocytes | Ifi204 |
| Monocytes | B2m |
| Monocytes | Irf7 |
| Monocytes | Gpr141 |
| Monocytes | Klf2 |
| Monocytes | Ly6e |
| Monocytes | Pydc4 |
| Monocytes | F13a1 |
| Monocytes | S100a6 |
| Monocytes | Pyhin1 |
| Monocytes | Adgre5 |
| Monocytes | H2-D1 |
| Monocytes | Nxpe4 |
| Monocytes | Slfn1 |
| Monocytes | Samhd1 |
| Monocytes | Rasgrp2 |
| Monocytes | Rtp4 |
| Monocytes | Gm4955 |
| Monocytes | Serpinb10 |
| Monocytes | AB124611 |
| Monocytes | Coro1a |
| Monocytes | Adgre4 |
| Monocytes | Ly6i |
| Monocytes | H3f3a |
| Monocytes | Sell |
| Monocytes | Limd2 |
| Monocytes | Cx3cr1 |
| Monocytes | Napsa |
| Monocytes | Zbp1 |
| Monocytes | Oasl2 |
| Monocytes | Igsf6 |
| Monocytes | Rps27 |
| Monocytes | Fcer1g |
| Monocytes | Itgam |
| Monocytes | Cd177 |
| Monocytes | Thbs1 |
| Monocytes | Adssl1 |
| Monocytes | Emilin2 |
| Monocytes | Ace |
| Monocytes | Lyl1 |
| Monocytes | Atp1a3 |
| Monocytes | Clec4a3 |
| Monocytes | Isg15 |
| Monocytes | Ccl9 |
| Monocytes | Tyrobp |
| Monocytes | Cd52 |
| Monocytes | Fau |
| Monocytes | Mnda |
| Monocytes | Slc11a1 |
| Monocytes | Hpgd |
| Monocytes | Ccr2 |
| Monocytes | Apoe |
| Monocytes | Ifi47 |
| Monocytes | Ly86 |
| Monocytes | Ceacam1 |
| Monocytes | Pglyrp1 |
| Monocytes | Arpc1b |
| Monocytes | Sorl1 |
| Monocytes | Sp100 |
| Monocytes | Xdh |
| Monocytes | Rpl18a |
| Monocytes | Srgn |
| Monocytes | Aldh2 |
| Monocytes | Eno3 |
| Monocytes | Stxbp6 |
| Monocytes | Emb |
| Monocytes | Fam49a |
| Monocytes | Stat1 |
| Monocytes | Arhgdib |
| Monocytes | Oas1a |
| Monocytes | Susd3 |
| Monocytes | Gm15987 |
| Monocytes | Wfdc17 |
| Monocytes | Nr4a1 |
| Monocytes | Smpdl3a |
| Monocytes | Plbd1 |
| Monocytes | Sema4d |
| Monocytes | Eno1 |
| Monocytes | Ptpro |
| Monocytes | Rap1b |
| Monocytes | Lyz2 |
| Monocytes | Irf5 |
| Monocytes | Csf1r |
| Monocytes | Epsti1 |
| Monocytes | Psmb8 |
| Monocytes | AI839979 |
| Monocytes | Gbp3 |
| Monocytes | Abi3 |
| Monocytes | Itga4 |
| Monocytes | Ldlrad3 |
| Monocytes | BC028528 |
| Monocytes | Nadk |
| Monocytes | Tmsb10 |
| Monocytes | Pla2g7 |
| Monocytes | Xaf1 |
| Monocytes | S1pr5 |
| Monocytes | Trem3 |
| Monocytes | Rpl27a |
| Monocytes | Rplp0 |
| Monocytes | Hes1 |
| Monocytes | Eif3f |
| Monocytes | Itgb7 |
| Monocytes | Btg1 |
| Monocytes | Stap1 |
| Monocytes | Slfn5 |
| Monocytes | Ifngr1 |
| Monocytes | Gsr |
| Monocytes | Mndal |
| Monocytes | Psma7 |
| Monocytes | Crip1 |
| Monocytes | Rpl34 |

**Supplementary Table 1: Cluster defining genes in scRNAseq**

| | |
|---|---|
| Monocytes | Rps14 |
| Monocytes | H2-K1 |
| Monocytes | Rac2 |
| Monocytes | Cyp4f18 |
| Monocytes | Msr1 |
| Monocytes | Cd300e |
| Monocytes | Ifi203 |
| Monocytes | Gltscr2 |
| Monocytes | Treml4 |
| Monocytes | Tmpo |
| Monocytes | Clec2i |
| Monocytes | Pid1 |
| Monocytes | Rps19 |
| Monocytes | Npc2 |
| Monocytes | Clec4e |
| Monocytes | Tmcc1 |
| Monocytes | Cmtm7 |
| Monocytes | Ptpn18 |
| Monocytes | Lrrc25 |
| Monocytes | Sik1 |
| Monocytes | Spn |
| Monocytes | Pkm |
| Monocytes | Tap1 |
| Monocytes | Rps27a |
| Monocytes | Ethe1 |
| Monocytes | Arl5c |
| Monocytes | Acer3 |
| Monocytes | Rpl19 |
| Monocytes | Rnf213 |
| Monocytes | Ccnd3 |
| Monocytes | Bin2 |
| Monocytes | Tspan13 |
| Monocytes | Lyn |
| Monocytes | Nupr1 |
| Monocytes | Ap1s2 |
| Monocytes | Tsc22d3 |
| Monocytes | Ms4a4b |
| Monocytes | Rpl13 |
| Monocytes | Irgm1 |
| Monocytes | Sat1 |
| Monocytes | Gngt2 |
| Monocytes | Rps23 |
| Monocytes | Pld4 |
| Monocytes | Prdx5 |
| Monocytes | Metrnl |
| Monocytes | Hck |
| Monocytes | Gm9843 |
| Monocytes | Ubb |
| Monocytes | Sap30 |
| Monocytes | Klra2 |
| Monocytes | Tpd52 |
| Monocytes | Tln1 |
| Monocytes | Rps6 |
| Monocytes | Rps29 |
| Monocytes | Phf11b |
| Monocytes | Rap1a |
| Monocytes | Alox5ap |
| Monocytes | Rgs2 |
| Monocytes | Rpl6 |
| Monocytes | Camkk2 |
| Monocytes | Lpar6 |
| Monocytes | Gpx1 |
| Monocytes | Samsn1 |
| Monocytes | Phf11d |
| Monocytes | Ncf4 |
| Monocytes | Actr3 |
| Monocytes | Clec12a |
| Monocytes | Rps27rt |
| Monocytes | Igtp |
| Monocytes | Slfn2 |
| Monocytes | Rps16 |
| Monocytes | Rpl24 |
| Monocytes | Irf1 |
| Monocytes | Rpl18 |
| Monocytes | Fam49b |
| Monocytes | Eif4e3 |
| Monocytes | Atf3 |
| Monocytes | Cebpb |
| Monocytes | Sirpb1a |
| Monocytes | Rpl9 |
| Monocytes | Klf3 |
| Monocytes | Prkcb |
| Monocytes | Lsp1 |
| Monocytes | Bst2 |
| Monocytes | Vsir |
| Monocytes | Tspo |
| Monocytes | Lbr |
| Monocytes | Gpr35 |
| Monocytes | Mef2c |
| Monocytes | Cd53 |
| Monocytes | Calhm2 |
| Monocytes | Gbp2 |
| Monocytes | Rnase6 |
| Monocytes | Itm2b |
| Monocytes | Prdx6 |
| Monocytes | Sp110 |
| Monocytes | Klf13 |
| Monocytes | Arl6ip5 |
| Monocytes | Tmem51 |
| Monocytes | Hspa1a |
| Monocytes | Cd48 |
| Monocytes | Ptprj |
| Monocytes | Tiam1 |
| Monocytes | Emp3 |
| Monocytes | Zeb2 |
| Monocytes | Mtus1 |
| Monocytes | Slc16a3 |
| Monocytes | Rpl4 |
| Monocytes | Tifa |
| Monocytes | Nfkbiz |
| Monocytes | Pstpip1 |
| Monocytes | Rpl6l |
| Monocytes | H2-T23 |
| Monocytes | Rasa3 |
| Monocytes | Apobec1 |
| Monocytes | Lpcat2 |
| Monocytes | Sirpb1b |
| Monocytes | Nrros |
| Monocytes | Mettl9 |
| Monocytes | Tnfrsf1a |
| Monocytes | Ctsb |
| Monocytes | Prkcd |
| Monocytes | Atp2b1 |
| Monocytes | Msn |
| Monocytes | Ypel3 |
| Monocytes | Capzb |
| Monocytes | Ptprc |
| Monocytes | Plxnb2 |
| Monocytes | Msrb1 |

**Supplementary Table 1: Cluster defining genes in scRNAseq**

| | | | | | |
|---|---|---|---|---|---|
| Monocytes | Rps11 | Monocytes | Marcks | Monocytes | 1110008F13Rik |
| Monocytes | Arhgap15 | Monocytes | Csgalnact2 | Monocytes | Fam46a |
| Monocytes | H2-T22 | Monocytes | Fcgr1 | Monocytes | Zfand5 |
| Monocytes | Csf3r | Monocytes | Skint3 | Monocytes | Gmfg |
| Monocytes | Cnn2 | Monocytes | Rasgrp4 | Monocytes | Plekhf2 |
| Monocytes | Fam111a | Monocytes | Parp14 | Monocytes | Dusp1 |
| Monocytes | Fcgr3 | Monocytes | Ifnar2 | Monocytes | Zyx |
| Monocytes | Gsdmd | Monocytes | Il10ra | Monocytes | Lbh |
| Monocytes | Kdm7a | Monocytes | Selplg | Monocytes | Il6ra |
| Monocytes | Sub1 | Monocytes | Capza2 | Monocytes | Oaz2 |
| Monocytes | Pilrb1 | Monocytes | Arpc2 | Monocytes | Tpm4 |
| Monocytes | H2-Q7 | Monocytes | Cox7a2l | Monocytes | Adrbk2 |
| Monocytes | Gm8730 | Monocytes | Psme2 | Monocytes | Cdkn2d |
| Monocytes | Pim1 | Monocytes | C1galt1c1 | Monocytes | Ltb4r1 |
| Monocytes | St8sia4 | Monocytes | Cybb | Monocytes | Supt4a |
| Monocytes | Pot1b | Monocytes | Rps13 | Monocytes | Dusp16 |
| Monocytes | Tnfrsf1b | Monocytes | Cbfa2t3 | Monocytes | Chst12 |
| Monocytes | Eif3h | Monocytes | Add3 | Monocytes | H2-Q4 |
| Monocytes | Psmb9 | Monocytes | Hmha1 | Monocytes | Kras |
| Monocytes | Ptpn6 | Monocytes | Il1b | Monocytes | Cd244 |
| Monocytes | Slc44a2 | Monocytes | Trim30a | Monocytes | Ptpn1 |
| Monocytes | Rpl17 | Monocytes | Ogfrl1 | Monocytes | Tpm3 |
| Monocytes | Ddit4 | Monocytes | Rpl3 | Monocytes | Smpdl3b |
| Monocytes | Pycard | Monocytes | Rps9 | Monocytes | Rpl11 |
| Monocytes | Junb | Monocytes | Sirpb1c | Monocytes | Mrpl33 |
| Monocytes | Neurl3 | Monocytes | Ninj1 | Monocytes | Casp1 |
| Monocytes | Ccdc109b | Monocytes | Rpl8 | Monocytes | Tifab |
| Monocytes | Rps15a | Monocytes | Psme1 | Monocytes | Serinc3 |
| Monocytes | Rpl36 | Monocytes | H3f3b | Monocytes | Rpl7 |
| Monocytes | Cdk2ap2 | Monocytes | Dbnl | Monocytes | Syf2 |
| Monocytes | Eif4ebp1 | Monocytes | Rnf114 | Monocytes | Gm2a |
| Monocytes | Rpl37 | Monocytes | Rnf115 | Monocytes | Lims1 |
| Monocytes | Hfe | Monocytes | Rnaset2a | Monocytes | Rpl26 |
| Monocytes | Bcl10 | Monocytes | Rpl21 | Monocytes | F630028O10Rik |
| Monocytes | Il17ra | Monocytes | Nmi | Monocytes | Trps1 |
| Monocytes | Ube2l6 | Monocytes | Myo1g | Monocytes | Ier2 |
| Monocytes | Ifngr2 | Monocytes | Fgl2 | Monocytes | Gm9844 |
| Monocytes | Ikbkb | Monocytes | Rps7 | Monocytes | Rsrp1 |
| Monocytes | Ssh2 | Monocytes | Hgsnat | Monocytes | Card19 |
| Monocytes | Slc25a5 | Monocytes | Spi1 | Monocytes | Uba52 |
| Monocytes | Eif3e | Monocytes | Zeb2os | Monocytes | Arpc3 |
| Monocytes | H2afj | Monocytes | Pmaip1 | Monocytes | Pomp |
| Monocytes | Irf9 | Monocytes | Ccdc12 | Monocytes | Tmem243 |

**Supplementary Table 1: Cluster defining genes in scRNAseq**

| | | | | | |
|---|---|---|---|---|---|
| Monocytes | Ehd4 | Monocytes | Mar-01 | Monocytes | Ogfr |
| Monocytes | Ppp1ca | Monocytes | Ppdpf | Monocytes | Nfil3 |
| Monocytes | Fxyd5 | Monocytes | H2afy | Monocytes | Milr1 |
| Monocytes | Zfos1 | Monocytes | Fgr | Monocytes | Snrpb |
| Monocytes | Unc93b1 | Monocytes | Tuba1a | Monocytes | Scpep1 |
| Monocytes | D1Ertd622e | Monocytes | Gnas | Monocytes | Mef2a |
| Monocytes | Cd300ld | Monocytes | Ctsh | Monocytes | Tsc22d4 |
| Monocytes | Nsa2 | Monocytes | Cnpy3 | Monocytes | Ahcyl2 |
| Monocytes | Btg2 | Monocytes | Prr13 | Monocytes | Rnaset2b |
| Monocytes | Arl4c | Monocytes | Psmb10 | Monocytes | Tbpl1 |
| Monocytes | Rpl10 | Monocytes | Smchd1 | Monocytes | Arid4a |
| Monocytes | Thbd | Monocytes | Myo1f | Monocytes | Tor3a |
| Monocytes | Clic1 | Monocytes | Ndufa6 | Monocytes | Pnp |
| Monocytes | Trim30d | Monocytes | Rnf130 | Monocytes | Anp32a |
| Monocytes | Eif3k | Monocytes | St3gal4 | Monocytes | Tmem50b |
| Monocytes | Degs1 | Monocytes | Rpl15 | Monocytes | Fam32a |
| Monocytes | Foxp1 | Monocytes | AI607873 | Monocytes | Evi2a |
| Monocytes | Gm11808 | Monocytes | Glipr1 | Monocytes | Ppfia4 |
| Monocytes | Tpst2 | Monocytes | Ctage5 | Monocytes | Filip1l |
| Monocytes | Spop | Monocytes | Shisa5 | Monocytes | Tab2 |
| Monocytes | Mbp | Monocytes | Arhgef1 | Monocytes | Stk38 |
| Monocytes | Shfm1 | Monocytes | Sp140 | Monocytes | Aip |
| Monocytes | Fis1 | Monocytes | Tmem50a | Monocytes | Rara |
| Monocytes | Lgals9 | Monocytes | Stx16 | Monocytes | Ptpre |
| Monocytes | Rras | Monocytes | Sh3kbp1 | Monocytes | Fcgr4 |
| Monocytes | Arpc4 | Monocytes | Dok1 | Monocytes | Arap1 |
| Monocytes | Adipor1 | Monocytes | Samd9l | Monocytes | Tmem59 |
| Monocytes | Relt | Monocytes | Trafd1 | Monocytes | Use1 |
| Monocytes | Ncf2 | Monocytes | Lmo2 | Monocytes | Ubac2 |
| Monocytes | Stk24 | Monocytes | Rpl29 | Monocytes | Tmem38b |
| Monocytes | Casp4 | Monocytes | Ppp1r15a | Monocytes | Necap2 |
| Monocytes | Irf2 | Monocytes | Btf3 | Monocytes | Rpl9-ps6 |
| Monocytes | 9930111J21Rik2 | Monocytes | Prkx | Monocytes | Sepw1 |
| Monocytes | Il2rg | Monocytes | Rnf149 | Monocytes | Rbfa |
| Monocytes | Pilrb2 | Monocytes | Polb | Monocytes | Ifi35 |
| Monocytes | Gnb2 | Monocytes | Dok3 | Monocytes | G6pdx |
| Monocytes | Fam174a | Monocytes | Wasf2 | Monocytes | Whsc1l1 |
| Monocytes | Eif1 | Monocytes | Rhoa | Monocytes | Fos |
| Monocytes | Ppp1r12a | Monocytes | Yipf1 | Monocytes | 2310001H17Rik |
| Monocytes | Hacd4 | Monocytes | Lat2 | Monocytes | Sumo1 |
| Monocytes | Pirb | Monocytes | Rpl36al | Monocytes | Gstp1 |
| Monocytes | Htra2 | Monocytes | Tnfrsf13b | Monocytes | Stk10 |
| Monocytes | Tmem55b | Monocytes | Anxa6 | Monocytes | Tpgs1 |

**Supplementary Table 1: Cluster defining genes in scRNAseq**

| | | | | | |
|---|---|---|---|---|---|
| Monocytes | Fam107b | Monocytes | Fgd4 | Monocytes | Vapa |
| Monocytes | Tlr7 | Monocytes | Tax1bp1 | Monocytes | Chmp2b |
| Monocytes | Maf1 | Monocytes | Birc3 | Monocytes | Glipr2 |
| Monocytes | Ppp1cb | Monocytes | Herc4 | Monocytes | Nipbl |
| Monocytes | Snx20 | Monocytes | Surf1 | Monocytes | Cope |
| Monocytes | Pdcd4 | Monocytes | Zc3hav1 | Monocytes | Ube2d2a |
| Monocytes | Cxcr4 | Monocytes | Fmnl1 | Monocytes | Pnrc2 |
| Monocytes | Dtx3l | Monocytes | Kdm6b | Monocytes | Was |
| Monocytes | Rel | Monocytes | Pip4k2a | Monocytes | Mcmbp |
| Monocytes | Zfp36 | Monocytes | Bach1 | Monocytes | Eif4h |
| Monocytes | 4930523C07Rik | Monocytes | Cmpk1 | Monocytes | Eif3m |
| Monocytes | Zfp706 | Monocytes | Gch1 | Monocytes | Sephs2 |
| Monocytes | Tnfaip8l2 | Monocytes | Cers6 | Monocytes | Snw1 |
| Monocytes | Tcof1 | Monocytes | Calm1 | Monocytes | Oser1 |
| Monocytes | Spg21 | Monocytes | Plekho1 | Monocytes | Mapkapk2 |
| Monocytes | Cdc42se1 | Monocytes | Dusp6 | Monocytes | Ccdc50 |
| Monocytes | Sgms1 | Monocytes | Nin | Monocytes | Map2k1 |
| Monocytes | Jarid2 | Monocytes | Fcgr2b | Monocytes | Klf4 |
| Monocytes | Fn1 | Monocytes | Trim12a | Monocytes | Dpm1 |
| Monocytes | Bcap31 | Monocytes | Gm5150 | Monocytes | Pxn |
| Monocytes | Lilrb4a | Monocytes | Atp5c1 | Monocytes | Brd7 |
| Monocytes | Cldnd1 | Monocytes | H13 | Monocytes | Max |
| Monocytes | Csk | Monocytes | Vdac3 | Monocytes | Il10rb |
| Monocytes | Hps3 | Monocytes | Ahnak | Monocytes | Cct5 |
| Monocytes | Aif1 | Monocytes | Nuak2 | Monocytes | Stk17b |
| Monocytes | Nsmce4a | Monocytes | Vbp1 | Monocytes | Ccm2 |
| Monocytes | Psme2b | Monocytes | N4bp2l1 | Monocytes | Clint1 |
| Monocytes | Atp1a1 | Monocytes | Bri3bp | Monocytes | Macf1 |
| Monocytes | H2-M3 | Monocytes | Vmp1 | Monocytes | Rab8a |
| Monocytes | Ube2a | Monocytes | Dck | Monocytes | Dnajb1 |
| Monocytes | Stard3 | Monocytes | Ndel1 | Monocytes | Taf6l |
| Monocytes | Ms4a6d | Monocytes | Nab1 | Monocytes | Rpl10-ps3 |
| Monocytes | Map3k1 | Monocytes | Ly96 | Monocytes | Sgk3 |
| Monocytes | Fbxw2 | Monocytes | Mrpl30 | Monocytes | Aup1 |
| Monocytes | Rchy1 | Monocytes | Rnf166 | Monocytes | Polr2l |
| Monocytes | Scarb1 | Monocytes | Hmox2 | Monocytes | Rpp21 |
| Monocytes | Itgb2 | Monocytes | Ppp2r5a | Monocytes | Wsb1 |
| Monocytes | Gm26740 | Monocytes | Mkrn1 | Monocytes | Lgals3bp |
| Monocytes | Psmd8 | Monocytes | Mrpl52 | Monocytes | Plcg2 |
| Monocytes | Itch | Monocytes | Cmtm6 | Monocytes | Dnaja1 |
| Monocytes | Tmem234 | Monocytes | Mgst1 | Monocytes | Ddx24 |
| Monocytes | Nfam1 | Monocytes | Agpat4 | Monocytes | Cript |
| Monocytes | Esd | Monocytes | Gm9493 | Monocytes | Glud1 |

**Supplementary Table 1: Cluster defining genes in scRNAseq**

| | |
|---|---|
| Monocytes | B4galt5 |
| Monocytes | Ogt |
| Monocytes | Efhd2 |
| Monocytes | Tnpo3 |
| Monocytes | Cdkn1b |
| Monocytes | Jak1 |
| Monocytes | Cd2ap |
| Monocytes | Bak1 |
| Monocytes | Cebpz |
| Monocytes | Chrac1 |
| Monocytes | Tppp3 |
| Monocytes | Notch2 |
| Monocytes | Ppm1h |
| Monocytes | Tpr |
| Monocytes | Il13ra1 |
| Monocytes | Emd |
| Monocytes | Cks2 |
| Monocytes | Kpna4 |
| Monocytes | Cd37 |
| Monocytes | Frg1 |
| Monocytes | Dram2 |
| Monocytes | Sirt7 |
| Monocytes | Gpbp1 |
| Monocytes | Nfkb1 |
| Monocytes | Mkln1 |
| Monocytes | Crlf2 |
| Monocytes | Pdcd10 |
| Monocytes | Itgal |
| I.Macs | C1qc |
| I.Macs | Stab1 |
| I.Macs | Pf4 |
| I.Macs | Tmem176a |
| I.Macs | Tmem176b |
| I.Macs | C3ar1 |
| I.Macs | Ms4a7 |
| I.Macs | Maf |
| I.Macs | C1qa |
| I.Macs | C1qb |
| I.Macs | Gas6 |
| I.Macs | Cxcl16 |
| I.Macs | Mafb |
| I.Macs | Igfbp4 |
| I.Macs | Folr2 |
| I.Macs | Cd63 |
| I.Macs | Cacnb3 |
| I.Macs | Fcrls |
| I.Macs | Vcam1 |
| I.Macs | Cd163 |
| I.Macs | Marcks |
| I.Macs | Zmynd15 |
| I.Macs | Fxyd2 |
| I.Macs | Igf1 |
| I.Macs | Col14a1 |
| I.Macs | Ccr7 |
| I.Macs | Ccl12 |
| I.Macs | Fscn1 |
| I.Macs | Timp2 |
| I.Macs | Tspan3 |
| I.Macs | Cbr2 |
| I.Macs | Marcksl1 |
| I.Macs | Sdc4 |
| I.Macs | Aif1 |
| I.Macs | Hpgd |
| I.Macs | Pmp22 |
| I.Macs | Tmem37 |
| I.Macs | Apoe |
| I.Macs | Hpgds |
| I.Macs | Adam19 |
| I.Macs | Cfh |
| I.Macs | Zfp36l1 |
| I.Macs | Cd72 |
| I.Macs | Lgmn |
| I.Macs | Mgl2 |
| I.Macs | Basp1 |
| I.Macs | Ninj1 |
| I.Macs | Cmklr1 |
| I.Macs | Cx3cr1 |
| I.Macs | Pla2g7 |
| I.Macs | Il4i1 |
| I.Macs | Adap2 |
| I.Macs | Rgs10 |
| I.Macs | Ccl22 |
| I.Macs | Cd74 |
| I.Macs | Ms4a6b |
| I.Macs | Mef2c |
| I.Macs | Rassf2 |
| I.Macs | H2-Eb1 |
| I.Macs | H2-Aa |
| I.Macs | Blnk |
| I.Macs | Csf1r |
| I.Macs | Adap2os |
| I.Macs | Cd83 |
| I.Macs | Il7r |
| I.Macs | Ptafr |
| I.Macs | Gatm |
| I.Macs | Ccl8 |
| I.Macs | Cst3 |
| I.Macs | Slc11a1 |
| I.Macs | Spred1 |
| I.Macs | Fyb |
| I.Macs | Cd81 |
| I.Macs | Gas7 |
| I.Macs | Stard8 |
| I.Macs | Man1a |
| I.Macs | Ifitm2 |
| I.Macs | Traf1 |
| I.Macs | Ms4a6c |
| I.Macs | Clec4a1 |
| I.Macs | Epsti1 |
| I.Macs | Ccl9 |
| I.Macs | Rogdi |
| I.Macs | Serpinb6b |
| I.Macs | Ifi27l2a |
| I.Macs | Gpr65 |
| I.Macs | B4galt6 |
| I.Macs | Anxa3 |
| I.Macs | Hspa1a |
| I.Macs | Ly86 |
| I.Macs | Arl4c |
| I.Macs | Gpr34 |
| I.Macs | Tagap |
| I.Macs | Arhgap22 |
| I.Macs | Relb |
| I.Macs | B2m |
| I.Macs | Lpcat2 |
| I.Macs | H2-Ab1 |

**Supplementary Table 1: Cluster defining genes in scRNAseq**

| | | | | | |
|---|---|---|---|---|---|
| I.Macs | Pid1 | I.Macs | Myo5a | I.Macs | Bmp2k |
| I.Macs | Rgs1 | I.Macs | Adam8 | I.Macs | H2-D1 |
| I.Macs | Lifr | I.Macs | Lst1 | I.Macs | Tcf4 |
| I.Macs | Pea15a | I.Macs | Malat1 | I.Macs | Bin1 |
| I.Macs | Gpr183 | I.Macs | Fcgr3 | I.Macs | Rtn4 |
| I.Macs | Gadd45b | I.Macs | Hexb | I.Macs | Fam60a |
| I.Macs | Tubb2a | I.Macs | Socs3 | I.Macs | Phf11b |
| I.Macs | Fcgr2b | I.Macs | Ccnd1 | I.Macs | Eno3 |
| I.Macs | Smagp | I.Macs | Slamf7 | I.Macs | Map3k14 |
| I.Macs | Rbpj | I.Macs | Serpinb9 | I.Macs | Cysltr1 |
| I.Macs | Cnn2 | I.Macs | Il10rb | I.Macs | Neurl3 |
| I.Macs | Zfhx3 | I.Macs | Tuba1a | I.Macs | Ahr |
| I.Macs | Crip1 | I.Macs | Dusp2 | I.Macs | Birc2 |
| I.Macs | Pou2f2 | I.Macs | Dok1 | I.Macs | Rab3il1 |
| I.Macs | Apobec3 | I.Macs | Ppp1r15a | I.Macs | Tmem123 |
| I.Macs | Tbc1d4 | I.Macs | Swap70 | I.Macs | H2-Q7 |
| I.Macs | Cd86 | I.Macs | Nfkbia | I.Macs | Lpar6 |
| I.Macs | Frmd4b | I.Macs | Susd3 | I.Macs | Cyp27a1 |
| I.Macs | Etv3 | I.Macs | Serinc3 | I.Macs | Birc3 |
| I.Macs | Plekho1 | I.Macs | Cadm1 | I.Macs | Htra2 |
| I.Macs | Tsc22d3 | I.Macs | Rabgap1l | I.Macs | H2-DMb1 |
| I.Macs | Itm2b | I.Macs | Lyl1 | I.Macs | Acer3 |
| I.Macs | Plxnb2 | I.Macs | Lat2 | I.Macs | Trem2 |
| I.Macs | Prkcb | I.Macs | Ptprj | I.Macs | Ntpcr |
| I.Macs | F13a1 | I.Macs | Ptpro | I.Macs | Runx3 |
| I.Macs | Hfe | I.Macs | Ptpn18 | I.Macs | Rnase4 |
| I.Macs | Fam49a | I.Macs | P2rx4 | I.Macs | St8sia4 |
| I.Macs | Lacc1 | I.Macs | Wfdc17 | I.Macs | Pmaip1 |
| I.Macs | Sepp1 | I.Macs | H2-Q6 | I.Macs | Tifa |
| I.Macs | Fgd2 | I.Macs | Ehd4 | I.Macs | Arrb2 |
| I.Macs | Ifitm3 | I.Macs | Il10ra | I.Macs | Blvrb |
| I.Macs | Nfkbiz | I.Macs | Scpep1 | I.Macs | Cd14 |
| I.Macs | Zcchc11 | I.Macs | Ly96 | I.Macs | Ogfrl1 |
| I.Macs | Ccr5 | I.Macs | Got1 | I.Macs | Nav1 |
| I.Macs | Abca9 | I.Macs | P2ry6 | I.Macs | Itgam |
| I.Macs | Ctsh | I.Macs | Rps29 | I.Macs | Lpcat1 |
| I.Macs | Ccl2 | I.Macs | Gpr132 | I.Macs | Oasl2 |
| I.Macs | Batf3 | I.Macs | Ctsb | I.Macs | Srgn |
| I.Macs | Atp2b1 | I.Macs | Bmyc | I.Macs | Il15 |
| I.Macs | Limd2 | I.Macs | H2-K1 | I.Macs | Hist3h2a |
| I.Macs | Clec10a | I.Macs | Unc93b1 | I.Macs | Irf1 |
| I.Macs | Rcsd1 | I.Macs | Tbc1d8 | I.Macs | Adgre5 |
| I.Macs | Arl5c | I.Macs | Tnip3 | I.Macs | Fchsd2 |

**Supplementary Table 1: Cluster defining genes in scRNAseq**

| | | | | | |
|---|---|---|---|---|---|
| I.Macs | Rtp4 | I.Macs | S100a4 | I.Macs | Ppp1r12a |
| I.Macs | Bst2 | I.Macs | Arap1 | I.Macs | Aldh2 |
| I.Macs | Wwp1 | I.Macs | Sp100 | I.Macs | Triap1 |
| I.Macs | Ccdc50 | I.Macs | Rap1b | I.Macs | N4bp2l1 |
| I.Macs | Adrbk2 | I.Macs | Rgs18 | I.Macs | Clec12a |
| I.Macs | BC028528 | I.Macs | Pstpip1 | I.Macs | Cd300a |
| I.Macs | Wnk1 | I.Macs | Ccl4 | I.Macs | Nfkb2 |
| I.Macs | Rnaset2a | I.Macs | Tmem243 | I.Macs | Hmha1 |
| I.Macs | Cmtm7 | I.Macs | Tapbpl | I.Macs | Psmb9 |
| I.Macs | Rpl3 | I.Macs | Ccdc109b | I.Macs | Avpi1 |
| I.Macs | Parp14 | I.Macs | Slc29a3 | I.Macs | Ncf4 |
| I.Macs | Csrp1 | I.Macs | Npc2 | I.Macs | Sbf2 |
| I.Macs | Adgre1 | I.Macs | Jun | I.Macs | Herc6 |
| I.Macs | Cbfa2t3 | I.Macs | Pltp | I.Macs | Itga4 |
| I.Macs | Apobec1 | I.Macs | Efhd2 | I.Macs | Evi2a |
| I.Macs | Lrrc25 | I.Macs | Stap1 | I.Macs | Tctex1d2 |
| I.Macs | Msr1 | I.Macs | Mthfd2 | I.Macs | Ly6e |
| I.Macs | Dennd5a | I.Macs | Rpl18a | I.Macs | Snx6 |
| I.Macs | Antxr2 | I.Macs | Map4k4 | I.Macs | Ass1 |
| I.Macs | Ccr2 | I.Macs | Tspan13 | I.Macs | Tspo |
| I.Macs | Rnf19b | I.Macs | Mfsd11 | I.Macs | Lrrk1 |
| I.Macs | Adgre4 | I.Macs | Tpm4 | I.Macs | Mndal |
| I.Macs | Ifi204 | I.Macs | Prkcd | I.Macs | Lilr4b |
| I.Macs | H2-Q4 | I.Macs | Rnaset2b | I.Macs | Psme2 |
| I.Macs | Tank | I.Macs | Fam46a | I.Macs | Psme2b |
| I.Macs | Nfat5 | I.Macs | Rps27 | I.Macs | Vsir |
| I.Macs | Ubb | I.Macs | Fgl2 | I.Macs | Ypel3 |
| I.Macs | Ahcyl2 | I.Macs | Rel | I.Macs | Eid1 |
| I.Macs | Uvrag | I.Macs | Arhgap15 | I.Macs | Foxp1 |
| I.Macs | Gbp2 | I.Macs | Il1b | I.Macs | Pyhin1 |
| I.Macs | Ifnar1 | I.Macs | Ifi203 | I.Macs | Crlf3 |
| I.Macs | Ccl17 | I.Macs | Stxbp3 | I.Macs | Dok2 |
| I.Macs | Jak2 | I.Macs | Zeb2os | I.Macs | Ms4a6d |
| I.Macs | Irf5 | I.Macs | Gna12 | I.Macs | Fam32a |
| I.Macs | Dynll2 | I.Macs | Sec14l1 | I.Macs | Jarid2 |
| I.Macs | Ncoa3 | I.Macs | Pfkfb3 | I.Macs | Aph1c |
| I.Macs | Kctd12 | I.Macs | Sh2b3 | I.Macs | Mxd1 |
| I.Macs | Ctsc | I.Macs | Junb | I.Macs | AI607873 |
| I.Macs | Igsf6 | I.Macs | Itgb5 | I.Macs | Ap1b1 |
| I.Macs | Cd300ld | I.Macs | Eps15 | I.Macs | Ankle2 |
| I.Macs | Ddit4 | I.Macs | Ptger4 | I.Macs | Lmo2 |
| I.Macs | Abi3 | I.Macs | Pik3r1 | I.Macs | Polr3c |
| I.Macs | Abca1 | I.Macs | Ncoa7 | I.Macs | Spop |

**Supplementary Table 1: Cluster defining genes in scRNAseq**

| | |
|---|---|
| I.Macs | PISD |
| I.Macs | Ier3 |
| I.Macs | Hacd4 |
| I.Macs | Tes |
| I.Macs | Gng12 |
| I.Macs | Dusp6 |
| I.Macs | Irf7 |
| I.Macs | Ifngr1 |
| I.Macs | Tor3a |
| I.Macs | Psmb8 |
| I.Macs | Syngr2 |
| I.Macs | Ktn1 |
| I.Macs | Tgif1 |
| I.Macs | Zfp36 |
| I.Macs | Tmem59 |
| I.Macs | Ubc |
| I.Macs | Nrros |
| I.Macs | Psmg4 |
| I.Macs | Rnf213 |
| I.Macs | Necap2 |
| I.Macs | Cyth4 |
| I.Macs | Iscu |
| I.Macs | Fcgr1 |
| I.Macs | Pepd |
| I.Macs | Isg15 |
| I.Macs | Prkacb |
| I.Macs | Fam177a |
| I.Macs | Fli1 |
| I.Macs | Sh3kbp1 |
| I.Macs | Tubb6 |
| I.Macs | Tnfaip8 |
| I.Macs | Sepw1 |
| I.Macs | Itm2c |
| I.Macs | Ccl5 |
| I.Macs | Nisch |
| I.Macs | Cited2 |
| I.Macs | Dapp1 |
| I.Macs | Vwa5a |
| I.Macs | Nfkbib |
| I.Macs | Tcn2 |
| I.Macs | Tab2 |
| I.Macs | Bcl10 |
| I.Macs | Azi2 |
| I.Macs | Snx2 |
| I.Macs | Slc3a2 |
| I.Macs | Klf6 |
| I.Macs | Myo1g |
| I.Macs | Fnbp1 |
| I.Macs | Gm26522 |
| I.Macs | Nars |
| I.Macs | Dnajb1 |
| I.Macs | Selplg |
| I.Macs | Snx5 |
| I.Macs | Tmem261 |
| I.Macs | Cltc |
| I.Macs | Idi1 |
| I.Macs | Cmtm6 |
| I.Macs | Herpud1 |
| I.Macs | Atf3 |
| I.Macs | Daglb |
| I.Macs | Ctnna1 |
| I.Macs | Marf1 |
| I.Macs | Rap2b |
| I.Macs | Samsn1 |
| I.Macs | Nudt9 |
| I.Macs | Kdm6b |
| I.Macs | Prdx4 |
| I.Macs | Rnf115 |
| I.Macs | Gpbp1 |
| I.Macs | Mef2a |
| I.Macs | Snx3 |
| I.Macs | Psme1 |
| I.Macs | Fkbp3 |
| I.Macs | Zscan26 |
| I.Macs | Pld4 |
| I.Macs | Ccl3 |
| I.Macs | Bcl2a1b |
| I.Macs | Sptssa |
| I.Macs | Hist1h1c |
| I.Macs | Rgs2 |
| I.Macs | Atox1 |
| I.Macs | Tnfaip3 |
| I.Macs | Sub1 |
| I.Macs | Zfand6 |
| I.Macs | Pnrc1 |
| I.Macs | Pim1 |
| I.Macs | Dnajb6 |
| I.Macs | Id2 |
| I.Macs | Socs2 |
| I.Macs | Btg1 |
| A.Macs | Chil3 |
| A.Macs | Ear2 |
| A.Macs | Cd9 |
| A.Macs | Ctsd |
| A.Macs | Abcg1 |
| A.Macs | Cd44 |
| A.Macs | Plet1 |
| A.Macs | Lpl |
| A.Macs | Ear1 |
| A.Macs | Fth1 |
| A.Macs | Ltc4s |
| A.Macs | Sgk1 |
| A.Macs | Tcf7l2 |
| A.Macs | Ccl6 |
| A.Macs | Atp6v0d2 |
| A.Macs | Cd164 |
| A.Macs | Mrc1 |
| A.Macs | Fabp1 |
| A.Macs | Krt19 |
| A.Macs | Pld3 |
| A.Macs | Wfdc21 |
| A.Macs | Axl |
| A.Macs | Hebp1 |
| A.Macs | Ftl1 |
| A.Macs | Plin2 |
| A.Macs | mt-Co1 |
| A.Macs | Nceh1 |
| A.Macs | Krt79 |
| A.Macs | Reep5 |
| A.Macs | mt-Co3 |
| A.Macs | App |
| A.Macs | Acaa1b |
| A.Macs | Klhdc4 |
| A.Macs | Prdx1 |
| A.Macs | Il18 |
| A.Macs | Sirpa |

**Supplementary Table 1: Cluster defining genes in scRNAseq**

| | |
|---|---|
| A.Macs | C5ar1 |
| A.Macs | Clec4n |
| A.Macs | Iqgap1 |
| A.Macs | Cd2 |
| A.Macs | Laptm5 |
| A.Macs | Adipor2 |
| A.Macs | Olr1 |
| A.Macs | Slpi |
| A.Macs | AU020206 |
| A.Macs | F7 |
| A.Macs | Aprt |
| A.Macs | Ptpn12 |
| A.Macs | Cidec |
| A.Macs | Cdc42ep3 |
| A.Macs | Aplp2 |
| A.Macs | mt-Cytb |
| A.Macs | mt-Nd4 |
| A.Macs | AI504432 |
| A.Macs | Slc6a6 |
| A.Macs | Mcemp1 |
| A.Macs | Hvcn1 |
| A.Macs | Slc7a2 |
| A.Macs | Fpr1 |
| A.Macs | Tnfaip2 |
| A.Macs | Lmna |
| A.Macs | Cstb |
| A.Macs | Dmxl2 |
| A.Macs | Anxa2 |
| A.Macs | mt-Nd4l |
| A.Macs | Trf |
| A.Macs | Txnip |
| A.Macs | S100a1 |
| A.Macs | Lpin1 |
| A.Macs | mt-Co2 |
| A.Macs | Fpr2 |
| A.Macs | Mt1 |
| A.Macs | Net1 |
| A.Macs | Runx1 |
| A.Macs | Dab2 |
| A.Macs | Txn1 |
| A.Macs | Dbi |
| A.Macs | Capg |
| A.Macs | Kazald1 |
| A.Macs | Pnpla8 |
| A.Macs | Ctss |
| A.Macs | Siglecf |
| A.Macs | Glul |
| A.Macs | Car4 |
| A.Macs | Cd302 |
| A.Macs | Dst |
| A.Macs | Dstn |
| A.Macs | Lmo4 |
| A.Macs | Lrp1 |
| A.Macs | Baz1a |
| A.Macs | Perp |
| A.Macs | Ctnnb1 |
| A.Macs | Pla2g15 |
| A.Macs | mt-Nd2 |
| A.Macs | mt-Atp6 |
| A.Macs | mt-Nd3 |
| A.Macs | Gpcpd1 |
| A.Macs | Cebpb |
| A.Macs | Hist1h2bc |
| A.Macs | Abhd12 |
| A.Macs | S100a11 |
| A.Macs | mt-Nd1 |
| A.Macs | Spp1 |
| A.Macs | Lgals3 |
| A.Macs | Atxn1 |
| A.Macs | Cox5a |
| A.Macs | Rexo2 |
| A.Macs | Il1rn |
| A.Macs | Msrb1 |
| A.Macs | Serpinb1a |
| A.Macs | Blvra |
| A.Macs | Ramp1 |
| A.Macs | Sort1 |
| A.Macs | Slc15a3 |
| A.Macs | Kcnn3 |
| A.Macs | Gns |
| A.Macs | Sulf2 |
| A.Macs | Hsp90b1 |
| A.Macs | Nabp1 |
| A.Macs | Prkar2b |
| A.Macs | Myl6 |
| A.Macs | Akr1a1 |
| A.Macs | Cd300lf |
| A.Macs | Mgst1 |
| A.Macs | Ccnd2 |
| A.Macs | Itpripl2 |
| A.Macs | Gal |
| A.Macs | Mpeg1 |
| A.Macs | Itgax |
| A.Macs | Atp6v1g1 |
| A.Macs | Etfb |
| A.Macs | Canx |
| A.Macs | Fabp5 |
| A.Macs | Kcnq1ot1 |
| A.Macs | Sep-09 |
| A.Macs | Phgdh |
| A.Macs | Pald1 |
| A.Macs | Atp13a3 |
| A.Macs | Camk1 |
| A.Macs | Pros1 |
| A.Macs | Dhrs3 |
| A.Macs | Atp5g3 |
| A.Macs | Lrp12 |
| A.Macs | Tlr2 |
| A.Macs | Lima1 |
| A.Macs | Egr2 |
| A.Macs | Gpx4 |
| A.Macs | Gstm1 |
| A.Macs | Tob1 |
| A.Macs | Aldoa |
| A.Macs | Serpine1 |
| A.Macs | Calr |
| A.Macs | Tmbim1 |
| A.Macs | Phlda1 |
| A.Macs | Plaur |
| A.Macs | Camk2d |
| A.Macs | Ptp4a1 |
| A.Macs | Bst1 |
| A.Macs | Ucp2 |
| A.Macs | Mapk3 |
| A.Macs | Snx10 |
| A.Macs | Mertk |
| A.Macs | Actn1 |
| A.Macs | Pygl |
| A.Macs | Alas1 |

**Supplementary Table 1: Cluster defining genes in scRNAseq**

| | | | | | |
|---|---|---|---|---|---|
| A.Macs | Cd200r4 | A.Macs | Comt | A.Macs | Tpp1 |
| A.Macs | Anxa5 | A.Macs | Chp1 | A.Macs | Lrrfip1 |
| A.Macs | Slc9a4 | A.Macs | Myo7a | A.Macs | Atp6v0b |
| A.Macs | Gabarap | A.Macs | Gcnt2 | A.Macs | Lipa |
| A.Macs | Mgll | A.Macs | Ppp1r14b | A.Macs | Spns1 |
| A.Macs | Cpt1a | A.Macs | Xbp1 | A.Macs | Scp2 |
| A.Macs | Bhlhe41 | A.Macs | Cyba | A.Macs | Cib2 |
| A.Macs | Tfec | A.Macs | Cd84 | A.Macs | Lgals1 |
| A.Macs | Unc119 | A.Macs | Npc1 | A.Macs | Mapk6 |
| A.Macs | Tmem14c | A.Macs | Cyb5r3 | A.Macs | Myof |
| A.Macs | Uqcr11 | A.Macs | Ndufa4 | A.Macs | Adcy3 |
| A.Macs | Frmd4a | A.Macs | Gsap | A.Macs | Creg1 |
| A.Macs | Furin | A.Macs | Acp5 | A.Macs | B3gnt7 |
| A.Macs | Sh2d1b1 | A.Macs | Ctsl | A.Macs | Pnpla7 |
| A.Macs | Sdcbp | A.Macs | Hmgn2 | A.Macs | Noct |
| A.Macs | Clic4 | A.Macs | Itgal | A.Macs | Ywhab |
| A.Macs | Bnip3l | A.Macs | Abracl | A.Macs | Sh3bgrl2 |
| A.Macs | AW112010 | A.Macs | Rhoc | A.Macs | Tc2n |
| A.Macs | Mcoln3 | A.Macs | Flt1 | A.Macs | Adarb1 |
| A.Macs | Tgfbr2 | A.Macs | Pag1 | A.Macs | Ezr |
| A.Macs | Gm10116 | A.Macs | Chic2 | A.Macs | Ski |
| A.Macs | Cytip | A.Macs | Atp6v0e | A.Macs | Fcor |
| A.Macs | Ddhd1 | A.Macs | Fcgrt | A.Macs | Rtn3 |
| A.Macs | Scarb2 | A.Macs | Abcd2 | A.Macs | Anxa4 |
| A.Macs | Cpne5 | A.Macs | Cox4i1 | A.Macs | Rab44 |
| A.Macs | Afap1 | A.Macs | Tmem154 | A.Macs | F10 |
| A.Macs | Marco | A.Macs | Acox1 | A.Macs | Alox5 |
| A.Macs | Ms4a8a | A.Macs | Plgrkt | A.Macs | mt-Atp8 |
| A.Macs | Zfp703 | A.Macs | Dhrs7b | A.Macs | Ech1 |
| A.Macs | Card11 | A.Macs | Lilra5 | A.Macs | Grn |
| A.Macs | Irf2bp2 | A.Macs | Rhob | A.Macs | Akr1b3 |
| A.Macs | Angptl4 | A.Macs | Cox6b1 | A.Macs | Clmn |
| A.Macs | Nrip1 | A.Macs | Klhl9 | A.Macs | Serpinb6a |
| A.Macs | Atp6v1b2 | A.Macs | Rplp1 | A.Macs | Snx1 |
| A.Macs | Evl | A.Macs | Abhd5 | A.Macs | Hsd17b4 |
| A.Macs | Colgalt1 | A.Macs | Cenpb | A.Macs | Abcc5 |
| A.Macs | Lamtor2 | A.Macs | Tns1 | A.Macs | Cndp2 |
| A.Macs | Cybb | A.Macs | Ctsk | A.Macs | Tmbim6 |
| A.Macs | Idh1 | A.Macs | Gapdh | A.Macs | P4hb |
| A.Macs | Pon2 | A.Macs | Ptgfrn | A.Macs | Romo1 |
| A.Macs | Ptbp3 | A.Macs | Capn1 | A.Macs | Tagln2 |
| A.Macs | Pilra | A.Macs | Myh9 | A.Macs | Irs2 |
| A.Macs | Cd47 | A.Macs | Iqsec1 | A.Macs | Cttnbp2nl |

**Supplementary Table 1: Cluster defining genes in scRNAseq**

| | | | | | |
|---|---|---|---|---|---|
| A.Macs | Cpne8 | A.Macs | Tbca | A.Macs | Ptms |
| A.Macs | Prdx5 | A.Macs | Slc12a7 | A.Macs | Mycbp2 |
| A.Macs | Ttyh2 | A.Macs | Uqcrb | A.Macs | Tor1aip2 |
| A.Macs | Pparg | A.Macs | Qdpr | A.Macs | Emilin1 |
| A.Macs | Gm42418 | A.Macs | Plekhg1 | A.Macs | B4galnt1 |
| A.Macs | Hcar2 | A.Macs | Cox6c | A.Macs | Lcor |
| A.Macs | Gpnmb | A.Macs | Nus1 | A.Macs | AF251705 |
| A.Macs | Plek | A.Macs | Slc16a10 | A.Macs | Gas2l1 |
| A.Macs | Atp6v1e1 | A.Macs | Reep3 | A.Macs | Pdia4 |
| A.Macs | Trim25 | A.Macs | Amz1 | A.Macs | Ifrd1 |
| A.Macs | Tgm2 | A.Macs | Vamp8 | A.Macs | Soat1 |
| A.Macs | Dapk1 | A.Macs | Mcl1 | A.Macs | Nedd9 |
| A.Macs | Psen2 | A.Macs | Gda | A.Macs | Sypl |
| A.Macs | Hspa5 | A.Macs | Fosl2 | A.Macs | Dnase2a |
| A.Macs | C530008M17Rik | A.Macs | Trappc2l | A.Macs | Lyrm4 |
| A.Macs | Renbp | A.Macs | Fam129a | A.Macs | Larp4b |
| A.Macs | Lamp1 | A.Macs | Mgat4b | A.Macs | Lcp1 |
| A.Macs | Ly75 | A.Macs | Dnajb9 | A.Macs | Sla |
| A.Macs | Cdk6 | A.Macs | Samd8 | A.Macs | Mfsd12 |
| A.Macs | Nme1 | A.Macs | Acot1 | A.Macs | Tgfbr1 |
| A.Macs | Hcfc1r1 | A.Macs | Mmd | A.Macs | Gadd45g |
| A.Macs | Ctsa | A.Macs | Atxn10 | A.Macs | Mapk1 |
| A.Macs | Plk3 | A.Macs | Cd200r1 | A.Macs | 2010107E04Rik |
| A.Macs | F11r | A.Macs | Fam89a | A.Macs | Abhd17c |
| A.Macs | Trim29 | A.Macs | Csf2rb | A.Macs | Ppt2 |
| A.Macs | Creb5 | A.Macs | Gnb1 | A.Macs | Cd33 |
| A.Macs | Gsto1 | A.Macs | Dock10 | A.Macs | Hiatl1 |
| A.Macs | Plscr1 | A.Macs | Hprt | A.Macs | Ctsz |
| A.Macs | Mgst3 | A.Macs | Shtn1 | A.Macs | Lpp |
| A.Macs | Hexa | A.Macs | C2cd2l | A.Macs | Ggta1 |
| A.Macs | Mpc2 | A.Macs | Neu1 | A.Macs | mt-Nd5 |
| A.Macs | A930007I19Rik | A.Macs | Insr | A.Macs | Tmcc3 |
| A.Macs | Inadl | A.Macs | Pqlc3 | A.Macs | Ano6 |
| A.Macs | Chchd2 | A.Macs | Clta | A.Macs | Cyth3 |
| A.Macs | Vat1 | A.Macs | Flna | A.Macs | Vim |
| A.Macs | Bcar3 | A.Macs | Acaa1a | A.Macs | Pcyox1 |
| A.Macs | Gpr155 | A.Macs | Gng5 | A.Macs | Rnh1 |
| A.Macs | Bcl2a1a | A.Macs | Fndc3b | A.Macs | Taldo1 |
| A.Macs | Cebpa | A.Macs | Map1lc3b | A.Macs | Rxra |
| A.Macs | Picalm | A.Macs | Trpv2 | A.Macs | S100a10 |
| A.Macs | Diaph1 | A.Macs | S100a13 | A.Macs | Nck1 |
| A.Macs | Abr | A.Macs | Pgd | A.Macs | Adgre1 |
| A.Macs | Slc36a4 | A.Macs | Eif4g2 | A.Macs | Bola2 |

**Supplementary Table 1: Cluster defining genes in scRNAseq**

| | | | | | |
|---|---|---|---|---|---|
| A.Macs | Plekhb2 | A.Macs | Ppp2cb | A.Macs | Sqstm1 |
| A.Macs | Nucb2 | A.Macs | Gnaq | A.Macs | Cox6b2 |
| A.Macs | Hnrnpa0 | A.Macs | Atp6v1a | A.Macs | Mif |
| A.Macs | Pdlim1 | A.Macs | Wdfy3 | A.Macs | Sav1 |
| A.Macs | Actr2 | A.Macs | Tuba4a | A.Macs | Arhgap31 |
| A.Macs | Asph | A.Macs | Ap3s1 | A.Macs | Apbb1ip |
| A.Macs | Cxcl2 | A.Macs | Ndufa2 | A.Macs | Pygo2 |
| A.Macs | Sep-11 | A.Macs | Siglece | A.Macs | Atp1b3 |
| A.Macs | Ccni | A.Macs | Snx27 | A.Macs | Frrs1 |
| A.Macs | Bzw2 | A.Macs | Mrps15 | A.Macs | Vdac2 |
| A.Macs | Cd274 | A.Macs | Man2a2 | A.Macs | Lgals8 |
| A.Macs | Clec2d | A.Macs | Fosb | A.Macs | Sh3bgrl |
| A.Macs | Tmed5 | A.Macs | Luzp1 | A.Macs | Map1lc3a |
| A.Macs | Mpp1 | A.Macs | Ndufa5 | A.Macs | Runx2 |
| A.Macs | Ndufv3 | A.Macs | Mum1 | A.Macs | Timm13 |
| A.Macs | Tmem238 | A.Macs | Arl8a | A.Macs | Acads |
| A.Macs | Lsr | A.Macs | Ctnnd1 | A.Macs | Arrdc4 |
| A.Macs | Znrf1 | A.Macs | BC005537 | A.Macs | Myl12a |
| A.Macs | Slc29a1 | A.Macs | Svil | A.Macs | Tmem189 |
| A.Macs | Gmds | A.Macs | Atp13a2 | A.Macs | Nufip2 |
| A.Macs | Ugcg | A.Macs | Atp6v0d1 | A.Macs | Fkbp2 |
| A.Macs | Dynll1 | A.Macs | Fabp4 | A.Macs | M6pr |
| A.Macs | Pfkfb4 | A.Macs | Tbxas1 | A.Macs | 1600014C10Rik |
| A.Macs | Ubash3b | A.Macs | Manf | A.Macs | Krcc1 |
| A.Macs | Pon3 | A.Macs | Mroh1 | A.Macs | Dennd4c |
| A.Macs | Rassf3 | A.Macs | Sh3tc1 | A.Macs | Srp14 |
| A.Macs | Sgpl1 | A.Macs | Lrrc58 | A.Macs | Snrk |
| A.Macs | Cyp4v3 | A.Macs | Tmem256 | A.Macs | Hspa9 |
| A.Macs | Uhmk1 | A.Macs | Rhoq | A.Macs | Klf7 |
| A.Macs | Atp6ap2 | A.Macs | Vkorc1l1 | A.Macs | Mpc1 |
| A.Macs | Cltc | A.Macs | Wdr1 | A.Macs | Tbc1d9 |
| A.Macs | Grina | A.Macs | Ddx3x | A.Macs | Dync1i2 |
| A.Macs | Ifi27 | A.Macs | Mrpl51 | A.Macs | Rassf5 |
| A.Macs | Dpep2 | A.Macs | Tnfrsf21 | A.Macs | Ivns1abp |
| A.Macs | Rpn1 | A.Macs | Hadhb | A.Macs | Gls |
| A.Macs | Atp6v1f | A.Macs | Tgfb1 | A.Macs | Fam118b |
| A.Macs | Papd5 | A.Macs | Top1 | A.Macs | Ptk2b |
| A.Macs | Mrps6 | A.Macs | Lamp2 | A.Macs | Nagk |
| A.Macs | Mien1 | A.Macs | Polr2a | A.Macs | Rabac1 |
| A.Macs | Eif3a | A.Macs | Rab1a | A.Macs | Rnf149 |
| A.Macs | Pdia6 | A.Macs | Atp6ap1 | A.Macs | Sel1l |
| A.Macs | Cox6a1 | A.Macs | Hpcal1 | A.Macs | Qk |
| A.Macs | Mtss1 | A.Macs | Hadha | A.Macs | Saraf |

**Supplementary Table 1: Cluster defining genes in scRNAseq**

| | | | | | |
|---|---|---|---|---|---|
| A.Macs | Atp6v1c1 | A.Macs | Ubqln1 | A.Macs | Myo9b |
| A.Macs | Fbxl5 | A.Macs | Emc8 | A.Macs | Aebp2 |
| A.Macs | Mfsd1 | A.Macs | Gmcl1 | A.Macs | Rock2 |
| A.Macs | P2ry14 | A.Macs | Rps6ka3 | A.Macs | Cib1 |
| A.Macs | Rab32 | A.Macs | Ndufs5 | A.Macs | Klf10 |
| A.Macs | Cd36 | A.Macs | Cav2 | A.Macs | Mark2 |
| A.Macs | Rras2 | A.Macs | Tbc1d20 | A.Macs | Ffar4 |
| A.Macs | Ndufa1 | A.Macs | Tmem160 | A.Macs | Riok3 |
| A.Macs | Hip1 | A.Macs | Map3k5 | A.Macs | Usp50 |
| A.Macs | Dnajc5 | A.Macs | Akap13 | A.Macs | Adam17 |
| A.Macs | Rin2 | A.Macs | Esyt2 | A.Macs | Chd4 |
| A.Macs | Diaph2 | A.Macs | Mfsd5 | A.Macs | Mical1 |
| A.Macs | Cln3 | A.Macs | Stxbp2 | A.Macs | Plod3 |
| A.Macs | Gpr137b | A.Macs | Adam9 | A.Macs | Aldh3b1 |
| A.Macs | Arl8b | A.Macs | Arhgef2 | A.Macs | Dip2b |
| A.Macs | Ccr1 | A.Macs | Sidt2 | A.Macs | Tspan14 |
| A.Macs | Copa | A.Macs | C3 | A.Macs | Tex264 |
| A.Macs | Fam168b | A.Macs | Hcst | A.Macs | Sod2 |
| A.Macs | Arf3 | A.Macs | Casp8 | A.Macs | Gars |
| A.Macs | Stom | A.Macs | Anxa11 | A.Macs | Itgb5 |
| A.Macs | Hnrnph1 | A.Macs | Vps29 | A.Macs | Gak |
| A.Macs | Lrrc8d | A.Macs | Cnpy2 | A.Macs | Gm10263 |
| A.Macs | N4bp1 | A.Macs | Usf2 | A.Macs | Rab8b |
| A.Macs | Atp5g1 | A.Macs | Znhit1 | A.Macs | Morf4l2 |
| A.Macs | Mt2 | A.Macs | Tnks2 | A.Macs | Capza1 |
| A.Macs | Emc7 | A.Macs | Il4ra | A.Macs | Purb |
| A.Macs | Bcl2l11 | A.Macs | Ddb1 | A.Macs | Rragc |
| A.Macs | Abca1 | A.Macs | Commd3 | A.Macs | Nipa2 |
| A.Macs | Il6st | A.Macs | Trip12 | A.Macs | Cat |
| A.Macs | B4galt1 | A.Macs | Adrb2 | A.Macs | Acadm |
| A.Macs | Acly | A.Macs | Slc25a36 | A.Macs | Stt3b |
| A.Macs | Senp2 | A.Macs | Pdcd6ip | A.Macs | Sptlc2 |
| A.Macs | Slc39a1 | A.Macs | Naa50 | A.Macs | Ppp1r18 |
| A.Macs | Fam96a | A.Macs | Tgoln1 | A.Macs | Taok1 |
| A.Macs | Tusc3 | A.Macs | Sec63 | A.Macs | Sowahc |
| A.Macs | Fry | A.Macs | B930036N10Rik | A.Macs | Tmod3 |
| A.Macs | Sec31a | A.Macs | Nsf | A.Macs | 2610001J05Rik |
| A.Macs | Ndufab1 | A.Macs | Gmfb | A.Macs | Wdr83os |
| A.Macs | Eif5b | A.Macs | Nktr | A.Macs | Srsf6 |
| A.Macs | Sh3glb1 | A.Macs | Nenf | A.Macs | Parl |
| A.Macs | Pnpla2 | A.Macs | Batf | A.Macs | Zdhhc21 |
| A.Macs | Setd8 | A.Macs | Ndufb2 | A.Macs | Tulp4 |
| A.Macs | Mylip | A.Macs | Nfix | A.Macs | Far1 |

**Supplementary Table 1: Cluster defining genes in scRNAseq**

| | |
|---|---|
| A.Macs | Pik3r5 |
| A.Macs | Cap1 |
| A.Macs | Etfa |
| A.Macs | Uap1l1 |
| A.Macs | Gm26917 |
| A.Macs | Epb41l2 |
| A.Macs | Ogdh |
| A.Macs | Esyt1 |
| A.Macs | Hmgcs1 |
| A.Macs | Acot13 |
| A.Macs | Fam168a |
| A.Macs | Fkbp5 |
| A.Macs | Eci2 |
| A.Macs | Npepps |
| A.Macs | Elovl5 |
| A.Macs | Kif1b |
| A.Macs | 1110008P14Rik |
| A.Macs | Ndufb6 |
| A.Macs | Trib1 |
| A.Macs | Ggnbp2 |
| A.Macs | Egr1 |
| A.Macs | Atp6v1d |
| A.Macs | Per1 |
| A.Macs | Csnk1e |
| A.Macs | Cisd1 |
| A.Macs | Tcirg1 |
| A.Macs | Acox3 |
| A.Macs | Syk |
| A.Macs | Arl11 |
| A.Macs | Tmem168 |
| A.Macs | Dync1h1 |
| A.Macs | Rab11fip1 |
| A.Macs | Ddx6 |
| A.Macs | Cyp4f16 |
| A.Macs | Atp2a2 |
| A.Macs | Ddx46 |
| A.Macs | Ap1s1 |
| A.Macs | Sec61a1 |
| A.Macs | Mxd4 |
| A.Macs | Rab6a |
| A.Macs | Hnrnpll |
| A.Macs | Plin3 |

| | |
|---|---|
| Prolif. A.Macs | 2810417H13Rik |
| Prolif. A.Macs | Top2a |
| Prolif. A.Macs | Nusap1 |
| Prolif. A.Macs | Mki67 |
| Prolif. A.Macs | Rrm2 |
| Prolif. A.Macs | Cdca3 |
| Prolif. A.Macs | Ccna2 |
| Prolif. A.Macs | Kif15 |
| Prolif. A.Macs | Prc1 |
| Prolif. A.Macs | Cdk1 |
| Prolif. A.Macs | Ncapd2 |
| Prolif. A.Macs | Birc5 |
| Prolif. A.Macs | Tk1 |
| Prolif. A.Macs | Cenpf |
| Prolif. A.Macs | Uhrf1 |
| Prolif. A.Macs | Spc25 |
| Prolif. A.Macs | Esco2 |
| Prolif. A.Macs | Mis18bp1 |
| Prolif. A.Macs | Stmn1 |
| Prolif. A.Macs | Asf1b |
| Prolif. A.Macs | Lockd |
| Prolif. A.Macs | Pbk |
| Prolif. A.Macs | Ccnb2 |
| Prolif. A.Macs | Ccne2 |
| Prolif. A.Macs | Ube2c |
| Prolif. A.Macs | Neil3 |
| Prolif. A.Macs | Fbxo5 |

| | |
|---|---|
| Prolif. A.Macs | Hmmr |
| Prolif. A.Macs | Rad51 |
| Prolif. A.Macs | Aspm |
| Prolif. A.Macs | Ccnb1 |
| Prolif. A.Macs | Kif11 |
| Prolif. A.Macs | Sgol1 |
| Prolif. A.Macs | Cenpe |
| Prolif. A.Macs | Ncapg |
| Prolif. A.Macs | Racgap1 |
| Prolif. A.Macs | Rad51ap1 |
| Prolif. A.Macs | Rrm1 |
| Prolif. A.Macs | Tpx2 |
| Prolif. A.Macs | Lig1 |
| Prolif. A.Macs | Cdca8 |
| Prolif. A.Macs | Spc24 |
| Prolif. A.Macs | Ube2t |
| Prolif. A.Macs | Cep55 |
| Prolif. A.Macs | Smc2 |
| Prolif. A.Macs | Bub1 |
| Prolif. A.Macs | Hist1h2ap |
| Prolif. A.Macs | Casc5 |
| Prolif. A.Macs | Mad2l1 |
| Prolif. A.Macs | Hells |
| Prolif. A.Macs | Kif20a |
| Prolif. A.Macs | Ckap2l |
| Prolif. A.Macs | E2f7 |
| Prolif. A.Macs | Clspn |
| Prolif. A.Macs | Anln |
| Prolif. A.Macs | Ckap2 |
| Prolif. A.Macs | Nuf2 |

**Supplementary Table 1: Cluster defining genes in scRNAseq**

| | | | | | |
|---|---|---|---|---|---|
| Prolif. A.Macs | Kif23 | Prolif. A.Macs | Fam64a | Prolif. A.Macs | Pcna |
| Prolif. A.Macs | Atad2 | Prolif. A.Macs | Tipin | Prolif. A.Macs | Incenp |
| Prolif. A.Macs | Ncaph | Prolif. A.Macs | Ndc80 | Prolif. A.Macs | Fignl1 |
| Prolif. A.Macs | Shcbp1 | Prolif. A.Macs | Dtl | Prolif. A.Macs | Aurkb |
| Prolif. A.Macs | Tcf19 | Prolif. A.Macs | Brip1 | Prolif. A.Macs | Cdkn2c |
| Prolif. A.Macs | Ska1 | Prolif. A.Macs | Tuba1b | Prolif. A.Macs | Dnmt1 |
| Prolif. A.Macs | Kif22 | Prolif. A.Macs | Melk | Prolif. A.Macs | Lsm2 |
| Prolif. A.Macs | C330027C09Rik | Prolif. A.Macs | Mcm7 | Prolif. A.Macs | Zfp367 |
| Prolif. A.Macs | Ccdc34 | Prolif. A.Macs | Cks1b | Prolif. A.Macs | Haus4 |
| Prolif. A.Macs | Dut | Prolif. A.Macs | Tubb5 | Prolif. A.Macs | Nasp |
| Prolif. A.Macs | Kif2c | Prolif. A.Macs | Tyms | Prolif. A.Macs | Cenpm |
| Prolif. A.Macs | Cenpa | Prolif. A.Macs | Hmgb2 | Prolif. A.Macs | Diaph3 |
| Prolif. A.Macs | Cdc20 | Prolif. A.Macs | Brca2 | Prolif. A.Macs | Rfc4 |
| Prolif. A.Macs | Mxd3 | Prolif. A.Macs | Cenpk | Prolif. A.Macs | 2700094K13Rik |
| Prolif. A.Macs | Hist1h1b | Prolif. A.Macs | Ptma | Prolif. A.Macs | Ect2 |
| Prolif. A.Macs | Cit | Prolif. A.Macs | Tmpo | Prolif. A.Macs | Hmgb1 |
| Prolif. A.Macs | Ezh2 | Prolif. A.Macs | Plk4 | Prolif. A.Macs | Mcm2 |
| Prolif. A.Macs | Kifc1 | Prolif. A.Macs | Pkmyt1 | Prolif. A.Macs | Nrm |
| Prolif. A.Macs | Cenph | Prolif. A.Macs | E2f8 | Prolif. A.Macs | Cbx5 |
| Prolif. A.Macs | Tacc3 | Prolif. A.Macs | Cenpw | Prolif. A.Macs | Blm |
| Prolif. A.Macs | Cenpi | Prolif. A.Macs | Gmnn | Prolif. A.Macs | Rfc3 |
| Prolif. A.Macs | Ncapg2 | Prolif. A.Macs | Mcm5 | Prolif. A.Macs | Dek |
| Prolif. A.Macs | Depdc1a | Prolif. A.Macs | Prim1 | Prolif. A.Macs | Trip13 |
| Prolif. A.Macs | Sgol2a | Prolif. A.Macs | Ccnf | Prolif. A.Macs | Wdr76 |
| Prolif. A.Macs | Kif4 | Prolif. A.Macs | Cenpq | Prolif. A.Macs | Atad5 |
| Prolif. A.Macs | Kif20b | Prolif. A.Macs | Lmnb1 | Prolif. A.Macs | Topbp1 |
| Prolif. A.Macs | Hist1h2ae | Prolif. A.Macs | H2afv | Prolif. A.Macs | Bub1b |
| Prolif. A.Macs | Dlgap5 | Prolif. A.Macs | Smc4 | Prolif. A.Macs | Mcm6 |
| Prolif. A.Macs | Cdca2 | Prolif. A.Macs | Dbf4 | Prolif. A.Macs | Whsc1 |
| Prolif. A.Macs | Dnajc9 | Prolif. A.Macs | H2afz | Prolif. A.Macs | Ckap5 |

**Supplementary Table 1: Cluster defining genes in scRNAseq**

| | | | | | |
|---|---|---|---|---|---|
| Prolif. A.Macs | Cbx3 | Prolif. A.Macs | Slbp | Prolif. A.Macs | Xpo1 |
| Prolif. A.Macs | Ube2s | Prolif. A.Macs | Cntln | Prolif. A.Macs | Cmc2 |
| Prolif. A.Macs | Nucks1 | Prolif. A.Macs | Hnrnpab | Prolif. A.Macs | Smc1a |
| Prolif. A.Macs | Cdc25b | Prolif. A.Macs | Nap1l1 | Prolif. A.Macs | Anp32e |
| Prolif. A.Macs | Apitd1 | Prolif. A.Macs | Rpa3 | Prolif. A.Macs | Usp37 |
| Prolif. A.Macs | Ran | Prolif. A.Macs | 2700029M09Rik | Prolif. A.Macs | G2e3 |
| Prolif. A.Macs | Dhfr | Prolif. A.Macs | Syne2 | Prolif. A.Macs | Rcc1 |
| Prolif. A.Macs | Anp32b | Prolif. A.Macs | Prdx4 | Prolif. A.Macs | Lsm5 |
| Prolif. A.Macs | Rfc5 | Prolif. A.Macs | Cenpl | Prolif. A.Macs | Psat1 |
| Prolif. A.Macs | Hmgn2 | Prolif. A.Macs | Smc3 | Prolif. A.Macs | Psip1 |
| Prolif. A.Macs | H2afx | Prolif. A.Macs | Rbbp7 | Prolif. A.Macs | Hmgn5 |
| Prolif. A.Macs | Orc6 | Prolif. A.Macs | Pold3 | Prolif. A.Macs | Mb21d1 |
| Prolif. A.Macs | Cks2 | Prolif. A.Macs | Hjurp | Prolif. A.Macs | Wbp5 |
| Prolif. A.Macs | Tubb4b | Prolif. A.Macs | Fen1 | Prolif. A.Macs | Tmem109 |
| Prolif. A.Macs | Trim59 | Prolif. A.Macs | Gm10282 | Prolif. A.Macs | Cbfb |
| Prolif. A.Macs | Dck | Prolif. A.Macs | Rpa2 | Prolif. A.Macs | Spp1 |
| Prolif. A.Macs | Ncapd3 | Prolif. A.Macs | Ppia | Prolif. A.Macs | Pmf1 |
| Prolif. A.Macs | Kpna2 | Prolif. A.Macs | Dctpp1 | Prolif. A.Macs | Chek2 |
| Prolif. A.Macs | Mcm4 | Prolif. A.Macs | Naa40 | Prolif. A.Macs | Baz1b |
| Prolif. A.Macs | Hat1 | Prolif. A.Macs | Rangap1 | Prolif. A.Macs | Cenpc1 |
| Prolif. A.Macs | Chaf1a | Prolif. A.Macs | Gins2 | Prolif. A.Macs | Calm2 |
| Prolif. A.Macs | Alyref | Prolif. A.Macs | Hist1h1e | Prolif. A.Macs | Arl6ip1 |
| Prolif. A.Macs | Pola1 | Prolif. A.Macs | Myef2 | Prolif. A.Macs | Dtymk |
| Prolif. A.Macs | Syce2 | Prolif. A.Macs | Cdk2 | Prolif. A.Macs | Hsp90aa1 |
| Prolif. A.Macs | Mcm3 | Prolif. A.Macs | Cdt1 | Prolif. A.Macs | Eri1 |
| Prolif. A.Macs | Usp1 | Prolif. A.Macs | Arhgap11a | Prolif. A.Macs | A430005L14Rik |
| Prolif. A.Macs | Snrpd1 | Prolif. A.Macs | Mis18a | Prolif. A.Macs | Hdgf |
| Prolif. A.Macs | Ranbp1 | Prolif. A.Macs | Hmgn1 | Prolif. A.Macs | Lbr |
| Prolif. A.Macs | Rad21 | Prolif. A.Macs | Siva1 | Prolif. A.Macs | Rdm1 |
| Prolif. A.Macs | Rbl1 | Prolif. A.Macs | E2f2 | Prolif. A.Macs | Emp1 |

**Supplementary Table 1: Cluster defining genes in scRNAseq**

| | | | | | |
|---|---|---|---|---|---|
| Prolif. A.Macs | Smc6 | Prolif. A.Macs | Nudt4 | Prolif. A.Macs | Ybx1 |
| Prolif. A.Macs | Cep57 | Prolif. A.Macs | Nedd1 | Prolif. A.Macs | Cpd |
| Prolif. A.Macs | Tdp2 | Prolif. A.Macs | Fam111a | Prolif. A.Macs | Fam178a |
| Prolif. A.Macs | Nup205 | Prolif. A.Macs | Haus8 | Prolif. A.Macs | Anapc11 |
| Prolif. A.Macs | Tagln2 | Prolif. A.Macs | Pa2g4 | Prolif. A.Macs | Cdkn1a |
| Prolif. A.Macs | Tubg1 | Prolif. A.Macs | Slf1 | Prolif. A.Macs | Hn1 |
| Prolif. A.Macs | Aes | Prolif. A.Macs | Srsf7 | Prolif. A.Macs | Nudt21 |
| Prolif. A.Macs | Fmr1 | Prolif. A.Macs | Banf1 | Prolif. A.Macs | Paics |
| Prolif. A.Macs | Impdh2 | Prolif. A.Macs | Rnaseh2b | Prolif. A.Macs | Raly |
| Prolif. A.Macs | Cse1l | Prolif. A.Macs | Lsm3 | Prolif. A.Macs | Sae1 |
| Prolif. A.Macs | Rfwd3 | Prolif. A.Macs | Ilf3 | Prolif. A.Macs | Srsf3 |
| Prolif. A.Macs | Rif1 | Prolif. A.Macs | Nsmce1 | Prolif. A.Macs | Pde7a |
| Prolif. A.Macs | Nup85 | Prolif. A.Macs | Nxt2 | Prolif. A.Macs | Sypl |
| Prolif. A.Macs | Dpy30 | Prolif. A.Macs | Sumo2 | Prolif. A.Macs | Rbm15 |
| Prolif. A.Macs | Prim2 | Prolif. A.Macs | Ttf2 | Prolif. A.Macs | Gins4 |
| Prolif. A.Macs | 1810037I17Rik | Prolif. A.Macs | Pcbp2 | Prolif. A.Macs | Csrp1 |
| Prolif. A.Macs | Ssrp1 | Prolif. A.Macs | Brd8 | Prolif. A.Macs | Smarca5 |
| Prolif. A.Macs | Ctcf | Prolif. A.Macs | Brip1os | Prolif. A.Macs | Mis12 |
| Prolif. A.Macs | Tex30 | Prolif. A.Macs | Cep83 | Prolif. A.Macs | Prdx1 |
| Prolif. A.Macs | Phgdh | Prolif. A.Macs | Cdk4 | Prolif. A.Macs | Exosc8 |
| Prolif. A.Macs | Hint1 | Prolif. A.Macs | Hn1l | Prolif. A.Macs | Stra13 |
| Prolif. A.Macs | Pds5a | Prolif. A.Macs | Gas2l3 | Prolif. A.Macs | Smarcc1 |
| Prolif. A.Macs | Pold1 | Prolif. A.Macs | Nxt1 | Prolif. A.Macs | Rfc1 |
| Prolif. A.Macs | Ipo5 | Prolif. A.Macs | Nudc | Prolif. A.Macs | Ccdc25 |
| Prolif. A.Macs | Ncaph2 | Prolif. A.Macs | Pole3 | Prolif. A.Macs | Alad |
| Prolif. A.Macs | Nsmce4a | Prolif. A.Macs | Terf1 | Prolif. A.Macs | Ppp1cc |
| Prolif. A.Macs | Hnrnpa3 | Prolif. A.Macs | Anapc5 | Prolif. A.Macs | Pgp |
| Prolif. A.Macs | Odf2 | Prolif. A.Macs | Plp2 | Prolif. A.Macs | Hnrnpa2b1 |
| Prolif. A.Macs | Bub3 | Prolif. A.Macs | Ctsk | Prolif. A.Macs | Rbm3 |
| Prolif. A.Macs | Tbc1d31 | Prolif. A.Macs | Cln6 | Prolif. A.Macs | Rfc2 |

**Supplementary Table 1: Cluster defining genes in scRNAseq**

| | |
|---|---|
| Prolif. A.Macs | Hnrnpf |
| Prolif. A.Macs | Samd1 |
| Prolif. A.Macs | Nfia |
| Prolif. A.Macs | Sephs1 |
| Prolif. A.Macs | Rpa1 |
| Prolif. A.Macs | Serinc3 |
| Prolif. A.Macs | Timm50 |
| Prolif. A.Macs | Lsm8 |
| Prolif. A.Macs | Pih1d1 |
| Prolif. A.Macs | Fundc2 |
| Prolif. A.Macs | Eea1 |
| Prolif. A.Macs | Eif1ad |
| Prolif. A.Macs | Dynlt1f |
| Prolif. A.Macs | Suz12 |
| Prolif. A.Macs | E2f1 |
| Prolif. A.Macs | Hnrnpul1 |
| Prolif. A.Macs | Supt16 |
| Prolif. A.Macs | Hnrnpu |
| Prolif. A.Macs | Uchl5 |
| Prolif. A.Macs | Actn4 |
| Prolif. A.Macs | Arl6ip6 |
| Prolif. A.Macs | Snrpe |
| Prolif. A.Macs | Marco |
| Prolif. A.Macs | Tuba1c |
| Prolif. A.Macs | Uqcr10 |
| Prolif. A.Macs | Naa50 |
| Prolif. A.Macs | Lnp |
| Prolif. A.Macs | Snrpb |
| Prolif. A.Macs | Tpm4 |
| Prolif. A.Macs | Txn1 |
| Prolif. A.Macs | Sep-10 |
| Prolif. A.Macs | Mxi1 |
| Prolif. A.Macs | Carnmt1 |
| Prolif. A.Macs | Psmg2 |
| Prolif. A.Macs | Magoh |
| Prolif. A.Macs | Phf5a |
| Prolif. A.Macs | Ubald2 |
| Prolif. A.Macs | Nop56 |
| Prolif. A.Macs | Numa1 |
| Prolif. A.Macs | Eny2 |
| Prolif. A.Macs | Pbdc1 |
| Prolif. A.Macs | Cisd1 |
| Prolif. A.Macs | Tmco1 |
| Prolif. A.Macs | Calm3 |
| Prolif. A.Macs | Nelfe |
| Prolif. A.Macs | Kpna3 |
| Prolif. A.Macs | Slc29a1 |
| Prolif. A.Macs | Bckdk |
| Prolif. A.Macs | Rexo2 |
| Prolif. A.Macs | B930036N10Rik |
| Prolif. A.Macs | Rpl22l1 |
| Prolif. A.Macs | Erh |
| Prolif. A.Macs | Stag1 |
| Prolif. A.Macs | Lyar |
| Prolif. A.Macs | Naa20 |
| Prolif. A.Macs | Nono |
| Prolif. A.Macs | Serbp1 |
| Prolif. A.Macs | Mif4gd |
| Prolif. A.Macs | Iws1 |
| Prolif. A.Macs | Pbx3 |
| Prolif. A.Macs | Cdkn2d |
| Prolif. A.Macs | Smchd1 |
| Prolif. A.Macs | Jade1 |
| Prolif. A.Macs | Dnajc2 |
| Prolif. A.Macs | Mrps17 |
| Prolif. A.Macs | Cbx1 |
| Prolif. A.Macs | Ccng2 |
| Prolif. A.Macs | Rbbp8 |
| Prolif. A.Macs | Hnrnpa1 |
| Prolif. A.Macs | Casp8ap2 |
| Prolif. A.Macs | Rdx |
| Prolif. A.Macs | Luc7l3 |
| Prolif. A.Macs | Tbc1d15 |
| Prolif. A.Macs | Atp5g3 |
| Prolif. A.Macs | Nde1 |
| Prolif. A.Macs | Ndufaf2 |
| Prolif. A.Macs | Ddx39 |
| Prolif. A.Macs | Mapre1 |
| Prolif. A.Macs | Taf15 |
| Prolif. A.Macs | Snrnp25 |
| Prolif. A.Macs | Rbbp4 |
| Prolif. A.Macs | Ddx27 |
| Prolif. A.Macs | Taf1 |
| Prolif. A.Macs | Ilf2 |
| Prolif. A.Macs | Reep4 |
| Prolif. A.Macs | Set |
| Prolif. A.Macs | Ipo9 |
| Prolif. A.Macs | Rnaseh2c |
| Prolif. A.Macs | Luc7l2 |
| Prolif. A.Macs | Matr3 |

**Supplementary Table 1: Cluster defining genes in scRNAseq**

| | | | | | |
|---|---|---|---|---|---|
| Prolif. A.Macs | Trp53 | Prolif. A.Macs | Mfsd10 | Prolif. A.Macs | Cav2 |
| Prolif. A.Macs | Hist1h4i | Prolif. A.Macs | Tmem245 | Prolif. A.Macs | Srsf1 |
| Prolif. A.Macs | Ctdnep1 | Prolif. A.Macs | Brd9 | Prolif. A.Macs | Plscr1 |
| Prolif. A.Macs | Arpp19 | Prolif. A.Macs | Vim | Prolif. A.Macs | Hp1bp3 |
| Prolif. A.Macs | Mthfd2 | Prolif. A.Macs | Casp7 | Prolif. A.Macs | Abcd2 |
| Prolif. A.Macs | Stub1 | Prolif. A.Macs | Tcerg1 | Prolif. A.Macs | Emsy |
| Prolif. A.Macs | Hprt | Prolif. A.Macs | Atf1 | Prolif. A.Macs | Med30 |
| Prolif. A.Macs | 1110059E24Rik | Prolif. A.Macs | Snrpg | Prolif. A.Macs | Gyg |
| Prolif. A.Macs | Myg1 | Prolif. A.Macs | Gtf2a2 | Prolif. A.Macs | Pank2 |
| Prolif. A.Macs | Cnot6 | Prolif. A.Macs | Ptgfrn | Prolif. A.Macs | Dna2 |
| Prolif. A.Macs | Vars | Prolif. A.Macs | Nup153 | Prolif. A.Macs | Mrpl51 |
| Prolif. A.Macs | Tada1 | Prolif. A.Macs | Cnot6l | Prolif. A.Macs | Tusc3 |
| Prolif. A.Macs | Fam96a | Prolif. A.Macs | Taf6 | Prolif. A.Macs | Thoc7 |
| Prolif. A.Macs | Ltn1 | Prolif. A.Macs | Nudt5 | Prolif. A.Macs | Smarcad1 |
| Prolif. A.Macs | Oat | Prolif. A.Macs | Upf3b | Prolif. A.Macs | Cdc123 |
| Prolif. A.Macs | Pole4 | Prolif. A.Macs | Mtch1 | Prolif. A.Macs | Cpsf6 |
| Prolif. A.Macs | Btf3l4 | Prolif. A.Macs | H2afy | Prolif. A.Macs | Azin1 |
| Prolif. A.Macs | Hnrnpm | Prolif. A.Macs | Cd164 | Prolif. A.Macs | Srsf10 |
| Prolif. A.Macs | Ppp1r7 | Prolif. A.Macs | Atp5g2 | Prolif. A.Macs | Khsrp |
| Prolif. A.Macs | AW112010 | Prolif. A.Macs | Sp1 | Prolif. A.Macs | Zfp91 |
| Prolif. A.Macs | Lin37 | Prolif. A.Macs | C330007P06Rik | Prolif. A.Macs | Pcbd2 |
| Prolif. A.Macs | Tardbp | Prolif. A.Macs | Pbrm1 | Prolif. A.Macs | Tsen34 |
| Prolif. A.Macs | Megf9 | Prolif. A.Macs | Ebp | Prolif. A.Macs | Npm1 |
| Prolif. A.Macs | Ptges3 | Prolif. A.Macs | Tmem128 | Prolif. A.Macs | Ube2e3 |
| Prolif. A.Macs | Rnps1 | Prolif. A.Macs | Adss | Prolif. A.Macs | Kpnb1 |
| Prolif. A.Macs | Cspp1 | Prolif. A.Macs | Sep-07 | Prolif. A.Macs | Rpia |
| Prolif. A.Macs | Setd8 | Prolif. A.Macs | Ubl4a | Prolif. A.Macs | Tuba4a |
| Prolif. A.Macs | Uba2 | Prolif. A.Macs | Nup62 | Prolif. A.Macs | Atp13a3 |
| Prolif. A.Macs | Mrpl18 | Prolif. A.Macs | Atp6v1c1 | Prolif. A.Macs | Rbmxl1 |
| Prolif. A.Macs | Zdhhc21 | Prolif. A.Macs | Mtx2 | Prolif. A.Macs | Pnpla8 |

**Supplementary Table 1: Cluster defining genes in scRNAseq**

| | |
|---|---|
| Prolif. A.Macs | Nenf |
| Prolif. A.Macs | Hspa14 |
| Prolif. A.Macs | Mtch2 |
| Prolif. A.Macs | Aebp2 |
| Prolif. A.Macs | Ywhaq |
| Prolif. A.Macs | Ssb |
| Prolif. A.Macs | Sf3b2 |
| Prolif. A.Macs | Tceb1 |
| Prolif. A.Macs | Cxcr4 |
| Prolif. A.Macs | Vrk1 |
| Prolif. A.Macs | Brd3 |
| Prolif. A.Macs | Gspt1 |
| Prolif. A.Macs | Tfec |
| Prolif. A.Macs | Eif4a3 |
| Prolif. A.Macs | Commd3 |
| Prolif. A.Macs | Cisd2 |
| Prolif. A.Macs | Il18 |
| Prolif. A.Macs | Ngfrap1 |
| Prolif. A.Macs | Gatad1 |
| Prolif. A.Macs | Gpx4 |
| Prolif. A.Macs | Ehmt2 |
| Prolif. A.Macs | Pcyox1 |
| Prolif. A.Macs | Ppid |
| Prolif. A.Macs | Hccs |
| Prolif. A.Macs | Uhrf2 |
| Prolif. A.Macs | Atp5o |
| Prolif. A.Macs | Cox7a2 |
| Prolif. A.Macs | Rheb |
| Prolif. A.Macs | Plgrkt |
| Prolif. A.Macs | Ginm1 |
| Prolif. A.Macs | Ddx39b |
| Prolif. A.Macs | G3bp1 |
| Prolif. A.Macs | Snrpd2 |
| Prolif. A.Macs | Rps27l |
| Prolif. A.Macs | U2af2 |
| Prolif. A.Macs | Cnih1 |
| Prolif. A.Macs | Nup210 |
| Prolif. A.Macs | Vkorc1 |
| Prolif. A.Macs | Acat1 |
| Prolif. A.Macs | Hnrnpd |
| Prolif. A.Macs | Phip |
| Prolif. A.Macs | Pcmt1 |
| Prolif. A.Macs | Sf3b5 |
| Prolif. A.Macs | Larp4b |
| Prolif. A.Macs | Ergic2 |
| Prolif. A.Macs | Tsn |
| Prolif. A.Macs | Umps |
| Prolif. A.Macs | Tgm2 |
| Prolif. A.Macs | Cd302 |
| Prolif. A.Macs | Tbc1d10b |
| Prolif. A.Macs | Txnl1 |
| Prolif. A.Macs | Vkorc1l1 |
| Prolif. A.Macs | Ncl |
| Prolif. A.Macs | Cyb5r3 |
| Prolif. A.Macs | Vdac3 |
| Prolif. A.Macs | Cdv3 |
| Prolif. A.Macs | Tra2b |
| Prolif. A.Macs | C530008M17Rik |
| Prolif. A.Macs | U2surp |
| Prolif. A.Macs | Pnp |
| Prolif. A.Macs | Coro1c |
| Prolif. A.Macs | Ndufab1 |
| Prolif. A.Macs | Dhrs7b |
| Prolif. A.Macs | Htatsf1 |
| Prolif. A.Macs | Ptbp3 |
| Prolif. A.Macs | Vdac1 |
| Prolif. A.Macs | Acadl |
| Prolif. A.Macs | Ndufa5 |
| Prolif. A.Macs | Lsm6 |
| Prolif. A.Macs | Maz |
| Prolif. A.Macs | Nrip1 |
| Prolif. A.Macs | Sf3b6 |
| Prolif. A.Macs | Snrpf |
| Prolif. A.Macs | M6pr |
| Prolif. A.Macs | Srsf2 |
| Prolif. A.Macs | Cox6b2 |
| Prolif. A.Macs | Mfsd1 |
| Prolif. A.Macs | Mrpl42 |
| Prolif. A.Macs | Pum2 |
| Prolif. A.Macs | Cox5b |
| Prolif. A.Macs | Rer1 |
| Prolif. A.Macs | Cd200r4 |
| Prolif. A.Macs | Ssbp1 |
| Prolif. A.Macs | Cox5a |
| Prolif. A.Macs | Yeats4 |
| Prolif. A.Macs | Ewsr1 |
| Prolif. A.Macs | Bcas2 |
| Prolif. A.Macs | Ppm1g |
| Prolif. A.Macs | Utp3 |
| Prolif. A.Macs | Meaf6 |

**Supplementary Table 1: Cluster defining genes in scRNAseq**

| | |
|---|---|
| Prolif. A.Macs | Cklf |
| Prolif. A.Macs | Ugp2 |
| Prolif. A.Macs | Clic4 |
| Prolif. A.Macs | Ccni |
| Prolif. A.Macs | Mapk3 |
| Prolif. A.Macs | Med28 |
| Prolif. A.Macs | Clic1 |
| Prolif. A.Macs | Eif4g2 |
| Prolif. A.Macs | Etfa |
| Prolif. A.Macs | Minos1 |
| Prolif. A.Macs | Idh2 |
| Prolif. A.Macs | Bod1l |
| Prolif. A.Macs | Eif3c |
| Prolif. A.Macs | Rbm8a |
| Prolif. A.Macs | Ppig |
| Prolif. A.Macs | Hnrnpa0 |
| Prolif. A.Macs | Mrpl14 |
| Prolif. A.Macs | Pcbp1 |
| Prolif. A.Macs | Mat2a |
| Prolif. A.Macs | Polr2k |
| Prolif. A.Macs | Mdh1 |
| Prolif. A.Macs | Lmo4 |
| Prolif. A.Macs | Fus |
| Prolif. A.Macs | Chil3 |
| Prolif. A.Macs | Metap2 |
| Prolif. A.Macs | Car4 |
| Prolif. A.Macs | Polr2f |
| Prolif. A.Macs | Lsm4 |
| Prolif. A.Macs | Mrfap1 |
| Prolif. A.Macs | Vps35 |
| Prolif. A.Macs | Tgfbi |
| Prolif. A.Macs | Gm42418 |
| cDC1 | Ifi205 |
| cDC1 | Xcr1 |
| cDC1 | Sep-03 |
| cDC1 | Itgae |
| cDC1 | Amica1 |
| cDC1 | Cadm1 |
| cDC1 | Ckb |
| cDC1 | Cxx1a |
| cDC1 | Gcsam |
| cDC1 | Clec9a |
| cDC1 | Hepacam2 |
| cDC1 | Cd207 |
| cDC1 | Btla |
| cDC1 | A530099J19Rik |
| cDC1 | Qpct |
| cDC1 | Klrb1b |
| cDC1 | Fcrla |
| cDC1 | Rab7b |
| cDC1 | Plbd1 |
| cDC1 | Cxx1b |
| cDC1 | H2-Ob |
| cDC1 | Tlr3 |
| cDC1 | St3gal5 |
| cDC1 | Fgd2 |
| cDC1 | Grap |
| cDC1 | Cystm1 |
| cDC1 | Tnni2 |
| cDC1 | Sult1a1 |
| cDC1 | Pianp |
| cDC1 | Irf8 |
| cDC1 | Mnda |
| cDC1 | Cst3 |
| cDC1 | Gm11837 |
| cDC1 | Sep-06 |
| cDC1 | Anpep |
| cDC1 | Cxcr3 |
| cDC1 | Flt3 |
| cDC1 | Ccr2 |
| cDC1 | H2-DMb1 |
| cDC1 | BC028528 |
| cDC1 | Cldn1 |
| cDC1 | Kit |
| cDC1 | Cd24a |
| cDC1 | Gatm |
| cDC1 | Itgb7 |
| cDC1 | Dbn1 |
| cDC1 | Id2 |
| cDC1 | F2rl2 |
| cDC1 | H2-Eb1 |
| cDC1 | Fam149a |
| cDC1 | Slamf7 |
| cDC1 | H2-Aa |
| cDC1 | Cbfa2t3 |
| cDC1 | Cyp8b1 |
| cDC1 | P2ry10 |
| cDC1 | H2-Ab1 |
| cDC1 | Alcam |
| cDC1 | Aif1 |
| cDC1 | Pak1 |
| cDC1 | Ppm1m |
| cDC1 | Naaa |
| cDC1 | Fgl2 |
| cDC1 | Ccnd1 |
| cDC1 | Naalad2 |
| cDC1 | Cd74 |
| cDC1 | Ass1 |
| cDC1 | Hmgn3 |
| cDC1 | Timeless |
| cDC1 | Crip1 |
| cDC1 | Kctd14 |
| cDC1 | Fchsd2 |
| cDC1 | H2-DMa |
| cDC1 | Marcks |
| cDC1 | Mycl |
| cDC1 | Sub1 |
| cDC1 | Cd52 |
| cDC1 | Wdfy4 |
| cDC1 | Tbc1d4 |
| cDC1 | Cfh |
| cDC1 | Gpr82 |

**Supplementary Table 1: Cluster defining genes in scRNAseq**

| | | | | | |
|---|---|---|---|---|---|
| cDC1 | Ltb4r1 | cDC1 | H2-Oa | cDC1 | Pvrl1 |
| cDC1 | Snx22 | cDC1 | 1700025G04Rik | cDC1 | Syngr2 |
| cDC1 | Nccrp1 | cDC1 | Ciita | cDC1 | Tap1 |
| cDC1 | Cxcl16 | cDC1 | Apobec3 | cDC1 | Olfm1 |
| cDC1 | Anxa6 | cDC1 | Kmo | cDC1 | Cd180 |
| cDC1 | Xlr | cDC1 | Rpl17 | cDC1 | Commd8 |
| cDC1 | Lpcat1 | cDC1 | Pglyrp1 | cDC1 | Rgs18 |
| cDC1 | Cd86 | cDC1 | Psmb8 | cDC1 | Itpr1 |
| cDC1 | Timd4 | cDC1 | Rpl3 | cDC1 | Gng10 |
| cDC1 | Rbpj | cDC1 | Lmo1 | cDC1 | Tmsb10 |
| cDC1 | Mndal | cDC1 | Gsn | cDC1 | Epcam |
| cDC1 | Dynlt1a | cDC1 | Tspan13 | cDC1 | Psme1 |
| cDC1 | Slc46a3 | cDC1 | Rps11 | cDC1 | Adam8 |
| cDC1 | Arhgap5 | cDC1 | Rnase6 | cDC1 | Kctd12 |
| cDC1 | Ap1s3 | cDC1 | Txndc15 | cDC1 | Rps7 |
| cDC1 | Ece1 | cDC1 | Tbc1d8 | cDC1 | Napsa |
| cDC1 | Klrd1 | cDC1 | Arhgap18 | cDC1 | Rpl21 |
| cDC1 | Ryr1 | cDC1 | Rasgrp4 | cDC1 | Endod1 |
| cDC1 | Cysltr1 | cDC1 | Tspan33 | cDC1 | Rpl27a |
| cDC1 | Siglecg | cDC1 | Rps6 | cDC1 | Rps15a |
| cDC1 | Batf3 | cDC1 | Fgfr1 | cDC1 | BC035044 |
| cDC1 | Atox1 | cDC1 | Irf5 | cDC1 | Efhd2 |
| cDC1 | Rgs10 | cDC1 | Lat2 | cDC1 | Rpl13 |
| cDC1 | Plpp1 | cDC1 | Tmsb4x | cDC1 | Arhgap15 |
| cDC1 | P3h2 | cDC1 | Clec12a | cDC1 | Rpl23a |
| cDC1 | Plekho1 | cDC1 | Rab43 | cDC1 | Gng12 |
| cDC1 | Gpr68 | cDC1 | Basp1 | cDC1 | Ppfia4 |
| cDC1 | Ifi203 | cDC1 | Dbnl | cDC1 | Rps4x |
| cDC1 | Psmb9 | cDC1 | Ngfrap1 | cDC1 | Phf11b |
| cDC1 | Ppt1 | cDC1 | Slamf8 | cDC1 | Pafah1b3 |
| cDC1 | Plekha5 | cDC1 | Gpr171 | cDC1 | Eps8 |
| cDC1 | Dpp4 | cDC1 | 2310040G24Rik | cDC1 | Actr3 |
| cDC1 | Cnn2 | cDC1 | Runx3 | cDC1 | Cd244 |
| cDC1 | Nsmaf | cDC1 | Rplp0 | cDC1 | Rpl4 |
| cDC1 | Gm2a | cDC1 | Ptpn18 | cDC1 | Rps18 |
| cDC1 | Naga | cDC1 | Itpripl1 | cDC1 | Ppp1r11 |
| cDC1 | Slco3a1 | cDC1 | Lmnb1 | cDC1 | Rps24 |
| cDC1 | Arsb | cDC1 | AA467197 | cDC1 | Hiat1 |
| cDC1 | Rpl14 | cDC1 | Rps19 | cDC1 | Rpl9 |
| cDC1 | Plp2 | cDC1 | Rps5 | cDC1 | Rpl19 |
| cDC1 | Fmnl2 | cDC1 | Sh3bgrl3 | cDC1 | Fnbp1l |
| cDC1 | Zbtb46 | cDC1 | Bmyc | cDC1 | Rpl11 |
| cDC1 | Rpl18a | cDC1 | Rps29 | cDC1 | Tiam1 |

**Supplementary Table 1: Cluster defining genes in scRNAseq**

| | | | | | |
|---|---|---|---|---|---|
| cDC1 | Rps23 | cDC1 | Rpl13a | cDC1 | Supt20 |
| cDC1 | Lsp1 | cDC1 | Slc25a38 | cDC1 | Got2 |
| cDC1 | Evi2a | cDC1 | Spc24 | cDC1 | Arpc5l |
| cDC1 | Bloc1s2 | cDC1 | Pkp3 | cDC1 | Eef1g |
| cDC1 | Rps27 | cDC1 | Ly6e | cDC1 | Cd37 |
| cDC1 | Rps15 | cDC1 | Ifngr1 | cDC1 | Sp100 |
| cDC1 | Ccl17 | cDC1 | Psap | cDC1 | Phactr2 |
| cDC1 | Pomp | cDC1 | Cdk14 | cDC1 | Rps25 |
| cDC1 | Nav1 | cDC1 | Cd81 | cDC1 | Clic1 |
| cDC1 | Pkib | cDC1 | Ahr | cDC1 | Adgre5 |
| cDC1 | Tbc1d10c | cDC1 | Tes | cDC1 | Tpm3 |
| cDC1 | Haao | cDC1 | Fyn | cDC1 | Slc31a2 |
| cDC1 | Stx17 | cDC1 | Rpl18 | cDC1 | Nfkbie |
| cDC1 | Stard3nl | cDC1 | Camk1d | cDC1 | Rpl27 |
| cDC1 | Rtn1 | cDC1 | Rps3 | cDC1 | Mctp1 |
| cDC1 | Tap2 | cDC1 | Acot7 | cDC1 | Mbnl1 |
| cDC1 | Rpl37 | cDC1 | Spint2 | cDC1 | Acvrl1 |
| cDC1 | Tubb2a | cDC1 | Ccdc12 | cDC1 | Cyb5a |
| cDC1 | Gna15 | cDC1 | Bri3bp | cDC1 | Rpl23a-ps3 |
| cDC1 | Ptpn22 | cDC1 | Lemd2 | cDC1 | Hmha1 |
| cDC1 | Rps3a1 | cDC1 | Rpl6 | cDC1 | Rpl10a |
| cDC1 | H2-Q7 | cDC1 | H2-M3 | cDC1 | Ptms |
| cDC1 | Mthfd2 | cDC1 | Tnfrsf13b | cDC1 | Arl4c |
| cDC1 | St8sia4 | cDC1 | H2-DMb2 | cDC1 | Mar-01 |
| cDC1 | Sigmar1 | cDC1 | Cmtm7 | cDC1 | Inpp5d |
| cDC1 | Rpl32 | cDC1 | Tifa | cDC1 | Pea15a |
| cDC1 | Hfe | cDC1 | Rps14 | cDC1 | Rpl34 |
| cDC1 | Eef1b2 | cDC1 | Hspb11 | cDC1 | Traf1 |
| cDC1 | Rpsa | cDC1 | H2afy | cDC1 | Mef2c |
| cDC1 | Pstpip1 | cDC1 | Dek | cDC1 | Hnrnpa1 |
| cDC1 | Rnf145 | cDC1 | Gng2 | cDC1 | Csrp1 |
| cDC1 | Cnp | cDC1 | Hps4 | cDC1 | Gas7 |
| cDC1 | Rps13 | cDC1 | Mrpl52 | cDC1 | Rps2 |
| cDC1 | Tuba1a | cDC1 | Samd9l | cDC1 | Gm11808 |
| cDC1 | Gpr183 | cDC1 | Ddit4 | cDC1 | Rpl35a |
| cDC1 | Rps28 | cDC1 | Sdf2l1 | cDC1 | Fcgr2b |
| cDC1 | Ubb | cDC1 | Gltp | cDC1 | Cdkn2c |
| cDC1 | Rps16 | cDC1 | Lbh | cDC1 | Fem1c |
| cDC1 | H2-Q6 | cDC1 | Fnbp1 | cDC1 | Pfkp |
| cDC1 | Gm8730 | cDC1 | Havcr2 | cDC1 | Kctd10 |
| cDC1 | Rogdi | cDC1 | Lpgat1 | cDC1 | Rpl13-ps3 |
| cDC1 | Sgsm3 | cDC1 | Klrk1 | cDC1 | Klhl6 |
| cDC1 | Sh3bp1 | cDC1 | Coro7 | cDC1 | Anp32e |

**Supplementary Table 1: Cluster defining genes in scRNAseq**

| | | | | | |
|---|---|---|---|---|---|
| cDC1 | Gdi2 | cDC1 | Tmpo | cDC1 | Gm26740 |
| cDC1 | Tubb6 | cDC1 | Gltscr2 | cDC1 | Rala |
| cDC1 | Gm9493 | cDC1 | Impdh2 | cDC1 | Ptma |
| cDC1 | Hps3 | cDC1 | Vrk1 | cDC1 | Snrpa |
| cDC1 | Coro1a | cDC1 | Susd3 | cDC1 | Slc1a5 |
| cDC1 | Twf2 | cDC1 | Tnfaip8 | cDC1 | Dnajc8 |
| cDC1 | Ahcyl2 | cDC1 | Srebf2 | cDC1 | Acsl5 |
| cDC1 | Fau | cDC1 | Rpl36 | cDC1 | Ezh2 |
| cDC1 | Prkd3 | cDC1 | Nudc | cDC1 | Cct5 |
| cDC1 | Ppib | cDC1 | Cdc37 | cDC1 | Rbbp7 |
| cDC1 | Lrrk2 | cDC1 | Pbx1 | cDC1 | Rpl15 |
| cDC1 | Dctpp1 | cDC1 | Skap2 | cDC1 | Eif2s1 |
| cDC1 | Snx3 | cDC1 | Rpl39 | cDC1 | Irf9 |
| cDC1 | Top2a | cDC1 | Cdk1 | cDC1 | Card19 |
| cDC1 | Ccr5 | cDC1 | Acer3 | cDC1 | Pole4 |
| cDC1 | Ncbp3 | cDC1 | Tbrg1 | cDC1 | Psip1 |
| cDC1 | H2afx | cDC1 | Rps27rt | cDC1 | Rpl22 |
| cDC1 | Antxr2 | cDC1 | Tmem50b | cDC1 | Stx16 |
| cDC1 | Sema4d | cDC1 | Ctnnbip1 | cDC1 | Rab11a |
| cDC1 | Ifi35 | cDC1 | Ncoa7 | cDC1 | Oxct1 |
| cDC1 | Vrk2 | cDC1 | Isyna1 | cDC1 | Efr3a |
| cDC1 | Ncf4 | cDC1 | Mdh1 | cDC1 | Zfp318 |
| cDC1 | Birc5 | cDC1 | Rpl26 | cDC1 | Set |
| cDC1 | Pqlc1 | cDC1 | 4930523C07Rik | cDC1 | P2ry6 |
| cDC1 | Hexb | cDC1 | Il13ra1 | cDC1 | Gm10036 |
| cDC1 | Anapc15 | cDC1 | Nsun2 | cDC1 | Eif3f |
| cDC1 | Dnaja1 | cDC1 | Uba52 | cDC1 | Plekho2 |
| cDC1 | Wasf2 | cDC1 | Fbl | cDC1 | Nop58 |
| cDC1 | Map4k1 | cDC1 | Rpl9-ps6 | cDC1 | Trim35 |
| cDC1 | Selplg | cDC1 | Eif3g | cDC1 | 5031439G07Rik |
| cDC1 | Atpif1 | cDC1 | Hcls1 | cDC1 | Sod1 |
| cDC1 | Gm9844 | cDC1 | Gm6133 | cDC1 | Lsm6 |
| cDC1 | Cnbp | cDC1 | Pdlim2 | cDC1 | Spint1 |
| cDC1 | Jak2 | cDC1 | Anxa1 | cDC1 | Agpat3 |
| cDC1 | Zyx | cDC1 | Ccdc107 | cDC1 | Vars |
| cDC1 | Zfp422 | cDC1 | Gm6377 | cDC1 | Dapp1 |
| cDC1 | Sms | cDC1 | Park7 | cDC1 | Ldha |
| cDC1 | Ctnna1 | cDC1 | Cks1b | cDC1 | Cd48 |
| cDC1 | Rpl8 | cDC1 | Tpm4 | cDC1 | Tyk2 |
| cDC1 | Rpl28 | cDC1 | Pla2g16 | cDC1 | Mcm6 |
| cDC1 | Aph1c | cDC1 | Hmgn1 | cDC1 | Ncoa3 |
| cDC1 | Mpp6 | cDC1 | Pmaip1 | cDC1 | Znrd1 |
| cDC1 | Rpl6l | cDC1 | Sh2b3 | cDC1 | Rraga |

**Supplementary Table 1: Cluster defining genes in scRNAseq**

| | |
|---|---|
| cDC1 | Dennd1b |
| cDC1 | Pdia3 |
| cDC1 | Srsf7 |
| cDC1 | Prpf19 |
| cDC1 | Cycs |
| cDC1 | Rgs2 |
| cDC1 | Ffar4 |
| cDC1 | Psme2 |
| cDC1 | Npm3 |
| cDC1 | Ndufa6 |
| cDC1 | Npm1 |
| cDC1 | Eif3h |
| cDC1 | Rtn4 |
| cDC1 | Chchd3 |
| cDC1 | Glipr1 |
| cDC1 | Rbm17 |
| cDC1 | Rpl5 |
| cDC1 | Rps12-ps3 |
| cDC1 | St3gal4 |
| cDC1 | Rassf2 |
| cDC1 | Trim28 |
| cDC1 | Exosc8 |
| cDC1 | Fundc1 |
| cDC1 | Arpc2 |
| cDC1 | 2810417H13Rik |
| cDC1 | Gfer |
| cDC1 | Arf5 |
| cDC1 | Rpl30 |
| cDC1 | Pura |
| cDC1 | Unc93b1 |
| cDC1 | Rps12 |
| cDC1 | Hspa1a |
| cDC1 | Sgpp1 |
| cDC1 | Tmem55b |
| cDC1 | Cct7 |
| cDC1 | Tnpo3 |
| cDC1 | Igbp1 |
| cDC1 | Cd82 |
| cDC1 | Rnaset2b |
| cDC1 | Limd1 |
| cDC1 | Srsf3 |
| cDC1 | Polr1d |
| cDC1 | Man2b1 |
| cDC1 | Srsf11 |
| cDC1 | Sptssa |
| cDC1 | Dynll2 |
| cDC1 | 0610030E20Rik |
| cDC1 | U2af1 |
| cDC1 | Itm2c |
| cDC1 | Pdrg1 |
| cDC1 | B230219D22Rik |
| cDC1 | Jup |
| cDC1 | Snrpf |
| cDC1 | Cmtm6 |
| cDC1 | Rpl7a |
| cDC1 | H13 |
| cDC1 | Trim30a |
| cDC1 | Uvrag |
| cDC1 | Rpl29 |
| cDC1 | Sepw1 |
| cDC1 | Dna2 |
| cDC1 | 2700094K13Rik |
| cDC1 | Fam105a |
| cDC1 | Rapgef6 |
| cDC1 | Ahnak |
| cDC1 | Smc4 |
| cDC1 | Atp5c1 |
| cDC1 | Khdrbs1 |
| cDC1 | Thoc7 |
| cDC1 | Eif3k |
| cDC1 | Notch2 |
| cDC1 | Eif3e |
| cDC1 | Sec11c |
| cDC1 | Rinl |
| cDC1 | Ube2s |
| cDC1 | Hspe1 |
| cDC1 | Rpl27-ps3 |
| cDC1 | Gm10073 |
| cDC1 | 2700060E02Rik |
| cDC1 | Nucks1 |
| cDC1 | Anp32b |
| cDC1 | Smc2 |
| cDC1 | Rab32 |
| cDC1 | H2afv |
| cDC1 | Stmn1 |
| cDC1 | Cd83 |
| cDC1 | Hmgb2 |
| cDC1 | H2afz |
| cDC1 | Btg2 |
| cDC1 | Hmgb1 |
| cDC1 | Tubb5 |

| | |
|---|---|
| cDC2 | Cd209a |
| cDC2 | Tnip3 |
| cDC2 | Mgl2 |
| cDC2 | Tmem176b |
| cDC2 | H2-DMb2 |
| cDC2 | Ccnd1 |
| cDC2 | S100a6 |
| cDC2 | Tmem176a |
| cDC2 | Rgs18 |
| cDC2 | Tnfsf9 |
| cDC2 | Klrd1 |
| cDC2 | Il1rl1 |
| cDC2 | H2-Aa |
| cDC2 | H2-Oa |
| cDC2 | Slamf7 |
| cDC2 | H2-Eb1 |
| cDC2 | Cd74 |
| cDC2 | S100a4 |
| cDC2 | Cd209d |
| cDC2 | Bcl11a |
| cDC2 | Socs3 |
| cDC2 | Ank |
| cDC2 | H2-Ab1 |
| cDC2 | H2-DMb1 |
| cDC2 | Clec10a |
| cDC2 | Kmo |
| cDC2 | Fgfr1 |
| cDC2 | Upb1 |
| cDC2 | Ly86 |
| cDC2 | Lmo1 |
| cDC2 | H2-DMa |
| cDC2 | Cd52 |
| cDC2 | Napsa |
| cDC2 | Cd72 |
| cDC2 | Bmyc |
| cDC2 | Amica1 |

**Supplementary Table 1: Cluster defining genes in scRNAseq**

| | | | | | |
|---|---|---|---|---|---|
| cDC2 | Cd7 | cDC2 | Acer3 | cDC2 | Ccr5 |
| cDC2 | Wfdc17 | cDC2 | Rpl3 | cDC2 | Clec4b1 |
| cDC2 | Crip1 | cDC2 | Evi2a | cDC2 | 1700025G04Rik |
| cDC2 | Cbfa2t3 | cDC2 | Abca9 | cDC2 | Gadd45b |
| cDC2 | Plbd1 | cDC2 | Rpl4 | cDC2 | Rps24 |
| cDC2 | Hepacam2 | cDC2 | Rps7 | cDC2 | Nfkbiz |
| cDC2 | Alcam | cDC2 | Rpl23a | cDC2 | Gng10 |
| cDC2 | Lat2 | cDC2 | Cysltr1 | cDC2 | Pak1 |
| cDC2 | Cst3 | cDC2 | H2afy | cDC2 | Flt3 |
| cDC2 | Marcks | cDC2 | Tnfrsf13b | cDC2 | Hfe |
| cDC2 | Gpr171 | cDC2 | Dpp4 | cDC2 | Pkp3 |
| cDC2 | Runx3 | cDC2 | Tuba1a | cDC2 | Acot7 |
| cDC2 | B4galt6 | cDC2 | Cfp | cDC2 | Sub1 |
| cDC2 | Rps27 | cDC2 | Kctd12 | cDC2 | Ifi27l2a |
| cDC2 | Lifr | cDC2 | Fcrls | cDC2 | Marcksl1 |
| cDC2 | Gm2a | cDC2 | Gpr132 | cDC2 | Sep-06 |
| cDC2 | Tspan13 | cDC2 | Fgd2 | cDC2 | Prkcb |
| cDC2 | Timp2 | cDC2 | Hexb | cDC2 | Rpl31 |
| cDC2 | Ptafr | cDC2 | Mef2c | cDC2 | Rpl11 |
| cDC2 | Ckb | cDC2 | Plekho1 | cDC2 | Rac2 |
| cDC2 | Cnn2 | cDC2 | Rps4x | cDC2 | Ddit4 |
| cDC2 | Ltb4r1 | cDC2 | Rbpj | cDC2 | Rps14 |
| cDC2 | Pid1 | cDC2 | Rpl6 | cDC2 | Rpl13a |
| cDC2 | Smim5 | cDC2 | Ccl4 | cDC2 | Cmtm7 |
| cDC2 | Ctsh | cDC2 | Aif1 | cDC2 | Limd2 |
| cDC2 | Fcgr2b | cDC2 | Rps15a | cDC2 | Lbr |
| cDC2 | Mmp12 | cDC2 | Ccl17 | cDC2 | Rpl17 |
| cDC2 | Gsn | cDC2 | Rtn1 | cDC2 | Rps13 |
| cDC2 | Lbh | cDC2 | H2-Ob | cDC2 | Arl2bp |
| cDC2 | Rps6 | cDC2 | Itgam | cDC2 | Trappc5 |
| cDC2 | Rpl18a | cDC2 | Syngr2 | cDC2 | Gm8730 |
| cDC2 | Qpct | cDC2 | Rpl19 | cDC2 | Pim1 |
| cDC2 | Ppfia4 | cDC2 | Rps29 | cDC2 | Rpl9 |
| cDC2 | Srgn | cDC2 | Rps5 | cDC2 | Rps27rt |
| cDC2 | Lsp1 | cDC2 | AA467197 | cDC2 | Rps19 |
| cDC2 | Tcf4 | cDC2 | Rps18 | cDC2 | Hspa1a |
| cDC2 | Itgb7 | cDC2 | Ccr2 | cDC2 | Rpl35a |
| cDC2 | Rpl14 | cDC2 | Rnase6 | cDC2 | Fau |
| cDC2 | Rpl13 | cDC2 | Etv3 | cDC2 | Rpl27a |
| cDC2 | Rplp0 | cDC2 | Gpr183 | cDC2 | Lrmp |
| cDC2 | Rps11 | cDC2 | Ifitm2 | cDC2 | Ms4a6c |
| cDC2 | Rpl21 | cDC2 | St3gal5 | cDC2 | Rps23 |
| cDC2 | Ifitm3 | cDC2 | Ciita | cDC2 | Eef1g |

**Supplementary Table 1: Cluster defining genes in scRNAseq**

| | | | | | |
|---|---|---|---|---|---|
| cDC2 | Rpl18 | cDC2 | Ccl9 | cDC2 | Sik1 |
| cDC2 | Rpl34 | cDC2 | Adgre5 | cDC2 | Rpl26 |
| cDC2 | Pdcd4 | cDC2 | Atox1 | cDC2 | Sell |
| cDC2 | St8sia4 | cDC2 | Stap1 | cDC2 | Cd300a |
| cDC2 | Arhgap15 | cDC2 | Clec12a | cDC2 | Fh1 |
| cDC2 | Ccrl2 | cDC2 | Bin1 | cDC2 | Adrbk2 |
| cDC2 | Rell1 | cDC2 | Cd180 | cDC2 | Gm11808 |
| cDC2 | Slamf8 | cDC2 | Itpr1 | cDC2 | Gm9493 |
| cDC2 | Knop1 | cDC2 | Klrk1 | cDC2 | Ptpn18 |
| cDC2 | Il1b | cDC2 | Rps25 | cDC2 | Ppp1r15a |
| cDC2 | Rpl10a | cDC2 | Cd48 | cDC2 | Eps8 |
| cDC2 | Rpl15 | cDC2 | Ifi30 | cDC2 | Fyn |
| cDC2 | Bloc1s2 | cDC2 | Fbrsl1 | cDC2 | Jak2 |
| cDC2 | Sh3bgrl3 | cDC2 | Ryr1 | cDC2 | Psmb8 |
| cDC2 | Traf1 | cDC2 | Foxp1 | cDC2 | Fam46a |
| cDC2 | Rps27a | cDC2 | Btg2 | cDC2 | Plxnd1 |
| cDC2 | Rps16 | cDC2 | Nfkbie | cDC2 | Id2 |
| cDC2 | Rpl36 | cDC2 | Rpl35 | cDC2 | Il2rg |
| cDC2 | Cxcl16 | cDC2 | Hpgd | cDC2 | Skint3 |
| cDC2 | BC035044 | cDC2 | Ccdc12 | cDC2 | Zfp36l1 |
| cDC2 | Tpm4 | cDC2 | Gng2 | cDC2 | Nav1 |
| cDC2 | Junb | cDC2 | Eif3f | cDC2 | Apobec3 |
| cDC2 | Pyhin1 | cDC2 | Bri3bp | cDC2 | Cdkn2d |
| cDC2 | Tspo | cDC2 | Batf3 | cDC2 | Cyth4 |
| cDC2 | Dbnl | cDC2 | Rpl37 | cDC2 | Rnaset2b |
| cDC2 | Rps3a1 | cDC2 | Arl3 | cDC2 | Rpl28 |
| cDC2 | Ccdc109b | cDC2 | Kdm2b | cDC2 | Psme1 |
| cDC2 | Stxbp6 | cDC2 | Scpep1 | cDC2 | Mrpl52 |
| cDC2 | Olfm1 | cDC2 | Ass1 | cDC2 | Rassf2 |
| cDC2 | Rpl6l | cDC2 | Pycard | cDC2 | Rpl7 |
| cDC2 | Sms | cDC2 | Rpl10 | cDC2 | Klrb1b |
| cDC2 | Nfkbia | cDC2 | Rps15 | cDC2 | Rpl29 |
| cDC2 | Rps28 | cDC2 | Nrros | cDC2 | Cdc42se2 |
| cDC2 | Cd244 | cDC2 | Irf5 | cDC2 | Arpc1b |
| cDC2 | Rpl32 | cDC2 | Eif4e3 | cDC2 | Park7 |
| cDC2 | 1600010M07Rik | cDC2 | Rpl27 | cDC2 | Uvrag |
| cDC2 | Zfp36 | cDC2 | Fgl2 | cDC2 | Pafah1b3 |
| cDC2 | Emb | cDC2 | Rpl23a-ps3 | cDC2 | Srebf2 |
| cDC2 | Tiam1 | cDC2 | Mbnl1 | cDC2 | Man1a |
| cDC2 | Tifa | cDC2 | Fyb | cDC2 | Zfhx3 |
| cDC2 | Arhgdib | cDC2 | Gas7 | cDC2 | Dek |
| cDC2 | Rogdi | cDC2 | Rps9 | cDC2 | Rpl24 |
| cDC2 | Tagap | cDC2 | Ptpro | cDC2 | Gnas |

**Supplementary Table 1: Cluster defining genes in scRNAseq**

| | | | | | |
|---|---|---|---|---|---|
| cDC2 | Bst2 | cDC2 | Pmaip1 | cDC2 | Ptger4 |
| cDC2 | Epcam | cDC2 | Selplg | cDC2 | Rpl9-ps6 |
| cDC2 | Chst12 | cDC2 | Tnfaip8 | cDC2 | H2-T23 |
| cDC2 | Myo5a | cDC2 | Ppib | cDC2 | Retnla |
| cDC2 | Gm9843 | cDC2 | Rps18-ps3 | cDC2 | Prkd3 |
| cDC2 | 0610007P14Rik | cDC2 | Pld4 | cDC2 | Arhgef6 |
| cDC2 | Stx7 | cDC2 | Uqcc2 | cDC2 | Sgpp1 |
| cDC2 | Emp3 | cDC2 | Hspb11 | cDC2 | Cox7a2l |
| cDC2 | Rgs10 | cDC2 | Clic1 | cDC2 | Sec11c |
| cDC2 | Smim14 | cDC2 | Gm10020 | cDC2 | Fbl |
| cDC2 | Rpl39 | cDC2 | Cd2ap | cDC2 | Arl4c |
| cDC2 | Cnbp | cDC2 | Ndufa6 | cDC2 | Kdm7a |
| cDC2 | Ifngr1 | cDC2 | Plac8 | cDC2 | Tnni2 |
| cDC2 | Ssr1 | cDC2 | Sigmar1 | cDC2 | Parp8 |
| cDC2 | Akr7a5 | cDC2 | Arl6ip5 | cDC2 | Prdx6 |
| cDC2 | Pstpip1 | cDC2 | Orai1 | cDC2 | Kdm6b |
| cDC2 | Atf3 | cDC2 | Rpl30 | cDC2 | Eno1 |
| cDC2 | BC028528 | cDC2 | Polr2e | cDC2 | Zfand5 |
| cDC2 | Eif3h | cDC2 | Rpl7a | cDC2 | Lactb |
| cDC2 | H13 | cDC2 | Ahnak | cDC2 | Sep-07 |
| cDC2 | Rnaset2a | cDC2 | Hmha1 | cDC2 | Dok2 |
| cDC2 | St3gal4 | cDC2 | Pepd | cDC2 | Itga4 |
| cDC2 | Card19 | cDC2 | Bhlhe40 | cDC2 | Hnrnpa1 |
| cDC2 | Coro1a | cDC2 | Cd24a | cDC2 | PISD |
| cDC2 | Plp2 | cDC2 | Fam96b | cDC2 | Anapc16 |
| cDC2 | Mpp6 | cDC2 | Nfkb1 | cDC2 | Mndal |
| cDC2 | Sdf2l1 | cDC2 | Pip4k2a | cDC2 | Dapp1 |
| cDC2 | Txn2 | cDC2 | Socs6 | cDC2 | Npm3 |
| cDC2 | H2-D1 | cDC2 | Hk2 | cDC2 | Eif3k |
| cDC2 | Rpl13-ps3 | cDC2 | Dnajc7 | cDC2 | Tmem55b |
| cDC2 | Zyx | cDC2 | Smdt1 | cDC2 | Gltp |
| cDC2 | Nfkbid | cDC2 | Tubb2a | cDC2 | Sepw1 |
| cDC2 | Psma7 | cDC2 | Ccdc88a | cDC2 | Siah2 |
| cDC2 | Slc29a3 | cDC2 | Uba52 | cDC2 | Sp110 |
| cDC2 | Rab43 | cDC2 | Ccnd3 | cDC2 | Ier5 |
| cDC2 | Dusp22 | cDC2 | Tmem173 | cDC2 | Map4k1 |
| cDC2 | Cnp | cDC2 | Rraga | cDC2 | Snrpf |
| cDC2 | Sema4d | cDC2 | Bmp2k | cDC2 | Polr3k |
| cDC2 | Sptssa | cDC2 | Rps26-ps1 | cDC2 | Rpgrip1 |
| cDC2 | Rps26 | cDC2 | Pacsin2 | cDC2 | Mt2 |
| cDC2 | Commd8 | cDC2 | H2-M3 | cDC2 | Nr4a1 |
| cDC2 | Efhd2 | cDC2 | Rel | cDC2 | Ccdc50 |
| cDC2 | Tep1 | cDC2 | Ywhah | cDC2 | Pebp1 |

**Supplementary Table 1: Cluster defining genes in scRNAseq**

| | |
|---|---|
| cDC2 | Hes6 |
| cDC2 | Tnpo3 |
| cDC2 | Cct5 |
| cDC2 | Oxct1 |
| cDC2 | Fem1c |
| cDC2 | Cd83 |
| cDC2 | Hspe1 |
| cDC2 | Csrnp1 |
| cDC2 | Rps6ka1 |
| cDC2 | Arf6 |
| cDC2 | Ly6c2 |
| cDC2 | Hist3h2a |
| cDC2 | Ifi35 |
| cDC2 | Gm6133 |
| cDC2 | Fdps |
| cDC2 | Crlf2 |
| cDC2 | Il13ra1 |
| cDC2 | Nol7 |
| cDC2 | Rbm22 |
| cDC2 | Eif3e |
| cDC2 | Lta4h |
| cDC2 | Polr2m |
| cDC2 | Capzb |
| cDC2 | Lmo2 |
| cDC2 | Khk |
| cDC2 | Cldnd1 |
| cDC2 | Nsa2 |
| cDC2 | Gltscr2 |
| cDC2 | Unc93b1 |
| cDC2 | Twf2 |
| cDC2 | Ehd4 |
| cDC2 | Phactr2 |
| cDC2 | Pabpc1 |
| cDC2 | Crlf3 |
| cDC2 | Ier2 |
| cDC2 | Ctnna1 |
| cDC2 | Higd2a |
| cDC2 | Mar-01 |
| cDC2 | Rps12-ps3 |
| cDC2 | Tsc22d1 |
| cDC2 | Macf1 |
| cDC2 | Oser1 |
| cDC2 | Prkcd |
| cDC2 | Jun |
| cDC2 | Mvb12a |
| cDC2 | Rad23b |
| cDC2 | Ly6e |
| cDC2 | H2-Q7 |
| cDC2 | Tmem59 |
| cDC2 | Rpl10-ps3 |
| cDC2 | Limd1 |
| cDC2 | Nek7 |
| cDC2 | Ifi203 |
| cDC2 | Sft2d2 |
| cDC2 | Slamf9 |
| cDC2 | Dguok |
| cDC2 | Gm10036 |
| cDC2 | Ywhaq |
| cDC2 | Egr1 |

**Supplementary Table 2**

| ID | log2FoldChange | Gene.name |
|---|---|---|
| **Downregulated in Cre+ (*Egr2* deficient)** | | |
| ENSMUSG00000050463 | -8.9238295 | Krt78 |
| ENSMUSG00000029377 | -8.8299607 | Ereg |
| ENSMUSG00000015665 | -8.4642595 | Awat1 |
| ENSMUSG00000061397 | -8.1662659 | Krt79 |
| ENSMUSG00000022096 | -7.7733436 | Hr |
| ENSMUSG00000046834 | -7.6160979 | Krt1 |
| ENSMUSG00000000359 | -7.3921633 | Rem1 |
| ENSMUSG00000025213 | -7.2194877 | Kazald1 |
| ENSMUSG00000017002 | -7.1935067 | Slpi |
| ENSMUSG00000052117 | -7.1539773 | D630039A03Rik |
| ENSMUSG00000035095 | -7.0438691 | Fam167a |
| ENSMUSG00000045394 | -6.8997025 | Epcam |
| ENSMUSG00000051748 | -6.8194366 | Wfdc21 |
| ENSMUSG00000031506 | -6.4771283 | Ptpn7 |
| ENSMUSG00000029304 | -6.4149182 | Spp1 |
| ENSMUSG00000064023 | -6.4067827 | Klk8 |
| ENSMUSG00000024424 | -6.2124133 | Ttc39c |
| ENSMUSG00000022512 | -6.1696262 | Cldn1 |
| ENSMUSG00000061825 | -6.1693856 | Ces2c |
| ENSMUSG00000002228 | -6.1295692 | Ppm1j |
| ENSMUSG00000079523 | -5.9369123 | Tmsb10 |
| ENSMUSG00000098489 | -5.9341154 | Mir7678 |
| ENSMUSG00000000805 | -5.9115921 | Car4 |
| ENSMUSG00000091955 | -5.7262758 | Gm9844 |
| ENSMUSG00000037868 | -5.6346782 | Egr2 |
| ENSMUSG00000071847 | -5.5955301 | Apcdd1 |
| ENSMUSG00000020836 | -5.46411 | Coro6 |
| ENSMUSG00000094392 | -5.2446459 | Gm3788 |
| ENSMUSG00000039013 | -5.2397313 | Siglec5 |
| ENSMUSG00000023243 | -5.1155471 | Kcnk5 |
| ENSMUSG00000042417 | -5.079883 | Ccno |
| ENSMUSG00000040663 | -4.9946984 | Clcf1 |
| ENSMUSG00000063600 | -4.9486609 | Egfem1 |
| ENSMUSG00000055865 | -4.8985002 | Fam19a3 |
| ENSMUSG00000001588 | -4.8729214 | Acap1 |
| ENSMUSG00000079298 | -4.8712875 | Klrb1b |
| ENSMUSG00000032131 | -4.8454719 | Abcg4 |
| ENSMUSG00000041324 | -4.8077974 | Inhba |
| ENSMUSG00000029378 | -4.7070558 | Mcub |
| ENSMUSG00000105954 | -4.6561915 | Gm42793 |

**Supplementary Table 2**

| | | |
|---|---|---|
| ENSMUSG00000009614 | -4.6097299 | Sardh |
| ENSMUSG00000002233 | -4.5849763 | Rhoc |
| ENSMUSG00000072601 | -4.5659602 | Ear1 |
| ENSMUSG00000005950 | -4.54166 | P2rx5 |
| ENSMUSG00000099032 | -4.4738977 | Tcf24 |
| ENSMUSG00000000915 | -4.3113298 | Hip1r |
| ENSMUSG00000031443 | -4.2471606 | F7 |
| ENSMUSG00000046961 | -4.187537 | Gpr156 |
| ENSMUSG00000029469 | -4.1725242 | Ift81 |
| ENSMUSG00000032192 | -4.1525952 | Gnb5 |
| ENSMUSG00000038764 | -4.1513595 | Ptpn3 |
| ENSMUSG00000005465 | -4.0216803 | Il27ra |
| ENSMUSG00000024768 | -3.9753065 | Lipf |
| ENSMUSG00000107429 | -3.9414118 | Gm44206 |
| ENSMUSG00000043017 | -3.934126 | Ptgir |
| ENSMUSG00000062372 | -3.9005478 | Otof |
| ENSMUSG00000044674 | -3.8875978 | Fzd1 |
| ENSMUSG00000027611 | -3.8370197 | Procr |
| ENSMUSG00000082629 | -3.798829 | Gm5639 |
| ENSMUSG00000048965 | -3.7755906 | Mrgpre |
| ENSMUSG00000005057 | -3.7681342 | Sh2b2 |
| ENSMUSG00000022102 | -3.7588193 | Dok2 |
| ENSMUSG00000072600 | -3.7522279 | Ear-ps9 |
| ENSMUSG00000046598 | -3.7326134 | Bdh1 |
| ENSMUSG00000055407 | -3.6813874 | Map6 |
| ENSMUSG00000035355 | -3.6602388 | Kcnh4 |
| ENSMUSG00000032911 | -3.6257766 | Cspg4 |
| ENSMUSG00000034435 | -3.5978899 | Tmem30b |
| ENSMUSG00000086862 | -3.5902862 | Gm13546 |
| ENSMUSG00000032500 | -3.5731041 | Dclk3 |
| ENSMUSG00000025402 | -3.544089 | Nab2 |
| ENSMUSG00000062148 | -3.5219445 | Ear6 |
| ENSMUSG00000035148 | -3.4994386 | Gpr33 |
| ENSMUSG00000049588 | -3.4742476 | Ccdc69 |
| ENSMUSG00000104068 | -3.4414497 | Gm37199 |
| ENSMUSG00000022661 | -3.3540249 | Cd200 |
| ENSMUSG00000030930 | -3.3262829 | Chst15 |
| ENSMUSG00000027030 | -3.3211124 | Stk39 |
| ENSMUSG00000048218 | -3.2998304 | Amigo2 |
| ENSMUSG00000079534 | -3.2715851 | Gm5640 |
| ENSMUSG00000049409 | -3.2566371 | Prokr1 |
| ENSMUSG00000095609 | -3.2326531 | Gm21188 |
| ENSMUSG00000036098 | -3.2298783 | Myrf |

**Supplementary Table 2**

| | | |
|---|---|---|
| ENSMUSG00000017417 | -3.1603855 | Plxdc1 |
| ENSMUSG00000028194 | -3.1578582 | Ddah1 |
| ENSMUSG00000006435 | -3.1532825 | Neurl1a |
| ENSMUSG00000003452 | -3.1443013 | Bicd1 |
| ENSMUSG00000027296 | -3.0996292 | Itpka |
| ENSMUSG00000026193 | -3.0963135 | Fn1 |
| ENSMUSG00000054200 | -3.0944743 | Ffar4 |
| ENSMUSG00000036390 | -3.065823 | Gadd45a |
| ENSMUSG00000032105 | -3.0568324 | Pdzd3 |
| ENSMUSG00000061013 | -3.054516 | Mkx |
| ENSMUSG00000022235 | -3.0490626 | Cmbl |
| ENSMUSG00000070529 | -3.0400993 | Wfdc10 |
| ENSMUSG00000079531 | -3.0374589 | Gm5936 |
| ENSMUSG00000049404 | -3.0278703 | Rarres1 |
| ENSMUSG00000041842 | -2.9960268 | Fhdc1 |
| ENSMUSG00000022949 | -2.9927789 | Clic6 |
| ENSMUSG00000051652 | -2.978601 | Lrrc3 |
| ENSMUSG00000037279 | -2.9783889 | Ovol2 |
| ENSMUSG00000108763 | -2.9727564 | Gm36028 |
| ENSMUSG00000027737 | -2.9721566 | Slc7a11 |
| ENSMUSG00000018570 | -2.9663535 | 2810408A11Rik |
| ENSMUSG00000039372 | -2.9278477 | Mar-04 |
| ENSMUSG00000038970 | -2.9259363 | Lmtk2 |
| ENSMUSG00000022421 | -2.8909905 | Nptxr |
| ENSMUSG00000030159 | -2.8774981 | Clec1b |
| ENSMUSG00000049526 | -2.8527377 | Tmem202 |
| ENSMUSG00000038473 | -2.8513398 | Nos1ap |
| ENSMUSG00000042439 | -2.8358346 | Zfp532 |
| ENSMUSG00000072964 | -2.8261791 | Bhlhb9 |
| ENSMUSG00000051177 | -2.8250334 | Plcb1 |
| ENSMUSG00000026220 | -2.8242563 | Slc16a14 |
| ENSMUSG00000085642 | -2.7828651 | 3110053B16Rik |
| ENSMUSG00000017400 | -2.7660194 | Stac2 |
| ENSMUSG00000044400 | -2.740567 | Sowahd |
| ENSMUSG00000024866 | -2.7299751 | Acy3 |
| ENSMUSG00000031355 | -2.7285045 | Arhgap6 |
| ENSMUSG00000026981 | -2.7271263 | Il1rn |
| ENSMUSG00000031373 | -2.7207156 | Car5b |
| ENSMUSG00000067771 | -2.7068238 | Gm14685 |
| ENSMUSG00000031078 | -2.7029191 | Cttn |
| ENSMUSG00000072572 | -2.6907509 | Slc39a2 |
| ENSMUSG00000034557 | -2.6474441 | Zfyve9 |
| ENSMUSG00000045725 | -2.6451752 | Prr15 |

**Supplementary Table 2**

| | | |
|---|---|---|
| ENSMUSG00000026893 | -2.6338836 | Gca |
| ENSMUSG00000024084 | -2.6282107 | Qpct |
| ENSMUSG00000036459 | -2.5860496 | Wtip |
| ENSMUSG00000041220 | -2.5677484 | Elovl6 |
| ENSMUSG00000060568 | -2.5574416 | Fam78b |
| ENSMUSG00000024008 | -2.5528773 | Cpne5 |
| ENSMUSG00000042115 | -2.5452767 | Klhdc8a |
| ENSMUSG00000074604 | -2.5402242 | Mgst2 |
| ENSMUSG00000069755 | -2.5309783 | Zfp125 |
| ENSMUSG00000034656 | -2.5283612 | Cacna1a |
| ENSMUSG00000029086 | -2.5271034 | Prom1 |
| ENSMUSG00000071745 | -2.5249291 | DXBay18 |
| ENSMUSG00000093598 | -2.5092539 | A730085K08Rik |
| ENSMUSG00000036103 | -2.4987732 | Colec12 |
| ENSMUSG00000097779 | -2.4965118 | 4833407H14Rik |
| ENSMUSG00000031444 | -2.4801399 | F10 |
| ENSMUSG00000017607 | -2.4793836 | Tns4 |
| ENSMUSG00000041757 | -2.4636188 | Plekha6 |
| ENSMUSG00000036402 | -2.4605451 | Gng12 |
| ENSMUSG00000102380 | -2.452079 | Gm38140 |
| ENSMUSG00000035900 | -2.4339019 | Gramd4 |
| ENSMUSG00000028943 | -2.4156002 | Espn |
| ENSMUSG00000068758 | -2.3909601 | Il3ra |
| ENSMUSG00000106609 | -2.3860832 | Gm43181 |
| ENSMUSG00000049420 | -2.3830945 | Tmem200a |
| ENSMUSG00000096950 | -2.3677924 | Gm9530 |
| ENSMUSG00000032786 | -2.360015 | Alas1 |
| ENSMUSG00000032902 | -2.3558687 | Slc16a1 |
| ENSMUSG00000087684 | -2.3507909 | 1200007C13Rik |
| ENSMUSG00000037148 | -2.3362288 | Arhgap10 |
| ENSMUSG00000108859 | -2.3326156 | Gm44776 |
| ENSMUSG00000030359 | -2.330848 | Pzp |
| ENSMUSG00000036273 | -2.3272117 | Lrrk2 |
| ENSMUSG00000020656 | -2.3167156 | Grhl1 |
| ENSMUSG00000061451 | -2.2958409 | Tmem151a |
| ENSMUSG00000026630 | -2.253943 | Batf3 |
| ENSMUSG00000019806 | -2.2474935 | Aig1 |
| ENSMUSG00000027313 | -2.2057464 | Chac1 |
| ENSMUSG00000032656 | -2.1974494 | Mar-03 |
| ENSMUSG00000040209 | -2.1901618 | Zfp704 |
| ENSMUSG00000026065 | -2.1855143 | Slc9a4 |
| ENSMUSG00000063193 | -2.1706351 | Cd300lb |
| ENSMUSG00000036661 | -2.1603813 | Dennd3 |

**Supplementary Table 2**

| | | |
|---|---|---|
| ENSMUSG00000024667 | -2.1517229 | Tmem216 |
| ENSMUSG00000026187 | -2.1479837 | Xrcc5 |
| ENSMUSG00000042688 | -2.1316903 | Mapk6 |
| ENSMUSG00000051832 | -2.1274829 | E230016K23Rik |
| ENSMUSG00000026765 | -2.1079142 | Lypd6b |
| ENSMUSG00000059956 | -2.0949592 | Serpinb12 |
| ENSMUSG00000045045 | -2.0659315 | Lrfn4 |
| ENSMUSG00000090626 | -2.0476988 | Tex9 |
| ENSMUSG00000059606 | -2.0464869 | Rnase2b |
| ENSMUSG00000037493 | -2.044698 | Cib2 |
| ENSMUSG00000041794 | -2.0394223 | Myrip |
| ENSMUSG00000001095 | -2.0328425 | Slc13a2 |
| ENSMUSG00000047293 | -2.0260745 | Gpr15 |
| ENSMUSG00000083307 | -2.0226246 | AA414768 |
| ENSMUSG00000002500 | -2.0214868 | Rpl3l |
| ENSMUSG00000026811 | -2.012799 | St6galnac6 |
| ENSMUSG00000013584 | -2.0007006 | Aldh1a2 |
| ENSMUSG00000062464 | -1.9975375 | Cyp4f37 |
| ENSMUSG00000074796 | -1.9885623 | Slc4a11 |
| ENSMUSG00000013523 | -1.9865443 | Bcas1 |
| ENSMUSG00000036560 | -1.9858972 | Lgi4 |
| ENSMUSG00000101848 | -1.9793907 | 4933417E11Rik |
| ENSMUSG00000062028 | -1.9771209 | Irgc1 |
| ENSMUSG00000000244 | -1.9519604 | Tspan32 |
| ENSMUSG00000037406 | -1.9400966 | Htra4 |
| ENSMUSG00000040899 | -1.923776 | Ccr6 |
| ENSMUSG00000057134 | -1.9114067 | Ado |
| ENSMUSG00000102752 | -1.8903301 | Gm7694 |
| ENSMUSG00000025804 | -1.8896343 | Ccr1 |
| ENSMUSG00000000253 | -1.88412 | Gmpr |
| ENSMUSG00000021057 | -1.8747873 | Akap5 |
| ENSMUSG00000034731 | -1.8645927 | Dgkh |
| ENSMUSG00000042268 | -1.8583282 | Slc26a9 |
| ENSMUSG00000038244 | -1.8553996 | Mical2 |
| ENSMUSG00000073409 | -1.8313669 | H2-Q6 |
| ENSMUSG00000094257 | -1.8283326 | Ap3s1-ps2 |
| ENSMUSG00000024074 | -1.8281759 | Crim1 |
| ENSMUSG00000073184 | -1.8158137 | Gm10479 |
| ENSMUSG00000041782 | -1.8036912 | Lad1 |
| ENSMUSG00000056671 | -1.783716 | Prelid2 |
| ENSMUSG00000076441 | -1.7626873 | Ass1 |
| ENSMUSG00000081534 | -1.7526681 | Slc48a1 |
| ENSMUSG00000090231 | -1.7519033 | Cfb |

**Supplementary Table 2**

| | | |
|---|---|---|
| ENSMUSG00000052516 | -1.7511019 | Robo2 |
| ENSMUSG00000075289 | -1.7484256 | Carns1 |
| ENSMUSG00000090394 | -1.7413069 | 4930523C07Rik |
| ENSMUSG00000027994 | -1.7245233 | Mcub |
| ENSMUSG00000032014 | -1.7216565 | Oaf |
| ENSMUSG00000026782 | -1.7196483 | Abi2 |
| ENSMUSG00000048216 | -1.693036 | Gpr85 |
| ENSMUSG00000070305 | -1.6883855 | Mpzl3 |
| ENSMUSG00000079009 | -1.6821978 | Gm14139 |
| ENSMUSG00000028937 | -1.6765012 | Acot7 |
| ENSMUSG00000017756 | -1.673554 | Slc12a7 |
| ENSMUSG00000089665 | -1.6722343 | Fcor |
| ENSMUSG00000079293 | -1.6655913 | Clec7a |
| ENSMUSG00000042289 | -1.6592885 | Hsd3b7 |
| ENSMUSG00000092349 | -1.6550381 | Platr17 |
| ENSMUSG00000037600 | -1.6445672 | Kdf1 |
| ENSMUSG00000026102 | -1.6386192 | Inpp1 |
| ENSMUSG00000036545 | -1.6334237 | Adamts2 |
| ENSMUSG00000091618 | -1.6313931 | H60c |
| ENSMUSG00000109378 | -1.6205394 | U2af1l4 |
| ENSMUSG00000052270 | -1.6146513 | Fpr2 |
| ENSMUSG00000032589 | -1.6072584 | Bsn |
| ENSMUSG00000017639 | -1.6071964 | Rab11fip4 |
| ENSMUSG00000036587 | -1.6058546 | Fut7 |
| ENSMUSG00000015312 | -1.6020869 | Gadd45b |
| ENSMUSG00000051397 | -1.6008258 | Tacstd2 |
| ENSMUSG00000078238 | -1.5956819 | Gm12854 |
| ENSMUSG00000081984 | -1.5842994 | Dnajb3 |
| ENSMUSG00000041954 | -1.5836515 | Tnfrsf18 |
| ENSMUSG00000049265 | -1.5745682 | Kcnk3 |
| ENSMUSG00000070462 | -1.5733034 | Tlnrd1 |
| ENSMUSG00000073179 | -1.5674749 | Gm10478 |
| ENSMUSG00000073197 | -1.5617492 | 5730507C01Rik |
| ENSMUSG00000030546 | -1.5519149 | Plin1 |
| ENSMUSG00000108713 | -1.5401201 | Gm33027 |
| ENSMUSG00000062300 | -1.5393436 | Nectin2 |
| ENSMUSG00000039530 | -1.5351316 | Tusc3 |
| ENSMUSG00000024302 | -1.534012 | Dtna |
| ENSMUSG00000045573 | -1.5307424 | Penk |
| ENSMUSG00000049327 | -1.516072 | Kmt5a |
| ENSMUSG00000039063 | -1.5124405 | Echdc3 |
| ENSMUSG00000023019 | -1.5119324 | Gpd1 |
| ENSMUSG00000053550 | -1.5114483 | Shisa7 |

### Supplementary Table 2

| | | |
|---|---|---|
| ENSMUSG00000033287 | -1.5096971 | Kctd17 |
| ENSMUSG00000108402 | -1.5068078 | 9430064I24Rik |
| ENSMUSG00000054422 | -1.5016594 | Fabp1 |
| ENSMUSG00000078735 | -1.5003475 | Il11ra2 |
| ENSMUSG00000020205 | -1.499965 | Phlda1 |
| ENSMUSG00000072663 | -1.4980738 | Spef2 |
| ENSMUSG00000026679 | -1.4965409 | Enkur |
| ENSMUSG00000022844 | -1.4948798 | Pdia5 |
| ENSMUSG00000026447 | -1.4938578 | Pik3c2b |
| ENSMUSG00000022439 | -1.4935403 | Parvg |
| ENSMUSG00000082088 | -1.4925089 | Gm15753 |
| ENSMUSG00000034758 | -1.4797008 | Tle6 |
| ENSMUSG00000058022 | -1.4786127 | Adtrp |
| ENSMUSG00000070291 | -1.474012 | Ddx43 |
| ENSMUSG00000086644 | -1.4735092 | Gm13470 |
| ENSMUSG00000008393 | -1.4681384 | Carhsp1 |
| ENSMUSG00000021453 | -1.462513 | Gadd45g |
| ENSMUSG00000015501 | -1.4606294 | Hivep2 |
| ENSMUSG00000043079 | -1.4524186 | Synpo |
| ENSMUSG00000029401 | -1.4482791 | Rilpl2 |
| ENSMUSG00000034993 | -1.4403915 | Vat1 |
| ENSMUSG00000014599 | -1.4362423 | Csf1 |
| ENSMUSG00000063628 | -1.4299348 | Gm7665 |
| ENSMUSG00000025324 | -1.4297942 | Atp10a |
| ENSMUSG00000037418 | -1.4294087 | Best1 |
| ENSMUSG00000021477 | -1.4261058 | Ctsl |
| ENSMUSG00000109420 | -1.4256807 | Gm44702 |
| ENSMUSG00000027580 | -1.424992 | Helz2 |
| ENSMUSG00000038372 | -1.4216779 | Gmds |
| ENSMUSG00000021097 | -1.4200339 | Clmn |
| ENSMUSG00000104484 | -1.416766 | Gm33142 |
| ENSMUSG00000032487 | -1.412451 | Ptgs2 |
| ENSMUSG00000037966 | -1.4102915 | Ninj1 |
| ENSMUSG00000027907 | -1.4047591 | S100a11 |
| ENSMUSG00000042106 | -1.4015145 | Fam212a |
| ENSMUSG00000041380 | -1.3992122 | Htr2c |
| ENSMUSG00000033467 | -1.3959803 | Crlf2 |
| ENSMUSG00000026390 | -1.3882987 | Marco |
| ENSMUSG00000033192 | -1.3876459 | Lpcat2 |
| ENSMUSG00000067321 | -1.384991 | Gm7931 |
| ENSMUSG00000037905 | -1.384611 | Bri3bp |
| ENSMUSG00000015476 | -1.3821298 | Prrt1 |
| ENSMUSG00000074682 | -1.3777548 | Zcchc3 |

**Supplementary Table 2**

| | | |
|---|---|---|
| ENSMUSG00000081467 | -1.3751014 | Gm13337 |
| ENSMUSG00000026819 | -1.3724824 | Slc25a25 |
| ENSMUSG00000034765 | -1.3672496 | Dusp5 |
| ENSMUSG00000052727 | -1.3632107 | Map1b |
| ENSMUSG00000060550 | -1.3624988 | H2-Q7 |
| ENSMUSG00000040084 | -1.3581758 | Bub1b |
| ENSMUSG00000066735 | -1.3535423 | Vkorc1l1 |
| ENSMUSG00000062312 | -1.3534095 | Erbb2 |
| ENSMUSG00000021262 | -1.337444 | Evl |
| ENSMUSG00000026822 | -1.3372316 | Lcn2 |
| ENSMUSG00000057093 | -1.3360597 | Zfp607b |
| ENSMUSG00000034271 | -1.3309803 | Jdp2 |
| ENSMUSG00000052698 | -1.3248772 | Tln2 |
| ENSMUSG00000018559 | -1.3244097 | Ctdnep1 |
| ENSMUSG00000074151 | -1.319192 | Nlrc5 |
| ENSMUSG00000003038 | -1.3171708 | Hmgn2 |
| ENSMUSG00000041695 | -1.3141677 | Kcnj2 |
| ENSMUSG00000029379 | -1.305664 | Cxcl3 |
| ENSMUSG00000041754 | -1.303941 | Trem3 |
| ENSMUSG00000025466 | -1.3038833 | Fuom |
| ENSMUSG00000002058 | -1.29812 | Unc119 |
| ENSMUSG00000031497 | -1.2917087 | Tnfsf13b |
| ENSMUSG00000039431 | -1.285694 | Mtmr7 |
| ENSMUSG00000094306 | -1.282474 | Gm24924 |
| ENSMUSG00000018927 | -1.2760249 | Ccl6 |
| ENSMUSG00000010660 | -1.2753083 | Plcd1 |
| ENSMUSG00000043252 | -1.2743764 | Tmem64 |
| ENSMUSG00000026207 | -1.2627638 | Speg |
| ENSMUSG00000030126 | -1.2558273 | Tmcc1 |
| ENSMUSG00000079845 | -1.2544932 | Xlr4a |
| ENSMUSG00000032776 | -1.2529979 | Mctp2 |
| ENSMUSG00000028645 | -1.2401112 | Slc2a1 |
| ENSMUSG00000081650 | -1.2348217 | Gm16181 |
| ENSMUSG00000086564 | -1.2327084 | Cd101 |
| ENSMUSG00000105771 | -1.2255486 | 2900064K03Rik |
| ENSMUSG00000070713 | -1.2232869 | Gm10282 |
| ENSMUSG00000036949 | -1.2211654 | Slc39a12 |
| ENSMUSG00000036634 | -1.2210014 | Mag |
| ENSMUSG00000064899 | -1.2165687 | Snord118 |
| ENSMUSG00000032263 | -1.2097788 | Bckdhb |
| ENSMUSG00000022221 | -1.2026882 | Ripk3 |
| ENSMUSG00000045414 | -1.2026852 | 1190002N15Rik |
| ENSMUSG00000030232 | -1.2015154 | Aebp2 |

**Supplementary Table 2**

| | | |
|---|---|---|
| ENSMUSG00000079067 | -1.1988451 | Hmgn2-ps1 |
| ENSMUSG00000033147 | -1.1941126 | Slc22a15 |
| ENSMUSG00000020808 | -1.1910926 | Pimreg |
| ENSMUSG00000039234 | -1.1876798 | Sec24d |
| ENSMUSG00000044149 | -1.1859245 | Nkrf |
| ENSMUSG00000021263 | -1.185874 | Degs2 |
| ENSMUSG00000037172 | -1.1785753 | E330009J07Rik |
| ENSMUSG00000028238 | -1.1784266 | Atp6v0d2 |
| ENSMUSG00000034006 | -1.1744484 | Pqlc1 |
| ENSMUSG00000024164 | -1.1741564 | C3 |
| ENSMUSG00000046794 | -1.1722929 | Ppp1r3b |
| ENSMUSG00000073889 | -1.169369 | Il11ra1 |
| ENSMUSG00000026991 | -1.1663952 | Pkp4 |
| ENSMUSG00000024640 | -1.1642921 | Psat1 |
| ENSMUSG00000006651 | -1.1640117 | Aplp1 |
| ENSMUSG00000041762 | -1.1626993 | Gpr155 |
| ENSMUSG00000069516 | -1.1625128 | Lyz2 |
| ENSMUSG00000004880 | -1.1593486 | Lbr |
| ENSMUSG00000042700 | -1.1502916 | Sipa1l1 |
| ENSMUSG00000075010 | -1.1481555 | AW112010 |
| ENSMUSG00000053469 | -1.1477244 | Tg |
| ENSMUSG00000073876 | -1.1360265 | Gm13305 |
| ENSMUSG00000055413 | -1.1353187 | H2-Q5 |
| ENSMUSG00000000184 | -1.1348703 | Ccnd2 |
| ENSMUSG00000091498 | -1.1313057 | Mpc1-ps |
| ENSMUSG00000110495 | -1.122607 | Gm45879 |
| ENSMUSG00000030187 | -1.1210962 | Klra2 |
| ENSMUSG00000103308 | -1.1191525 | Gm37800 |
| ENSMUSG00000029761 | -1.116638 | Cald1 |
| ENSMUSG00000018581 | -1.1138649 | Dnah11 |
| ENSMUSG00000034445 | -1.1107924 | Cyb561a3 |
| ENSMUSG00000023861 | -1.1090433 | Mpc1 |
| ENSMUSG00000038648 | -1.1007121 | Creb3l2 |
| ENSMUSG00000030659 | -1.1003546 | Nucb2 |
| ENSMUSG00000029127 | -1.0956853 | Zbtb49 |
| ENSMUSG00000017390 | -1.0847335 | Aldoc |
| ENSMUSG00000085152 | -1.0714608 | Gm11496 |
| ENSMUSG00000010663 | -1.0713069 | Fads1 |
| ENSMUSG00000034258 | -1.0703731 | Flvcr2 |
| ENSMUSG00000025178 | -1.0642563 | Pi4k2a |
| ENSMUSG00000110588 | -1.0635594 | Gm45774 |
| ENSMUSG00000027002 | -1.0628556 | Nckap1 |
| ENSMUSG00000034591 | -1.0626525 | Slc41a2 |

**Supplementary Table 2**

| | | |
|---|---|---|
| ENSMUSG00000073678 | -1.0607527 | Pgap1 |
| ENSMUSG00000068036 | -1.0590008 | Afdn |
| ENSMUSG00000085611 | -1.0576737 | Ap3s1-ps1 |
| ENSMUSG00000074738 | -1.0571048 | Fndc10 |
| ENSMUSG00000045312 | -1.0564587 | Lhfpl2 |
| ENSMUSG00000020911 | -1.0550267 | Krt19 |
| ENSMUSG00000022957 | -1.0479279 | Itsn1 |
| ENSMUSG00000110498 | -1.0477651 | A630001O12Rik |
| ENSMUSG00000050921 | -1.0468545 | P2ry10 |
| ENSMUSG00000106019 | -1.046751 | Gm43672 |
| ENSMUSG00000029925 | -1.0458493 | Tbxas1 |
| ENSMUSG00000032013 | -1.0408442 | Trim29 |
| ENSMUSG00000030206 | -1.0376512 | Gsg1 |
| ENSMUSG00000029098 | -1.0365593 | Acox3 |
| ENSMUSG00000056692 | -1.0353955 | D17Wsu92e |
| ENSMUSG00000037095 | -1.0352062 | Lrg1 |
| ENSMUSG00000059401 | -1.0348516 | Mamld1 |
| ENSMUSG00000044734 | -1.0341532 | Serpinb1a |
| ENSMUSG00000030172 | -1.0325283 | Erc1 |
| ENSMUSG00000033721 | -1.029456 | Vav3 |
| ENSMUSG00000034201 | -1.0255216 | Gas2l1 |
| ENSMUSG00000006378 | -1.0242807 | Gcat |
| ENSMUSG00000086606 | -1.0239586 | Gm13205 |
| ENSMUSG00000030160 | -1.0232815 | Tmem52b |
| ENSMUSG00000044199 | -1.0199856 | S1pr4 |
| ENSMUSG00000005045 | -1.0172628 | Chd5 |
| ENSMUSG00000044244 | -1.0157295 | Il20rb |
| ENSMUSG00000029373 | -1.015576 | Pf4 |
| ENSMUSG00000049577 | -1.0118696 | Zfpm1 |
| ENSMUSG00000106104 | -1.010261 | Gm42660 |
| ENSMUSG00000029814 | -1.007878 | Igf2bp3 |
| ENSMUSG00000056753 | -1.0065153 | C330011M18Rik |
| ENSMUSG00000040146 | -1.0015266 | Rgl3 |

### Supplementary Table 2

### Upregulated in Cre+ (*Egr2* deficient)

| | | |
|---|---|---|
| ENSMUSG00000031155 | 1.00030356 | Pim2 |
| ENSMUSG00000025006 | 1.00392092 | Sorbs1 |
| ENSMUSG00000049687 | 1.00401398 | Fam109b |
| ENSMUSG00000067199 | 1.00493534 | Frat1 |
| ENSMUSG00000032366 | 1.00799601 | Tpm1 |
| ENSMUSG00000031431 | 1.00932069 | Tsc22d3 |
| ENSMUSG00000033705 | 1.01074598 | Stard9 |
| ENSMUSG00000033213 | 1.01105267 | AA467197 |
| ENSMUSG00000027478 | 1.01354667 | Dnmt3b |
| ENSMUSG00000030231 | 1.01395033 | Plekha5 |
| ENSMUSG00000049744 | 1.01670362 | Arhgap15 |
| ENSMUSG00000030393 | 1.0180171 | Zik1 |
| ENSMUSG00000063172 | 1.01982521 | Hspb11 |
| ENSMUSG00000042599 | 1.02399403 | Kdm7a |
| ENSMUSG00000031951 | 1.02624163 | Tmem231 |
| ENSMUSG00000041642 | 1.03417669 | Kif21b |
| ENSMUSG00000049988 | 1.03748544 | Lrrc25 |
| ENSMUSG00000085457 | 1.04034431 | 1110046J04Rik |
| ENSMUSG00000021948 | 1.04450192 | Prkcd |
| ENSMUSG00000024222 | 1.04469412 | Fkbp5 |
| ENSMUSG00000032375 | 1.0449475 | Aph1b |
| ENSMUSG00000010651 | 1.04624865 | Acaa1b |
| ENSMUSG00000028933 | 1.04755303 | Xrcc2 |
| ENSMUSG00000021007 | 1.05139661 | Spata7 |
| ENSMUSG00000062098 | 1.05232516 | Btbd3 |
| ENSMUSG00000038578 | 1.05352419 | Susd1 |
| ENSMUSG00000074221 | 1.05443076 | Zfp568 |
| ENSMUSG00000030960 | 1.05527977 | Eef1akmt2 |
| ENSMUSG00000025044 | 1.0554828 | Msr1 |
| ENSMUSG00000045658 | 1.05770051 | Pid1 |
| ENSMUSG00000076431 | 1.06119527 | Sox4 |
| ENSMUSG00000040219 | 1.06772282 | Ttc12 |
| ENSMUSG00000053411 | 1.06954078 | Cbx7 |
| ENSMUSG00000030589 | 1.07538318 | Rasgrp4 |
| ENSMUSG00000029650 | 1.07714798 | Slc46a3 |
| ENSMUSG00000040339 | 1.07807705 | Fam102b |
| ENSMUSG00000049881 | 1.07869878 | 2810025M15Rik |
| ENSMUSG00000020422 | 1.08175058 | Tns3 |
| ENSMUSG00000030729 | 1.08807372 | Pgm2l1 |
| ENSMUSG00000068551 | 1.08989868 | Zfp467 |
| ENSMUSG00000030760 | 1.09299943 | Acer3 |

**Supplementary Table 2**

| | | |
|---|---|---|
| ENSMUSG00000086247 | 1.09310341 | Gm15787 |
| ENSMUSG00000038740 | 1.09504457 | Mvb12b |
| ENSMUSG00000017009 | 1.09797994 | Sdc4 |
| ENSMUSG00000050270 | 1.0981327 | Tmem220 |
| ENSMUSG00000043664 | 1.0984167 | Tmem221 |
| ENSMUSG00000024672 | 1.0990219 | Ms4a7 |
| ENSMUSG00000035776 | 1.09923538 | Cd99l2 |
| ENSMUSG00000060675 | 1.10401749 | Pla2g16 |
| ENSMUSG00000053063 | 1.11283065 | Clec12a |
| ENSMUSG00000035208 | 1.11486003 | Slfn8 |
| ENSMUSG00000100975 | 1.11797896 | Gm28875 |
| ENSMUSG00000024892 | 1.12225389 | Pcx |
| ENSMUSG00000070565 | 1.12514426 | Rasal2 |
| ENSMUSG00000015850 | 1.12926201 | Adamtsl4 |
| ENSMUSG00000086401 | 1.13297041 | Gm15559 |
| ENSMUSG00000002204 | 1.13495492 | Napsa |
| ENSMUSG00000022885 | 1.13644354 | St6gal1 |
| ENSMUSG00000030592 | 1.13671506 | Ryr1 |
| ENSMUSG00000007216 | 1.13771521 | Zfp775 |
| ENSMUSG00000020829 | 1.14178751 | Slc46a1 |
| ENSMUSG00000047182 | 1.14736898 | Irs3 |
| ENSMUSG00000035954 | 1.15015234 | Dock4 |
| ENSMUSG00000032507 | 1.15675575 | Fbxl2 |
| ENSMUSG00000021822 | 1.16338139 | Plau |
| ENSMUSG00000054150 | 1.16647197 | Syne3 |
| ENSMUSG00000017754 | 1.17094446 | Pltp |
| ENSMUSG00000019763 | 1.17371256 | Rmnd1 |
| ENSMUSG00000000686 | 1.17432493 | Abhd15 |
| ENSMUSG00000000673 | 1.178414 | Haao |
| ENSMUSG00000074194 | 1.1859914 | Zfp791 |
| ENSMUSG00000056724 | 1.18846174 | Nbeal2 |
| ENSMUSG00000027009 | 1.18855458 | Itga4 |
| ENSMUSG00000056529 | 1.18855974 | Ptafr |
| ENSMUSG00000050212 | 1.19565162 | Eva1b |
| ENSMUSG00000071537 | 1.19728562 | Klrg2 |
| ENSMUSG00000017309 | 1.19917966 | Cd300lg |
| ENSMUSG00000016526 | 1.20011847 | Dyrk3 |
| ENSMUSG00000022040 | 1.20092425 | Ephx2 |
| ENSMUSG00000021572 | 1.20201276 | Cep72 |
| ENSMUSG00000004791 | 1.20322899 | Pgf |
| ENSMUSG00000079038 | 1.20367671 | D130040H23Rik |
| ENSMUSG00000040270 | 1.20391484 | Bach2 |
| ENSMUSG00000030946 | 1.21190447 | Lhpp |

**Supplementary Table 2**

| | | |
|---|---|---|
| ENSMUSG00000087366 | 1.21473214 | Junos |
| ENSMUSG00000032845 | 1.21480395 | Alpk2 |
| ENSMUSG00000020546 | 1.21882428 | Stxbp4 |
| ENSMUSG00000058163 | 1.21922051 | Gm5431 |
| ENSMUSG00000052684 | 1.23041393 | Jun |
| ENSMUSG00000037185 | 1.23043806 | Krt80 |
| ENSMUSG00000005125 | 1.23086401 | Ndrg1 |
| ENSMUSG00000099767 | 1.23133537 | Gm28884 |
| ENSMUSG00000006362 | 1.23274167 | Cbfa2t3 |
| ENSMUSG00000031925 | 1.23284655 | Maml2 |
| ENSMUSG00000042035 | 1.23312778 | Igsf3 |
| ENSMUSG00000052821 | 1.23327284 | Cysltr1 |
| ENSMUSG00000056054 | 1.23425683 | S100a8 |
| ENSMUSG00000028035 | 1.23487729 | Dnajb4 |
| ENSMUSG00000046727 | 1.23677013 | Cystm1 |
| ENSMUSG00000108732 | 1.23956941 | 2310043P16Rik |
| ENSMUSG00000050530 | 1.24345233 | Fam171a1 |
| ENSMUSG00000049321 | 1.24976588 | Zfp2 |
| ENSMUSG00000018381 | 1.25110079 | Abi3 |
| ENSMUSG00000042515 | 1.25260922 | Mum1l1 |
| ENSMUSG00000033065 | 1.25402509 | Pfkm |
| ENSMUSG00000024049 | 1.25667831 | Myom1 |
| ENSMUSG00000022265 | 1.25763089 | Ank |
| ENSMUSG00000090799 | 1.26011446 | Klhl33 |
| ENSMUSG00000028859 | 1.26070366 | Csf3r |
| ENSMUSG00000072949 | 1.26121318 | Acot1 |
| ENSMUSG00000030465 | 1.26534568 | Psd3 |
| ENSMUSG00000046329 | 1.26882859 | Slc25a23 |
| ENSMUSG00000097296 | 1.27366275 | Gm26532 |
| ENSMUSG00000038456 | 1.27417552 | Dennd2a |
| ENSMUSG00000046169 | 1.27723141 | Adamts6 |
| ENSMUSG00000029298 | 1.27954257 | Gbp9 |
| ENSMUSG00000018459 | 1.28159824 | Slc13a3 |
| ENSMUSG00000026656 | 1.2818039 | Fcgr2b |
| ENSMUSG00000067714 | 1.28187811 | Lpar5 |
| ENSMUSG00000062794 | 1.28225507 | Zfp599 |
| ENSMUSG00000016382 | 1.2831801 | Pls3 |
| ENSMUSG00000020589 | 1.28471928 | Fam49a |
| ENSMUSG00000038248 | 1.28506392 | Sobp |
| ENSMUSG00000033209 | 1.28553552 | Ttc28 |
| ENSMUSG00000090115 | 1.28812022 | Usp49 |
| ENSMUSG00000053477 | 1.29190909 | Tcf4 |
| ENSMUSG00000074497 | 1.29294816 | A430078G23Rik |

**Supplementary Table 2**

| | | |
|---|---|---|
| ENSMUSG00000062373 | 1.29458461 | Tmem65 |
| ENSMUSG00000032739 | 1.30191548 | Pram1 |
| ENSMUSG00000007034 | 1.3040631 | Slc44a4 |
| ENSMUSG00000010142 | 1.30476128 | Tnfrsf13b |
| ENSMUSG00000030156 | 1.30614597 | Cd69 |
| ENSMUSG00000035299 | 1.32455248 | Mid1 |
| ENSMUSG00000032690 | 1.32648207 | Oas2 |
| ENSMUSG00000097284 | 1.32912173 | 4930480K23Rik |
| ENSMUSG00000035919 | 1.33513072 | Bbs9 |
| ENSMUSG00000043822 | 1.33531592 | Adamtsl5 |
| ENSMUSG00000018169 | 1.33604467 | Mfng |
| ENSMUSG00000032827 | 1.33787541 | Ppp1r9a |
| ENSMUSG00000047963 | 1.34375427 | Stbd1 |
| ENSMUSG00000021127 | 1.34465307 | Zfp36l1 |
| ENSMUSG00000085962 | 1.34770735 | Gm16984 |
| ENSMUSG00000029161 | 1.34951781 | Cgref1 |
| ENSMUSG00000044350 | 1.34999824 | Lacc1 |
| ENSMUSG00000044026 | 1.35580021 | Slc35g1 |
| ENSMUSG00000066842 | 1.35587252 | Hmcn1 |
| ENSMUSG00000041961 | 1.35601902 | Znrf3 |
| ENSMUSG00000093677 | 1.35754284 | Gm20712 |
| ENSMUSG00000027087 | 1.35841065 | Itgav |
| ENSMUSG00000033446 | 1.36197046 | Lpar6 |
| ENSMUSG00000036006 | 1.36271944 | Ripor2 |
| ENSMUSG00000029381 | 1.365717 | Shroom3 |
| ENSMUSG00000028047 | 1.36589803 | Thbs3 |
| ENSMUSG00000026271 | 1.36960805 | Gpr35 |
| ENSMUSG00000044807 | 1.37196817 | Zfp354c |
| ENSMUSG00000050967 | 1.37955602 | Creg2 |
| ENSMUSG00000036606 | 1.38427784 | Plxnb2 |
| ENSMUSG00000070044 | 1.38630599 | Fam149a |
| ENSMUSG00000032258 | 1.38868176 | Lca5 |
| ENSMUSG00000038782 | 1.39184351 | 1700028J19Rik |
| ENSMUSG00000085881 | 1.39396162 | Gm15912 |
| ENSMUSG00000111147 | 1.40314873 | AC160562.1 |
| ENSMUSG00000046006 | 1.4058791 | Gapt |
| ENSMUSG00000033450 | 1.41087086 | Tagap |
| ENSMUSG00000014496 | 1.41213935 | Ankrd28 |
| ENSMUSG00000020264 | 1.41251872 | Slc36a2 |
| ENSMUSG00000007872 | 1.41613305 | Id3 |
| ENSMUSG00000024654 | 1.41629553 | Asrgl1 |
| ENSMUSG00000030208 | 1.41783109 | Emp1 |
| ENSMUSG00000020577 | 1.42100628 | Tspan13 |

**Supplementary Table 2**

| | | |
|---|---|---|
| ENSMUSG00000005583 | 1.42213634 | Mef2c |
| ENSMUSG00000040722 | 1.42813898 | Scamp5 |
| ENSMUSG00000000303 | 1.4320551 | Cdh1 |
| ENSMUSG00000090877 | 1.43444073 | Hspa1b |
| ENSMUSG00000044468 | 1.43759828 | Fam46c |
| ENSMUSG00000014786 | 1.44530292 | Slc9a5 |
| ENSMUSG00000051043 | 1.44825914 | Gprc5c |
| ENSMUSG00000040410 | 1.45874467 | Fbxl4 |
| ENSMUSG00000046031 | 1.45878943 | Fam26f |
| ENSMUSG00000051339 | 1.45982401 | 2900026A02Rik |
| ENSMUSG00000015745 | 1.46748021 | Plekho1 |
| ENSMUSG00000055148 | 1.47237789 | Klf2 |
| ENSMUSG00000057191 | 1.47954937 | AB124611 |
| ENSMUSG00000040710 | 1.48062829 | St8sia4 |
| ENSMUSG00000031779 | 1.4834626 | Ccl22 |
| ENSMUSG00000048142 | 1.48387946 | Nat8l |
| ENSMUSG00000078763 | 1.48474839 | Slfn1 |
| ENSMUSG00000046323 | 1.48650198 | Dppa3 |
| ENSMUSG00000021573 | 1.4884087 | Tppp |
| ENSMUSG00000041658 | 1.49075562 | Rragb |
| ENSMUSG00000035547 | 1.49756762 | Capn5 |
| ENSMUSG00000031788 | 1.4990949 | Kifc3 |
| ENSMUSG00000025154 | 1.49913957 | Arhgap19 |
| ENSMUSG00000058290 | 1.51320332 | Espl1 |
| ENSMUSG00000068735 | 1.51495291 | Trp53i11 |
| ENSMUSG00000038074 | 1.51767787 | Fkbp14 |
| ENSMUSG00000020364 | 1.51783042 | Zfp354a |
| ENSMUSG00000081809 | 1.52050698 | Gm15539 |
| ENSMUSG00000092528 | 1.52389207 | Nlrp1c-ps |
| ENSMUSG00000093661 | 1.52865439 | Eif4e3 |
| ENSMUSG00000026815 | 1.53080461 | Gfi1b |
| ENSMUSG00000031778 | 1.53225321 | Cx3cl1 |
| ENSMUSG00000030523 | 1.53228804 | Trpm1 |
| ENSMUSG00000039997 | 1.55036038 | Ifi203 |
| ENSMUSG00000019820 | 1.55224958 | Utrn |
| ENSMUSG00000056832 | 1.55780812 | Ttc26 |
| ENSMUSG00000105687 | 1.55993732 | Gm6157 |
| ENSMUSG00000021640 | 1.56496873 | Naip1 |
| ENSMUSG00000091971 | 1.5672114 | Hspa1a |
| ENSMUSG00000026170 | 1.56893981 | Cyp27a1 |
| ENSMUSG00000034041 | 1.57653212 | Lyl1 |
| ENSMUSG00000040640 | 1.57789696 | Erc2 |
| ENSMUSG00000032420 | 1.57805412 | Nt5e |

### Supplementary Table 2

| | | |
|---|---|---|
| ENSMUSG00000107761 | 1.5982876 | 2010008C14Rik |
| ENSMUSG00000022756 | 1.60718707 | Slc7a4 |
| ENSMUSG00000085129 | 1.60760273 | 5031425F14Rik |
| ENSMUSG00000067206 | 1.61156996 | Lrrc66 |
| ENSMUSG00000019256 | 1.61302866 | Ahr |
| ENSMUSG00000021451 | 1.62260083 | Sema4d |
| ENSMUSG00000021986 | 1.62307459 | Amer2 |
| ENSMUSG00000060224 | 1.62626935 | Pyroxd2 |
| ENSMUSG00000006356 | 1.63256634 | Crip2 |
| ENSMUSG00000028445 | 1.63269336 | Enho |
| ENSMUSG00000073418 | 1.63363764 | C4b |
| ENSMUSG00000086150 | 1.63368831 | Bach2os |
| ENSMUSG00000074272 | 1.63380088 | Ceacam1 |
| ENSMUSG00000029919 | 1.63452681 | Hpgds |
| ENSMUSG00000039976 | 1.63929166 | Tbc1d16 |
| ENSMUSG00000073758 | 1.6440366 | Sh3d21 |
| ENSMUSG00000031565 | 1.64549413 | Fgfr1 |
| ENSMUSG00000030223 | 1.65495485 | Ptpro |
| ENSMUSG00000040703 | 1.66264846 | Cyp2s1 |
| ENSMUSG00000072621 | 1.67580806 | Slfn10-ps |
| ENSMUSG00000024140 | 1.67808044 | Epas1 |
| ENSMUSG00000037211 | 1.68417489 | Spry1 |
| ENSMUSG00000042810 | 1.68848823 | Krba1 |
| ENSMUSG00000022148 | 1.71183844 | Fyb |
| ENSMUSG00000026389 | 1.71617341 | Steap3 |
| ENSMUSG00000021356 | 1.71866217 | Irf4 |
| ENSMUSG00000042215 | 1.72151606 | Bag2 |
| ENSMUSG00000016283 | 1.72811247 | H2-M2 |
| ENSMUSG00000073274 | 1.7292352 | Gm14636 |
| ENSMUSG00000034917 | 1.72985663 | Tjp3 |
| ENSMUSG00000042766 | 1.73190196 | Trim46 |
| ENSMUSG00000057948 | 1.73539792 | Unc13d |
| ENSMUSG00000097313 | 1.7389395 | Gm26569 |
| ENSMUSG00000021322 | 1.73964816 | Aoah |
| ENSMUSG00000003206 | 1.74071962 | Ebi3 |
| ENSMUSG00000032348 | 1.74774944 | Gsta4 |
| ENSMUSG00000074622 | 1.74834296 | Mafb |
| ENSMUSG00000060512 | 1.74903873 | 0610040J01Rik |
| ENSMUSG00000053101 | 1.75027616 | Gpr141 |
| ENSMUSG00000021125 | 1.75429543 | Arg2 |
| ENSMUSG00000087242 | 1.7585295 | C78197 |
| ENSMUSG00000020053 | 1.76043534 | Igf1 |
| ENSMUSG00000031398 | 1.7628505 | Plxna3 |

**Supplementary Table 2**

| | | |
|---|---|---|
| ENSMUSG00000045625 | 1.76595218 | Pigz |
| ENSMUSG00000052336 | 1.7765219 | Cx3cr1 |
| ENSMUSG00000022901 | 1.78241816 | Cd86 |
| ENSMUSG00000025150 | 1.79269589 | Cbr2 |
| ENSMUSG00000079481 | 1.79621107 | Nhsl2 |
| ENSMUSG00000027843 | 1.81241425 | Ptpn22 |
| ENSMUSG00000028456 | 1.81494915 | Unc13b |
| ENSMUSG00000032278 | 1.82098663 | Paqr5 |
| ENSMUSG00000017466 | 1.82194363 | Timp2 |
| ENSMUSG00000032883 | 1.82831768 | Acsl3 |
| ENSMUSG00000018920 | 1.83144495 | Cxcl16 |
| ENSMUSG00000074093 | 1.83257619 | Svip |
| ENSMUSG00000024053 | 1.8379493 | Emilin2 |
| ENSMUSG00000021416 | 1.83887209 | Eci3 |
| ENSMUSG00000049191 | 1.87325869 | Rtl5 |
| ENSMUSG00000041058 | 1.87632103 | Wwp1 |
| ENSMUSG00000050994 | 1.87882963 | Adgb |
| ENSMUSG00000039485 | 1.87968495 | Tspyl4 |
| ENSMUSG00000004709 | 1.8835297 | Cd244 |
| ENSMUSG00000025372 | 1.89081225 | Baiap2 |
| ENSMUSG00000024913 | 1.97042317 | Lrp5 |
| ENSMUSG00000016552 | 1.98057752 | Foxred2 |
| ENSMUSG00000035678 | 1.98081305 | Tnfsf9 |
| ENSMUSG00000026589 | 1.99000407 | Sec16b |
| ENSMUSG00000024501 | 1.99036261 | Dpysl3 |
| ENSMUSG00000003153 | 1.9991831 | Slc2a3 |
| ENSMUSG00000030257 | 2.00109827 | Srgap3 |
| ENSMUSG00000049580 | 2.00749901 | Tsku |
| ENSMUSG00000014158 | 2.01199751 | Trpv4 |
| ENSMUSG00000032066 | 2.03611709 | Bco2 |
| ENSMUSG00000042784 | 2.03618733 | Muc1 |
| ENSMUSG00000074170 | 2.04740188 | Plekhf1 |
| ENSMUSG00000002885 | 2.05555763 | Adgre5 |
| ENSMUSG00000048240 | 2.06378702 | Gng7 |
| ENSMUSG00000053617 | 2.06949809 | Sh3pxd2a |
| ENSMUSG00000020623 | 2.07233534 | Map2k6 |
| ENSMUSG00000009418 | 2.09667204 | Nav1 |
| ENSMUSG00000019478 | 2.09939324 | Rab4a |
| ENSMUSG00000048498 | 2.10098379 | Cd300e |
| ENSMUSG00000026748 | 2.10986258 | Plxdc2 |
| ENSMUSG00000021508 | 2.1337765 | Cxcl14 |
| ENSMUSG00000079470 | 2.15811105 | Utp14b |
| ENSMUSG00000019987 | 2.17063097 | Arg1 |

**Supplementary Table 2**

| | | |
|---|---|---|
| ENSMUSG00000031958 | 2.17463596 | Ldhd |
| ENSMUSG00000041992 | 2.17489767 | Rapgef5 |
| ENSMUSG00000000489 | 2.17780823 | Pdgfb |
| ENSMUSG00000031891 | 2.1862198 | Hsd11b2 |
| ENSMUSG00000030468 | 2.19688566 | Siglecg |
| ENSMUSG00000027544 | 2.20713959 | Nfatc2 |
| ENSMUSG00000049848 | 2.21920775 | Ceacam19 |
| ENSMUSG00000036862 | 2.22084597 | Dchs1 |
| ENSMUSG00000036585 | 2.23606463 | Fgf1 |
| ENSMUSG00000018865 | 2.2635179 | Sult4a1 |
| ENSMUSG00000039239 | 2.26542185 | Tgfb2 |
| ENSMUSG00000032718 | 2.26667925 | Mansc1 |
| ENSMUSG00000032528 | 2.26677649 | Vipr1 |
| ENSMUSG00000059994 | 2.2797163 | Fcrl1 |
| ENSMUSG00000049866 | 2.30013156 | Arl4c |
| ENSMUSG00000021298 | 2.31021925 | Gpr132 |
| ENSMUSG00000015305 | 2.31880826 | Sash1 |
| ENSMUSG00000030246 | 2.31967201 | Ldhb |
| ENSMUSG00000039716 | 2.32494067 | Dock3 |
| ENSMUSG00000002985 | 2.33083713 | Apoe |
| ENSMUSG00000036718 | 2.3393814 | Micall2 |
| ENSMUSG00000027175 | 2.34407417 | Tcp11l1 |
| ENSMUSG00000085939 | 2.34427516 | Cd63-ps |
| ENSMUSG00000033788 | 2.34616961 | Dysf |
| ENSMUSG00000022123 | 2.34848933 | Scel |
| ENSMUSG00000027962 | 2.35296312 | Vcam1 |
| ENSMUSG00000010021 | 2.37357159 | Kif19a |
| ENSMUSG00000079227 | 2.38898936 | Ccr5 |
| ENSMUSG00000054951 | 2.39063747 | 9130008F23Rik |
| ENSMUSG00000035863 | 2.40803505 | Palm |
| ENSMUSG00000032122 | 2.43235084 | Slc37a2 |
| ENSMUSG00000034652 | 2.43889328 | Cd300a |
| ENSMUSG00000063506 | 2.45192004 | Arhgap22 |
| ENSMUSG00000000957 | 2.47303737 | Mmp14 |
| ENSMUSG00000022504 | 2.48413025 | Ciita |
| ENSMUSG00000024610 | 2.49825867 | Cd74 |
| ENSMUSG00000025351 | 2.50458998 | Cd63 |
| ENSMUSG00000106820 | 2.5283359 | D5Ertd605e |
| ENSMUSG00000038179 | 2.53248278 | Slamf7 |
| ENSMUSG00000030605 | 2.53401777 | Mfge8 |
| ENSMUSG00000045636 | 2.53448468 | Mtus1 |
| ENSMUSG00000053113 | 2.55251163 | Socs3 |
| ENSMUSG00000024381 | 2.55423056 | Bin1 |

### Supplementary Table 2

| | | |
|---|---|---|
| ENSMUSG00000026640 | 2.55461622 | Plxna2 |
| ENSMUSG00000041481 | 2.55859524 | Serpina3g |
| ENSMUSG00000044349 | 2.57535047 | Snhg11 |
| ENSMUSG00000040584 | 2.59051169 | Abcb1a |
| ENSMUSG00000086773 | 2.59472762 | Gm16192 |
| ENSMUSG00000032530 | 2.60229568 | Lyzl4 |
| ENSMUSG00000019558 | 2.60368817 | Slc6a8 |
| ENSMUSG00000073421 | 2.61075462 | H2-Ab1 |
| ENSMUSG00000039774 | 2.6291007 | Galnt12 |
| ENSMUSG00000062309 | 2.63900254 | Rpp25 |
| ENSMUSG00000091089 | 2.68240252 | NA |
| ENSMUSG00000022371 | 2.68586992 | Col14a1 |
| ENSMUSG00000055069 | 2.69490635 | Rab39 |
| ENSMUSG00000044206 | 2.70651976 | Vsig4 |
| ENSMUSG00000049709 | 2.70846061 | Nlrp10 |
| ENSMUSG00000036594 | 2.73641662 | H2-Aa |
| ENSMUSG00000050075 | 2.74320285 | Gpr171 |
| ENSMUSG00000034177 | 2.76605123 | Rnf43 |
| ENSMUSG00000006360 | 2.7799379 | Crip1 |
| ENSMUSG00000015647 | 2.79310594 | Lama5 |
| ENSMUSG00000053646 | 2.81276469 | Plxnb1 |
| ENSMUSG00000051379 | 2.82792292 | Flrt3 |
| ENSMUSG00000049723 | 2.83325767 | Mmp12 |
| ENSMUSG00000093327 | 2.83555783 | Mir5107 |
| ENSMUSG00000060586 | 2.86120033 | H2-Eb1 |
| ENSMUSG00000052974 | 2.88437053 | Cyp2f2 |
| ENSMUSG00000037362 | 2.90607073 | Nov |
| ENSMUSG00000023830 | 2.91153188 | Igf2r |
| ENSMUSG00000027955 | 2.91400926 | Fam198b |
| ENSMUSG00000005360 | 2.92171682 | Slc1a3 |
| ENSMUSG00000029765 | 2.97631874 | Plxna4 |
| ENSMUSG00000106062 | 2.98025631 | Gm43820 |
| ENSMUSG00000021719 | 3.00117934 | Rgs7bp |
| ENSMUSG00000028459 | 3.00368333 | Cd72 |
| ENSMUSG00000041797 | 3.02484138 | Abca9 |
| ENSMUSG00000026986 | 3.07134754 | Hnmt |
| ENSMUSG00000048442 | 3.07396179 | Smim5 |
| ENSMUSG00000087362 | 3.07542858 | Gm13710 |
| ENSMUSG00000040229 | 3.08416564 | Gpr34 |
| ENSMUSG00000025993 | 3.10986802 | Slc40a1 |
| ENSMUSG00000086763 | 3.1208957 | Plxna4os1 |
| ENSMUSG00000026437 | 3.15762971 | Cdk18 |
| ENSMUSG00000006445 | 3.18288027 | Epha2 |

### Supplementary Table 2

| | | |
|---|---|---|
| ENSMUSG00000100060 | 3.20136986 | Gm17944 |
| ENSMUSG00000047250 | 3.20231225 | Ptgs1 |
| ENSMUSG00000026365 | 3.27780705 | Cfh |
| ENSMUSG00000039232 | 3.28125034 | Stx11 |
| ENSMUSG00000045763 | 3.28571147 | Basp1 |
| ENSMUSG00000070407 | 3.2949122 | Hs3st3b1 |
| ENSMUSG00000025856 | 3.33278015 | Pdgfa |
| ENSMUSG00000008496 | 3.33680611 | Pou2f2 |
| ENSMUSG00000026938 | 3.36012386 | Fcna |
| ENSMUSG00000089991 | 3.36689683 | Gm16332 |
| ENSMUSG00000041552 | 3.41489907 | Ptchd1 |
| ENSMUSG00000039908 | 3.42090735 | Slc26a11 |
| ENSMUSG00000027347 | 3.45237966 | Rasgrp1 |
| ENSMUSG00000049103 | 3.47238916 | Ccr2 |
| ENSMUSG00000107622 | 3.50573765 | 4930512J16Rik |
| ENSMUSG00000001020 | 3.54008918 | S100a4 |
| ENSMUSG00000047180 | 3.57320597 | Neurl3 |
| ENSMUSG00000096630 | 3.58197246 | Vmn2r26 |
| ENSMUSG00000044485 | 3.6481691 | Klk1b11 |
| ENSMUSG00000057286 | 3.65698387 | St6galnac2 |
| ENSMUSG00000029096 | 3.74620288 | Htra3 |
| ENSMUSG00000031613 | 3.77958646 | Hpgd |
| ENSMUSG00000029299 | 3.80118505 | Abcg3 |
| ENSMUSG00000086712 | 3.8312582 | AI427809 |
| ENSMUSG00000035934 | 3.85122972 | Pknox2 |
| ENSMUSG00000031780 | 3.8659485 | Ccl17 |
| ENSMUSG00000047878 | 3.86649144 | A4galt |
| ENSMUSG00000025855 | 3.86659608 | Prkar1b |
| ENSMUSG00000032915 | 3.87092391 | Adgre4 |
| ENSMUSG00000007379 | 3.92406755 | Dennd2c |
| ENSMUSG00000045038 | 3.95073917 | Prkce |
| ENSMUSG00000026246 | 3.96072823 | Alppl2 |
| ENSMUSG00000028245 | 4.07735508 | Nsmaf |
| ENSMUSG00000005043 | 4.18069412 | Sgsh |
| ENSMUSG00000034416 | 4.18697596 | Pkd1l2 |
| ENSMUSG00000001025 | 4.24280149 | S100a6 |
| ENSMUSG00000027646 | 4.2770805 | Src |
| ENSMUSG00000028108 | 4.27874278 | Ecm1 |
| ENSMUSG00000022415 | 4.34950146 | Syngr1 |
| ENSMUSG00000038807 | 4.46022934 | Rap1gap2 |
| ENSMUSG00000033717 | 4.48605984 | Adra2a |
| ENSMUSG00000064057 | 4.78628858 | Scgb3a1 |
| ENSMUSG00000027485 | 4.79562629 | Bpifb1 |

**Supplementary Table 2**

| | | |
|---|---|---|
| ENSMUSG00000028970 | 4.95948473 | Abcb1b |
| ENSMUSG00000109713 | 5.08372663 | Pvrig |
| ENSMUSG00000044708 | 5.24687019 | Kcnj10 |
| ENSMUSG00000040552 | 5.55761185 | C3ar1 |
| ENSMUSG00000047592 | 5.58641528 | Nxpe5 |
| ENSMUSG00000025854 | 5.7367688 | Fam20c |
| ENSMUSG00000041272 | 5.80329289 | Tox |
| ENSMUSG00000045404 | 6.35043055 | Kcnk13 |
| ENSMUSG00000027483 | 6.5836229 | Bpifa1 |
| ENSMUSG00000042476 | 6.63077695 | Abcb4 |
| ENSMUSG00000022357 | 7.37547343 | Klhl38 |
| ENSMUSG00000042367 | 8.55820364 | Gjb3 |

**Supplementary Table 3**

| Process_name | Significant_genes_count | Total_genes _count | %_significant_genes | P-value | Padj-value |
|---|---|---|---|---|---|
| GO:0006935~chemotaxis | 18 | 119 | 15.12605042 | 3.81E-06 | 0.01100219 |
| GO:0060326~cell chemotaxis | 12 | 58 | 20.68965517 | 1.03E-05 | 0.01482853 |
| GO:0002376~immune system process | 33 | 384 | 8.59375 | 4.93E-05 | 0.03558469 |
| GO:0043931~ossification involved in bone maturation | 5 | 7 | 71.42857143 | 4.66E-05 | 0.03558469 |
| GO:0043406~positive regulation of MAP kinase activity | 10 | 51 | 19.60784314 | 8.29E-05 | 0.04786246 |

### Supplementary Table 4

| | Clone | Supplier | Cat. Number | RRID |
|---|---|---|---|---|
| **Flow cytometry** | | | | |
| ***Mouse*** | | | | |
| Rat monoclonal CD3 Biotin | 17A2 | Biolegend | 100244 | AB_2563947 |
| Rat monoclonal CD11a PE-Cy7 | I21/7 | Biolegend | 153108 | AB_2716057 |
| Rat monoclonal CD11b APC-Fire750 | M1/70 | Biolegend | 101262 | AB_2572122 |
| Rat monoclonal CD11b PE-Cy7 | M1/70 | Biolegend | 101216 | AB_312799 |
| Armenian hamster monoclonal CD11c BV785 | N418 | Biolegend | 117336 | AB_2565268 |
| Rat monoclonal CD19 Biotin | 6D5 | Biolegend | 115504 | AB_313639 |
| Rat monoclonal CD31 PE-Cy7 | 390 | Biolegend | 102418 | AB_830757 |
| Rat monoclonal CD31 BV421 | 390 | Biolegend | 102424 | AB_2650892 |
| Mouse monoclonal CD36 SuperBright 660 | HM36 | eBioscience | 63-0362-82 | AB_2734971 |
| Rat monoclonla CD45 BV510 | 30-F11 | Biolegend | 103138 | AB_2563061 |
| Mouse monoclonal CD45.1 FITC | A20 | Biolegend | 110706 | AB_313495 |
| Mouse monoclonal CD45.2 AF700 | 104 | Biolegend | 109822 | AB_493731 |
| Rat monoclonal CD63 PE | NVG-2 | Biolegend | 143904 | AB_11204430 |
| Mouse monoclonal CD64 PE-Cy7 | X54-5/7.1 | Biolegend | 139314 | AB_2563904 |
| Mouse monoclonal CD64 APC | X54-5/7.1 | Biolegend | 139306 | AB_11219391 |
| Mouse monoclonal CD64 BV421 | X54-5/7.1 | Biolegend | 139309 | AB_2562694 |
| Rat monoclonal CD90.2 FITC | 30-H12 | Biolegend | 105305 | AB_313176 |
| Rat monoclonal CD102 AF647 | 3C4 | Biolegend | 105612 | AB_2122182 |
| Rat monoclonal CD103 AF488 | 2E7 | Biolegend | 121408 | AB_535950 |
| Rat monoclonal CD103 AF647 | 2E7 | Biolegend | 121410 | AB_535952 |
| Rat monoclonal CD106 (VCAM1) AF647 | 429 | Biolegend | 105711 | AB_493430 |
| Rat monoclonal CD115 APC | AFS98 | Biolegend | 135510 | AB_2085221 |
| Rat monoclonal CD140a (PDGFRa) PE | APA5 | BD Bioseince | 526776 | AB_2737787 |
| Rat monoclonal CD146 APC | ME-9F1 | Biolegend | 134712 | AB_2563088 |
| Rat monoclonal CD192 (CCR2) PE | SA203G11 | Biolegend | 150610 | AB_2616982 |
| Rat monoclonal CD326 (EpCAM) BV605 | G8.8 | Biolegend | 118227 | AB_2563984 |
| Rat monoclonal CD326 (EpCAM) PE | G8.8 | Biolegend | 118206 | AB_1134172 |
| Rat monoclonal CD354 (TREM1) eF660 | TR3MBL1 | eBioscience | 50-3541-82 | AB_2574205 |
| Mouse monoclonal C/EBPβ AF647 | H7 | Santa Cruz Biotech | Sc-7962 | AB_626772 |
| Mouse monoclonal CX3CR1 AF647 | SA011F11 | Biolegend | 149004 | AB_2564273 |
| Rat monoclonal EGR2 PE | erongr2 | eBioscience | 12-6691-82 | AB_10717804 |
| Rat monoclonal EGR2 APC | erongr2 | eBioscience | 17-6691-82 | AB_11151502 |
| Rat monoclonal F4/80 PE | BM8 | Biolegend | 123110 | AB_893486 |
| Rat monoclonal GP38 PE-Cy7 | 8.1.1 | Biolegend | 127412 | AB_10613648 |
| Recombinant Ki67 FITC | REA183 | Miltenyi Biotec | 130-117-691 | AB_2733585 |
| Rat monoclonal Ly6C eFluor450 | HK1.4 | eBioscience | 48-5932-82 | AB_10805519 |
| Rat monoclonal Ly6C PerCP-Cy5.5 | HK1.4 | Biolegend | 128012 | AB_1659241 |
| Rat monoclonal Ly6G Biotin | 1A8 | Biolegend | 127604 | AB_1186108 |
| Rat monoclonal Lyve1 eFluor 660 | ALY7 | eBioscience | 50-0443-82 | AB_10597449 |
| Rat monoclonal MerTK PE | 2B10C42 | Biolegend | 151506 | AB_2617037 |
| Rat monoclonal MHCII (IA-IE) AF700 | M5/114.15.2 | Biolegend | 107622 | AB_493727 |
| Mouse monoclonal NK1.1 Biotin | PK136 | Biolegend | 108704 | AB_313391 |
| Rat monoclonal SiglecF PE-CF594 | E50-2440 | BD Bioscience | 562757 | AB_2687994 |
| Rat monoclonal Siglec H PE | 551 | Biolegend | 129605 | AB_1227763 |
| ***Human*** | | | | |
| Rabbit polyclonal EGR2 (unconjugated) | | Invitrogen | PA565091 | AB_2662529 |
| Mouse monoclonal HLA-DR eFluor450 | LN3 | eBioscience | 48-9956-42 | AB_10718248 |
| Mouse monoclonal CD3 FITC | UCHT1 | Biolegend | 300452 | AB_2564148 |
| Mouse monoclonal CD19 FITC | HIB19 | Biolegend | 302206 | AB_314236 |
| Mouse monoclonal CD56 FITC | 5.1H11 | Biolegend | 362546 | AB_2565964 |
| Mouse monoclonal CD66b FITC | G10F5 | Biolegend | 305104 | AB_314496 |
| Mouse monoclonal CD163 APC | RM3/1 | Biolegend | 326510 | AB_2564015 |
| Rat monoclonal CD11b PE-Cy7 | M1/70 | Biolegend | 101216 | AB_312799 |
| ***Other flow cytometry reagents*** | | | | |
| Streptavidin BV650 | N/A | Biolegend | 405232 | |
| 7-AAD | | Biolegend | 420404 | |
| Zombie NIR Fiable Viability Dye | | Biolegend | 423106 | |
| **Immunofluorescence imaging** | | | | |
| Rabbit polyclonal CD68 (unconjugated) | N/A | Abcam | Ab125212 | AB_10975465 |
| Mouse monoclonal aSMA Cy3 | 1A4 | Merck (Sigma) | C6198 | AB_476856 |
| Donkey anti-rabbit AF488 | N/A | ThermoFisher | A-21206 | AB_2535792 |
